# AI-guided analysis of human pancreatic islet sociology reveals distinct cell compositional changes in type 1 diabetes

**DOI:** 10.64898/2026.06.05.730081

**Authors:** Caroline Ward, Tabitha Banks-Tibbs, Henry Thorpe, Antonia M. Giles, Samuel J. Mabry, Jia-Jun Liu, Hung-Ching Chang, Jinting Yang, Aidan F. Carney, Halla M. Shaikh, Lauren M. Woolley, Paul N. Joseph, Jenesis Kozel, Emily M. Rocha, Guy A. Rutter, George C. Tseng, Silvia Liu, Lora L. Pless, Zachary Freyberg

**Author notes:** **Corresponding Author:** Zachary Freyberg M.D., Ph.D. 203 Lothrop Street, Eye and Ear Institute, Room 115 Pittsburgh, PA 15213, USA. These authors contributed equally (Caroline Ward, Tabitha Banks-Tibbs, Henry Thorpe).

## Abstract

Human pancreatic islets exhibit greater anatomic and cellular heterogeneity than previously appreciated, raising fundamental questions about how their composition varies with age, sex, region, and islet size and how type 1 diabetes (T1D) alters these relationships. Yet these questions remained largely unresolved due to the bottleneck of manual tissue inspection. Here, we developed an integrated artificial intelligence (AI)-guided imaging, processing, and statistical pipeline enabling unbiased, high-throughput analysis of more than 2 million candidate islets from 106 non-diabetic (ND) and T1D donors. We identified age-, region-, sex-, and islet size-dependent differences in islet distribution and composition between ND and T1D donors. Profound β-cell loss in T1D was accompanied by reciprocal α-cell expansion, whereas δ-cells and pancreatic polypeptide cells were largely resilient. Cell area and pseudotime analyses uncovered regional and age-dependent trajectories of islet remodeling across T1D progression, along with distinct patterns of cytoarchitectural reorganization of the endocrine pancreas.

## Introduction

The endocrine pancreas is composed of islets of Langerhans, which contain four major hormone-producing cell types: insulin-secreting β-cells, glucagon-secreting α-cells, somatostatin-secreting δ-cells, and pancreatic polypeptide (PP)-cells^1^. These distinct cell populations do not function in isolation but instead rely upon ongoing communication via autocrine and paracrine signaling to coordinate regulated hormone secretion^2–5^. Conversely, the loss of local intra-islet communication leads to dysglycemia and is increasingly linked to the development of diabetes^2,6^. Type 2 diabetes (T2D), the most prevalent form of diabetes, represents ∼95% of cases with heterogeneous pathophysiology ^6,7^. Type 1 diabetes (T1D), which constitutes a minority of diabetes cases, is also a complex multifactorial illness caused by autoimmune destruction of β-cells^8^. Nevertheless, T1D prevalence continues to steadily rise in pediatric populations^9,10^, creating a strong rationale for better understanding T1D pathophysiology. Yet despite decades of investigation, fundamental questions remain about how T1D reshapes the full complement of islet cell types, and how anatomic, demographic, and architectural factors shape this response.

Foundational histopathological studies established that β-cell loss in T1D is profoundly heterogeneous across patient, islet, and cellular levels. Early quantitative and immunocytochemical analyses demonstrated near-complete β-cell depletion in most long-duration cases, while a minority of patients retained residual β-cells at reduced but variable levels^11,12^. Meier and colleagues subsequently reported a ∼40-fold reduction in β-cell area in long-standing T1D compared to non-diabetic (ND) subjects, with substantial between-patient variation^13^. Yet residual β-cell content can persist decades into disease. Indeed, in “Joslin Medalists” with more than 50 years of T1D, pancreata retained detectable insulin-positive cells; in a subset of patients, particularly those with later disease onset, 10-20% of normal β-cell content was preserved^14^. This heterogeneity also extends to the cellular level where individual β-cells differ in their vulnerability to autoimmune destruction^15–18^. Growing evidence indicates that a notable degree of T1D-associated β-cell loss may reflect dedifferentiation and transdifferentiation rather than cellular destruction^19–22^. Despite consistently revealing this multi-scale heterogeneity, however, earlier studies were constrained by the bottleneck of manual histology, which limited them to modest donor cohorts and small numbers of manually inspected islets per case. The resulting designs were underpowered in part by demographic and clinical determinants of β-cell variability. Collectively, these studies established that β-cell loss in T1D is non-uniform but left the sources of that non-uniformity largely unresolved. As a result, this has motivated the development of higher-throughput approaches capable of scaling to islet-level resolution and with larger cohort sizes required to map islet and disease heterogeneity systematically.

A new wave of imaging and analytical methods has begun to overcome prior limitations in studying T1D islet biology. A central theme emerging from this work is that islet vulnerability to T1D is contingent on the size and composition of endocrine objects (EOs). Recent imaging studies of intact human pancreata revealed that islet distribution is non-uniform across the head, body, and tail regions^23–27^. Such heterogeneity is also increasingly recognized at the EO level. For example, among ND donors, substantial numbers of EOs are largely composed of β-cells and lacking α-cells, often found as small, extra-islet objects which were previously largely overlooked^28^. Subsequent studies in the disease context revealed that these small, β-cell-rich populations are particularly vulnerable in T1D^29^. Complementary artificial intelligence (AI)-assisted two-dimensional (2D) whole-slide analysis confirmed that small extra-islet β-cell clusters are virtually absent in T1D pancreata, most strikingly in donors diagnosed in early childhood^30^. Higher-dimensional multiplexed immunohistochemistry has extended these findings, identifying histopathological correlates of T1D pathology already evident at preclinical stages of the illness^31^. Collectively, these advances establish that T1D-induced islet remodeling is anatomically heterogeneous, size-dependent, and cell-type-specific.

To date, several critical knowledge gaps remain. The majority of work has thus far focused on β-cells alone, or on β- and α-cells in tandem. Instead, comprehensive quantification across all four major endocrine cell types remains rare. Consequently, whether δ-cells and PP-cells are resilient, vulnerable, or actively remodeled in T1D is largely unresolved. Moreover, many fundamental donor variables such as donor age, sex, pancreatic region, disease duration, and islet size have not been systematically interrogated. Therefore, it remains unclear if and/or how these patient characteristics contribute to T1D’s effects on islet composition. These limitations are in large part because of low sample size due to the constraints associated with low-throughput manual counts. On the other hand, with the emergence of machine learning approaches which enable rapid analysis of thousands of islets nested within donors, it has become challenging to rigorously accommodate the hierarchical nature of these much larger datasets. This includes handling the bounded distribution of compositional proportions within islets, or the high-dimensional covariate interactions through which T1D pathology may be mediated. Finally, many open questions remain concerning whether trajectories of compositional remodeling vary systematically across pancreatic region, sex, and age. Effectively resolving these questions requires substantially larger cohorts and islet-level scale compared to most earlier methods.

In addressing these gaps, here we have built an integrated platform combining AI-guided whole-slide image analysis with a tailored statistical framework. Applied to immunohistochemically stained pancreatic tissue from 106 ND and T1D donors, our AI pipeline simultaneously quantified all four major islet endocrine cell populations (β, α, δ, PP) across more than two million candidate islets. The large volume and hierarchical structure of these data exceeded what standard statistical approaches could accommodate. We therefore developed a dedicated analytical framework built on mixed-effects regression models with donor-level random effects, each matched to the natural distribution of its outcome (bounded proportions, gamma-distributed densities, counts, and areas), and capable of rigorously testing three- and four-way interactions among diagnosis, pancreas region, sex, age, and EO size. This analytical framework revealed age-, region-, sex-, and islet-size-specific differences in islet distribution and composition between ND and T1D donors. Profound β-cell loss in T1D was accompanied by reciprocal α-cell expansion, whereas δ-cells and PP-cells proved largely resilient. These findings therefore demonstrate a divergence in cell-type vulnerability to T1D. Per-cell area analyses further dissociated changes in cell number from changes in cell size, exposing distinct cell type-specific signatures to islet remodeling. Additionally, pseudotime trajectory inference identified region- and age-dependent paths of compositional remodeling across T1D progression, with the pancreatic head and younger donors exhibiting the most distinct trajectories.

## Materials and Methods

### Donor cohorts

Human postmortem pancreata were recovered from cadaveric organ donors and processed by nPOD as previously described^32–35^. All procedures were in accordance with federal and institutional guidelines and approved by the University of Florida Institutional Review Board. Our original dataset included a total of 106 donors (71 ND, 35 T1D) who contributed 2,696 pancreatic tissue images for subsequent analyses. Islet density was calculated from this full cohort of 106 donors (71 ND, 35 T1D) consisting of either single-, double-, or quadruple-stained images from each donor. For islet composition analyses, we examined a subset of the samples that were quadruple-stained via immunohistochemistry (IHC) for β-cells, α-cells, δ -cells, and PP-cells; this quadruple-stained subcohort consisted of 93 donors (59 ND, 34 T1D) who contributed 835 pancreatic tissue images. Inclusion criteria for the study included availability of high-quality digital scans of IHC-stained pancreatic sections. Exclusion criteria included the presence of a type 2 diabetes (T2D) diagnosis, autoantibody positivity in the absence of a T1D diagnosis, and donors >40 years of age. Demographic information from the subjects within the cohort was obtained from the nPOD database (see **Table 1**, **Supplementary Table S1** for subject demographic information).

**Table 1.** Subject demographics. Donor numbers (N), Age (years), sex, race/ethnicity, hemoglobin A1c (HbA1c, %), and body mass index (BMI, kg/m^2^). Age, HbA1c, and BMI are presented as mean ± SD.

| **Diagnosis** | **ND** | **T1D** |
| --- | --- | --- |
| **N** | 71 | 35 |
| **Age** | 17.26 ± 11.33 | 20.83 ± 8.35 |
| **Sex** |  |  |
| Female | 28 | 17 |
| Male | 43 | 18 |
| **Race/Ethnicity** |  |  |
| African American | 13 | 8 |
| Native American | 1 | 0 |
| Asian | 1 | 0 |
| Caucasian | 46 | 24 |
| Hispanic | 9 | 3 |
| Multiracial | 1 | 0 |
| **HbA1c** | 5.42 ± 0.49 | 10.24 ± 2.54 |
| **BMI** | 22.28 ± 5.29 | 25.75 ± 5.61 |

### Tissue histology, immunohistochemistry, and imaging

Pancreatic samples were divided into three anatomic regions whenever possible: pancreatic head, body, and tail. Cross sections (approximately 5 mm thick) were taken from each region and processed into paraffin and OCT blocks as previously described^32,34,35^. Briefly, 4 µm-thick formalin-fixed paraffin-embedded (FFPE) or 5 µm-thick OCT sections from each respective pancreatic region were stained with hematoxylin and eosin for tissue histology. For immunohistochemical (IHC) analysis, heat-induced epitope retrieval was performed using Borg Decloaker (Biocare Medical, Pacheco, CA) according to the manufacturer’s instructions prior to staining with two antibody panels as previously described^32,35,36^. Panel one included a triple stain with rabbit monoclonal anti-insulin (clone EPR17359, 1:2000; RRID:AB_2716761; Abcam, Waltham, MA, USA), mouse monoclonal anti-glucagon (clone K79bB10, 1:1000; RRID:AB_297642; Abcam), rabbit anti-Ki67 (clone EPR3610, diluted 1:1000 RRID:AB_10562976; Abcam), or rabbit anti-CD3 (1:100; RRID:AB_2335677; Agilent Technologies, Inc., Santa Clara, CA, USA); Ki67 and CD3 data were excluded from our analyses. Panel two included a quadruple stain with rabbit anti-insulin (clone EPR17359, 1:2000; RRID:AB_2716761; Abcam), mouse anti-glucagon (clone K79bB10, 1:1000; RRID:AB_297642; Abcam), rabbit polyclonal anti-somatostatin (1:1000; RRID:AB_572264; Agilent Technologies), and mouse monoclonal anti-pancreatic polypeptide (clone MM0858-31R25, 1:750; RRID:AB_2904511; Abcam). The respective antibody labeling was detected via chromogenic IHC staining using the Mach2 Double Stain1/Mach2 Double Stain 2 HRP-AP Polymer Detection Kit according to the manufacturer’s instructions (Biocare Medical). Slides were counterstained with hematoxylin and digitally imaged using an Aperio CS2 slide scanner (Leica Biosystems, Inc., Wetzlar, Germany) at 20x magnification.

### Pancreatic tissue analyses

**Tissue type analysis.** Imaged pancreatic tissue data was segmented and analyzed via the HALO image analysis platform (v3.5.3577.318; Indica Labs, Albuquerque, NM). The data were initially processed through a Tissue Classifier module to accurately detect pancreatic islets within the imaged tissue. Separate classifier modules were constructed for each donor to account for potential differences in marker positivity, label intensity, or tissue features between individuals. This included the generation of separate modules for different IHC stain combinations, *i.e*., doubled-stained (insulin, glucagon) versus quadruple-stained tissue (insulin, glucagon, somatostatin, PP). Tissue folds or bubbles associated with tissue staining, which could potentially distort imaging analyses were manually excluded from analysis using HALO’s annotation tool. The accuracy of classification was confirmed through expert pathological review. The Tissue Classifier was also implemented to calculate the proportional area of each pancreatic tissue type (tissue type divided by tissue area) per total tissue area as described previously^32^.

**Cell type analysis.** To identify and quantify different islet cell populations, we employed two AI-based classifier modules, Nuclei Seg and Nuclei Phenotyper, which use iterative machine learning approaches to identify individual islet cell nuclei. We first used Nuclei Seg for accurate nuclear segmentation of hematoxylin-stained nuclei. Though the module was pre-trained from various hematoxylin & eosin (H&E)-stained and 3,3’-diaminobenzidine (DAB)-stained IHC images to detect “Nuclei” versus “Background” to improve nuclei disclosure, we provided additional training to boost the accuracy of nuclear segmentation within our dataset. We trained the Nuclei Seg module with 8,793 manually annotated nuclei along with 128 non-nuclear “background” annotations across 11 randomly selected images from 8 donors (6423, 6562, 6547, 6557, HPAP048, HPAP068, HPAP078).

We then utilized the Nuclei Phenotyper to assign identified nuclei according to their respective cells based on cell type-specific cytoplasmic chromogenic staining of four distinct islet cell populations: β-cells, α-cells, δ-cells, and PP-cells within islets along with the exocrine acinar cells adjacent to the islets. The Nuclei Phenotyper module was initially trained using 5 ND donor images containing a total of 4,565 manually annotated cells: 1,390 β-cells, 776 α-cells, 349 δ-cells, 206 PP-cells and 1,844 acinar cells; AI training terminated after 930 training iterations (cross-entropy: 0.186). For analysis of T1D donor images, our newly trained Nuclei Phenotyper module required additional training to improve accurate cell type identification. Building on the Nuclei Phenotyper’s initial ND donor tissue training, the module was further trained on 3 T1D donor images containing a total of 2,290 manually annotated islet cells: 319 β-cells, 986 α-cells, 187 δ-cells, 86 PP-cells, and 721 acinar cells; AI training stopped after 225 training iterations (cross-entropy: 0.206).

Finally, with our newly trained AI modules we generated a complete image analysis workflow which incorporates: 1) the trained Tissue Classifier module to distinguish endocrine islets from local exocrine acinar cells within donor pancreatic tissue; followed by 2) the combination of our trained Nuclei Seg and Nuclei Phenotyper modules with the Multiplex IHC module to accurately quantify the constituent cell types within the identified islets.

**AI classifier accuracy assessment.** To evaluate the accuracy of the trained AI Nuclei Seg and Nuclei Phenotyper classifiers, we compared AI-generated counts of the respective islet cell populations to our ground truth: manually annotated quadruple-stained islets that were randomly selected from our ND and T1D datasets. To generate the ground truth, we manually annotated and counted IHC-labeled cells for each of the four islet cell populations (β-cells, α-cells, δ-cells, PP-cells). To quantify these comparisons, we calculated the coefficient of determination (r^2^) and root mean squared error (RMSE) between the AI-guided versus manually counted data via linear regression plots; separate comparisons were made for the ND and T1D groups. The AI classifiers were trained until the r^2^ values stopped improving, reaching a cutoff range of r^2^=∼0.7-0.9 to ensure sufficient confidence in the accuracy of the subsequent unsupervised recognition of the different islet cell types.

### Data Processing

Initial processing. All HALO data were exported as .csv files according to image. Additional information needed for analyses (*i.e.*, pancreatic region, IHC stain, age, and donor identifier) were added to datasets. Regarding age, all analyzed data came from subjects ≤40 years. We established a cutoff range of ≥15 and ≤244 cells/islet to be considered an islet via comprehensive expert review of the imaging data and mapping the distribution of cells/islet across all donors (Supplementary Figure S4). Additionally, this number was validated based on recent estimates of human islet composition, including in the context of T1D^37,38^, enabling us to obtain more accurate representations of islet composition within the respective groups. We also examined single β-cells, α-cells, δ-cells, and PP-cells as single EOs. Small endocrine objects (SEOs) were defined as 2-14 cells.

Following our initial processing, based on the above cutoff criteria, 106 donors remained for the islet density analysis. From this initial set of 106 donors, we used a subset of 93 donors for analyses of islet cell composition and cell counts; donors 6112, 6117, 6122, 6250, 6294, 6305, 6309, 6313, 6348, 6356, 6357, 6366, 6432 were excluded since the images from these subjects did not have all four cell types labeled, *i.e*., only single- or double-staining. For islet β-cell/(α-cell + β-cell) ratio analyses, we used a subset of 95 donors based on double- and quadruple-stained images; donors 6112, 6117, 6122, 6294, 6305, 6309, 6313, 6348, 6356, 6366, 6473 were excluded on the basis of single staining only and/or poor image quality. For islet cell area analyses, we employed a further subset of donors; from the 93 donors above, we excluded one additional donor 6547 on the grounds of technical mismatch to rigorously couple area data with cell count data.

Islet density analyses. Islet density was computed as the number of islets identified on an imaged slide divided by the effective tissue area (islets/mm^2^). Tissue area was quantified according to the equation: *Tissue area analyzed = Classified area − Duct area − Glass area*. Islets were previously identified using the Tissue Classifier module.

Islet cell composition. Islet cell composition was assessed from our AI classifier outputs to calculate: 1) the islet β-cell/(α-cell + β-cell) ratio, and 2) the relative proportions of different islet cell types including β-cells, α-cells, δ-cells, and PP-cells per islet (expressed as percentages of the total islet cell number). Islet β-cell/(α-cell + β-cell) ratios were calculated from both double-stained pancreatic tissue images (featuring IHC-labeled β-cells, α-cells) and quadruple-stained images (featuring IHC-labeled β-cells, α-cells, δ-cells, PP-cells). Islet cell type proportions were calculated according to the respective percentages of each of the four labeled cell types per islet; we divided total numbers of each cell type (numerator) by the total number of cells within each islet (denominator). Finally, given our specific focus on endocrine islets and their constituent cell types, we did not include non-endocrine acinar cells potentially recognized in our images for islet composition analyses.

**Cell type area and per-cell area.** Cell type area was calculated via the Area Quantification Module in HALO. We analyzed slides single-labeled for β-cells, double-labeled for β-cell and α-cells, as well as quadruple-labeled for β-cells, α-cells, δ-cells, and PP-cells. Cell type areas were calculated for single EOs (*i.e*., single cells), SEOs, and islets. To calculate area per cell type, cell type area outputs were divided by the respective cell type counts for each EO.

### Statistical analyses

**Model selection.** We used a generalized linear mixed model (GLMM) to evaluate islet cell type data, differences according to available covariates (*i.e.,* sex, age, pancreatic region, diagnosis) as well as to discern potential covariate interactions. We accounted for donor as a random variable in models built using the ‘glmmTMB’ R package (version 1.1.9)^39^. We also used the ‘ordbetareg’ R package (version 0.8)^40^. Our choice of model was guided by the substantial numbers of islets and constituent islet cell types per subject generated by our AI-guided approaches. A subset of the data was transformed for islet β-cell/(α-cell + β-cell) ratio measurements as indicated by the formula. Models for all outcomes were selected based on outcome distribution and model metrics including Akaike Information Criterion (AIC) and Bayesian Information Criterion (BIC).

**Distribution estimation and selection.** To estimate the distribution of data, we generated histograms via the ‘fitdistrplus’ R package (version 1.2-2)^41^; Cullen and Frey graphs were also generated using ‘fitdistrplus’ to check the outcome distributions. In the case that multiple distributions were candidates for subsequent modeling, a goodness of fit comparison was performed using the ‘gofstat’ function within the ‘fitdistrplus’ package to identify the optimal distribution.

We next chose the most appropriate distribution of outcomes using Kolmogorov-Smirnov, Cramer-von Mises, and Anderson-Darling statistics. Based on these analyses, we chose a gamma distribution for islet density analyses and a beta distribution for islet cell composition, and β-cell/(α-cell + β-cell) ratio analyses. We employed a lognormal distribution for islet cell area analyses. A negative binomial distribution was used for cell count data.

**Table generation.** Tables were created for each set of interactions tested within each model. These tables describe the number of subjects who contributed to every category of each interaction and the total samples. Separate GLMMs were then created for islet density, each of the four islet cell types, for β-cell/(α-cell + β-cell) ratio analyses, islet cell areas, and islet cell counts. The models were created in the ‘glmmTMB’ (version 1.1.9) or ‘ordbetareg’ (version 0.8) R packages.

**Model validation.** We validated our models by calculating variance within the data (see Table S2). These model metrics included the calculation of conditional and marginal r^2^ values for the full model via the r.squaredGLMM() function within the MuMIn R package (version 1.48.19)^42^. We also calculated the r^2^ values for reduced models to find the f^2^ values for each of our covariates. Additionally, we employed the standardize_parameters() function as another metric for variable importance.

**Post-hoc Analyses.** To assess differences between subgroups, Tukey post-hoc analyses were performed on each model using the ‘emmeans’ R package (version 1.10.4)^43^. Statistical significance was determined according to posterior probabilities, Tukey-adjusted p-values, and z-scores.

**Data Visualization.** Graphs related to AI validation were generated using GraphPad Prism (version 10.4.0; GraphPad Software, San Diego, CA). All other graphs were generated using the ‘ggplot2’ R package (version 3.5.1)^44^. To ensure consistent representation across samples with varying numbers of images, we selected 100 images per donor, corresponding to each percentile of cell proportion, and used a heatmap to present the distribution of cell type proportions for each donor. This down-sampling strategy in hierarchical data enabled us to avoid over-emphasizing donors with larger numbers of images and under-representing donors with fewer images. Split violin plots were also generated via ‘ggplot2’ to compare the distribution of cell proportions between T1D and ND donors for each islet cell type. For each significant interaction within regression analyses, we created graphs using ‘ggplot2’ to visualize potential changes in cell type composition according to the different covariates analyzed.

**Pseudotime trajectory inference.** Dimension reduction of the islet cell composition data was first performed based on the proportions of β-cells, α-cells, δ-cells, and PP-cells followed by visualization using Uniform Manifold Approximation and Projection (UMAP)^45^. Islets were further categorized according to diagnosis (ND, T1D), region (head, body, tail), and age (<20-year-old, 20-40-year-old). Among each islet covariate subcategory, trajectory inference was performed using the Monocle3 software package^46,47^. Trajectory graphs and pseudotimes were calculated and mapped onto the respective UMAPs to display the impact of T1D progression on changes in islet cell composition. Wilcoxon rank-sum tests were performed to compare the pseudotime distributions between ND and T1D donors.

## Data Availability

Donor pancreatic image data is available through nPOD program. All custom-written code will be publicly available on Github (https://github.com/FreybergLab/nPOD-Project). Any intermediate files not available here can be requested from the corresponding author.

## Results

### Development of an AI-guided workflow for image analysis of human pancreatic tissue from ND and T1D donors

To enable unbiased, high-throughput interrogation of islet distribution and cellular composition in ND and T1D states, we established an integrated end-to-end pipeline that couples AI-guided whole-slide image analysis to a tailored data-processing and statistical workflow (**Figure 1**). High-resolution images of immunohistochemically (IHC)-stained pancreatic sections from 106 male and female donors (71 ND, 35 T1D) across a range of ages (see **Table 1**, **Supplementary Table S1** for donor demographics) were processed through a sequence of HALO classifier modules: (i) a per-donor Tissue Classifier was trained to identify islets and to exclude non-endocrine (*e.g*., exocrine ducts, acinar cells, fibrotic, adipose, and connective tissues), and artifactual features (*e.g*., air bubbles) [**Figure 1A(i-ii)**]. The module employs a random forest machine learning algorithm, enabling it to categorize different tissue types based on color, texture, and contextual features according to a learn-by-example tactic. (ii) a Nuclei Seg module, built on a ResNet-34 convolutional neural network (CNN) with watershed post-processing, was used to segment hematoxylin-stained nuclei within identified islets^48^ [**Figure 1A(iii)**]; and (iii) a Nuclei Phenotyper module, based on a ResNet-18 deep CNN^48^, assigned segmented nuclei to one of four endocrine cell populations on the basis of cytoplasmic chromogenic staining for insulin (β-cells), glucagon (α-cells), somatostatin (δ-cells), and pancreatic polypeptide (PP-cells) [**Figure 1A(iv-v)**]. Applied across the full image set, this pipeline identified an initial total of 2,081,140 islet candidates, with each constituent nucleus assigned to a defined cell type (**Supplementary Figure S1**).

**Figure 1.**
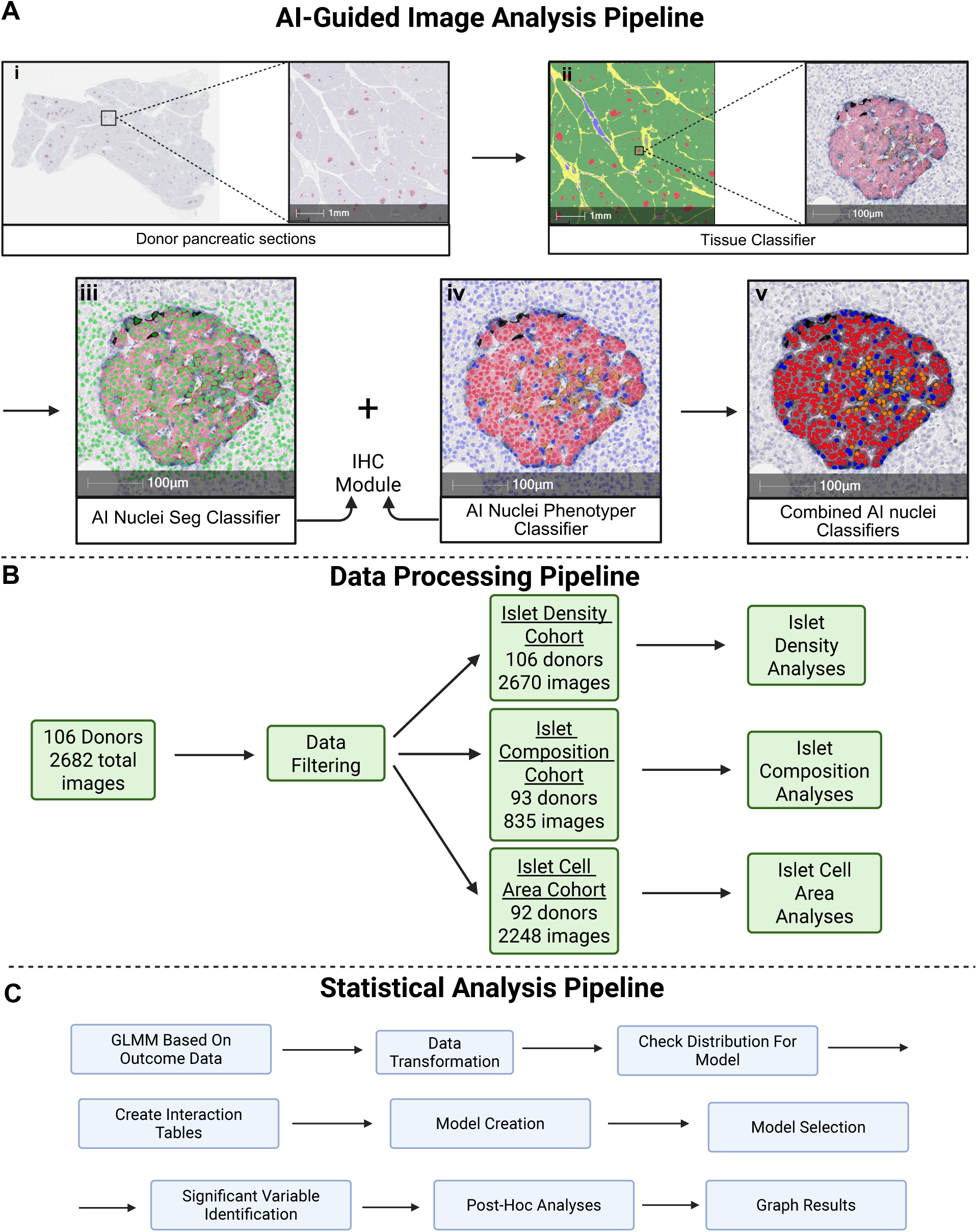
Overview of integrated pipelines for AI-guided image analysis of human pancreatic tissue, data processing, and statistical analyses. (A) Schematic of AI-guided analysis of high-resolution whole-slide images of human pancreatic tissue. Inset: magnified region highlights tissue islet distribution; scale bar=1 mm (i). An AI-guided Tissue Classifier module was established for each donor to identify islets (in red); scale bar=1 mm. Inset: representative image of an islet recognized by the classifier; scale bar=100 µm (ii). Manual segmentation of hematoxylin-stained islet cell nuclei (indicated by open green circles) trained the AI-guided Nuclei Seg classifier for accurate segmentation of nuclei within islets; scale bar=100 µm (iii). A trained AI-based Nuclei Phenotyper module assigned identified nuclei to distinct islet cell populations; scale bar=100 µm (iv). The Nuclei Seg and Nuclei Phenotyper modules were combined within the IHC module to analyze islet area along with cell type counts and densities; scale bar=100 µm. (v). (B) Flowchart of the data processing pipeline. Image analysis data from 106 donors (2696 images) were filtered according to donor age and cutoff criteria for islet cell numbers. This resulted in final cohorts of 106 donors for islet density analyses, 93 donors for islet composition analyses, and 92 donors for islet cell area analyses (see Methods). (C) Flowchart of the statistical analysis pipeline. Our approach employed a generalized linear mixed models (GLMM) to statistically analyze islet data according to covariates (*i.e.,* diagnosis, disease duration, sex, age, pancreatic region) and their potential interactions.

### Training improves the annotation accuracy of AI-guided analyses of islet cell composition

We next assessed the accuracy of the trained AI classifiers in ND donor tissue against expert-curated manual annotations as ground truth (**Supplementary Figure S2**). For total cells per islet (all cell types combined), training improved overall annotation accuracy, lowering RMSE 1.7-fold (untrained: 172.16; trained: 100.03) with a higher coefficient of determination (untrained r^2^ = 0.80; trained r^2^ = 0.84) (**Supplementary Figure S2A**). Per-cell-type accuracy improved further: for β-cells, the r^2^ rose 1.1-fold (untrained: 0.76; trained: 0.85) and RMSE fell 2.1-fold (RMSE untrained: 132.20, trained: 61.75; **Supplementary Figure S2B**), and for α-cells RMSE diminished 5.8-fold (untrained: 307.00; trained: 52.85) while r^2^ values remained relatively stable (untrained: 0.70; trained: 0.64) (**Supplementary Figure S2C**). The largest gains were observed for δ-cells and PP-cells, the two sparsest islet populations. AI classifier training boosted δ-cell r^2^ 1.4-fold (untrained: 0.52; trained: 0.75) while RMSE was reduced from 16.86 to 12.08 (**Supplementary Figure S2D**). For PP-cells, classifier training resulted in a 10.7-fold increase in r^2^ (untrained: 0.07; trained: 0.75) and a 7.4-fold reduction in RMSE (untrained: 57.99; trained: 7.87) (**Supplementary Figure S2E**).

Applying the same approach to T1D donor tissue produced similar accuracy gains (**Supplementary Figure S3**). For total cell counts, RMSE dropped 3.2-fold (untrained: 292.40; trained: 90.89) with r^2^ values already high prior to additional training (untrained: 0.95; trained: 0.93) (**Supplementary Figure S3A**). β-cell and α-cell annotations again improved markedly. For β-cells r^2^ values improved 1.2-fold (untrained: 0.81; trained, 0.93) with RMSE dropping 3.1-fold (untrained: 137.20; trained: 43.92). For T1D donors, α-cell r^2^ values showed similar improvements (untrained: 0.80; trained: 0.92), together with RMSE (untrained: 470.50; trained: 90.44) (**Supplementary Figure S3B-C**). The greatest improvements were found in δ-cells with a 20.8-fold improvement in r^2^ values (untrained: 0.04; trained: 0.83) and a 2.9-fold drop in RMSE (untrained: 51.70; trained: 17.57) (**Supplementary Figure S3D**), as well as in PP-cells with a 5.1-fold rise in r^2^ values (untrained: 0.19; trained: 0.97) and 5.8-fold decrease in RMSE (untrained: 67.45; trained: 11.54) (**Supplementary Figure S3E**). Together, these data demonstrate that, with appropriate training, our AI-guided workflow can accurately recapitulate manual annotations for all four major islet endocrine cell populations across both ND and T1D donor tissue.

### Development of islet data-processing and statistical pipelines

To support reproducible, high-throughput downstream analysis, AI classifier outputs were channeled into a tailored data-processing pipeline (**Figure 1B**). Per-image data were merged with donor metadata (diagnosis, sex, age, pancreatic region, disease duration) and filtered according to age (≤40 years) and per-object cell counts. A 15-cell threshold was applied to distinguish islets (≥15 cells per object) from single endocrine cells and small endocrine objects (SEOs; 2-14 cells per object) (**Supplementary Figure S4**). After filtering, the final cohorts comprised 106 donors for islet density analyses (2,670 images), 93 donors for islet composition analyses (202,585 islets across 835 quadruple-stained images), and 92 donors for islet cell-area analyses.

Processed data were analyzed via a statistical pipeline built on two complementary frameworks (**Figure 1C**). For islet density, total islet cell counts, and cell area, we used generalized linear mixed models (GLMMs) with donor as a random effect, selecting Gamma, negative binomial, or Gaussian (log-link) families based on outcome distributions (see Methods). Islet cell composition (β-, α-, δ-, PP-cell proportions) and the β/(α + β) ratio are outcomes naturally bounded between 0 and 1 with substantial inflation at the extremes in T1D. We therefore used Bayesian ordered beta regression with donor and donor × image as nested random effects. All models accommodated full two-way interactions among diagnosis, region, sex, and age (with islet size added for composition and count outcomes), together with key three-way diagnosis × region × age and diagnosis × region × islet size interactions. Disease duration and age at onset were modeled separately within T1D-only sub-models. Throughout, we report frequentist p-values for GLMM outcomes (Tukey- or Holm-adjusted for multiple comparisons), and posterior medians with 95% highest posterior density (HPD) intervals together with posterior probabilities (pd) for Bayesian outcomes.

### Pancreatic islet density is reduced in T1D with regional and sex-dependent heterogeneity

We first asked whether pancreatic islet density varied as a function of diagnosis, sex, region, age, and disease duration (**Figure 2**, **Supplementary Figure S5**, **Supplementary Table S2**). Averaged across covariates, T1D donors exhibited a significant reduction in islet density relative to ND donors [ND: 1.95 islets/mm^2^ (95% CI: 1.73, 2.20); T1D: 1.39 islets/mm^2^ (95% CI: 1.19, 1.63); −28.7%, p = 0.0008; **Figure 2A**]. Examined as a function of disease duration, however, this reduction did not progress significantly within the T1D cohort: a likelihood-ratio test for disease-duration effects (modeled with a natural cubic spline) was not significant (p = 0.16), and the spline did not improve over a linear term (p = 0.09), indicating that islet density is largely set early and remains stable across the spectrum of disease durations sampled (**Figure 2B**). Additionally, disease duration was not significant when marginalized over region and sex (p = 0.660; **Supplementary Table S2**). Likewise, age at T1D onset showed no significant effect on density (slope = −0.0082; −0.8% per year; p = 0.431; **Figure 2C**).

**Figure 2.**
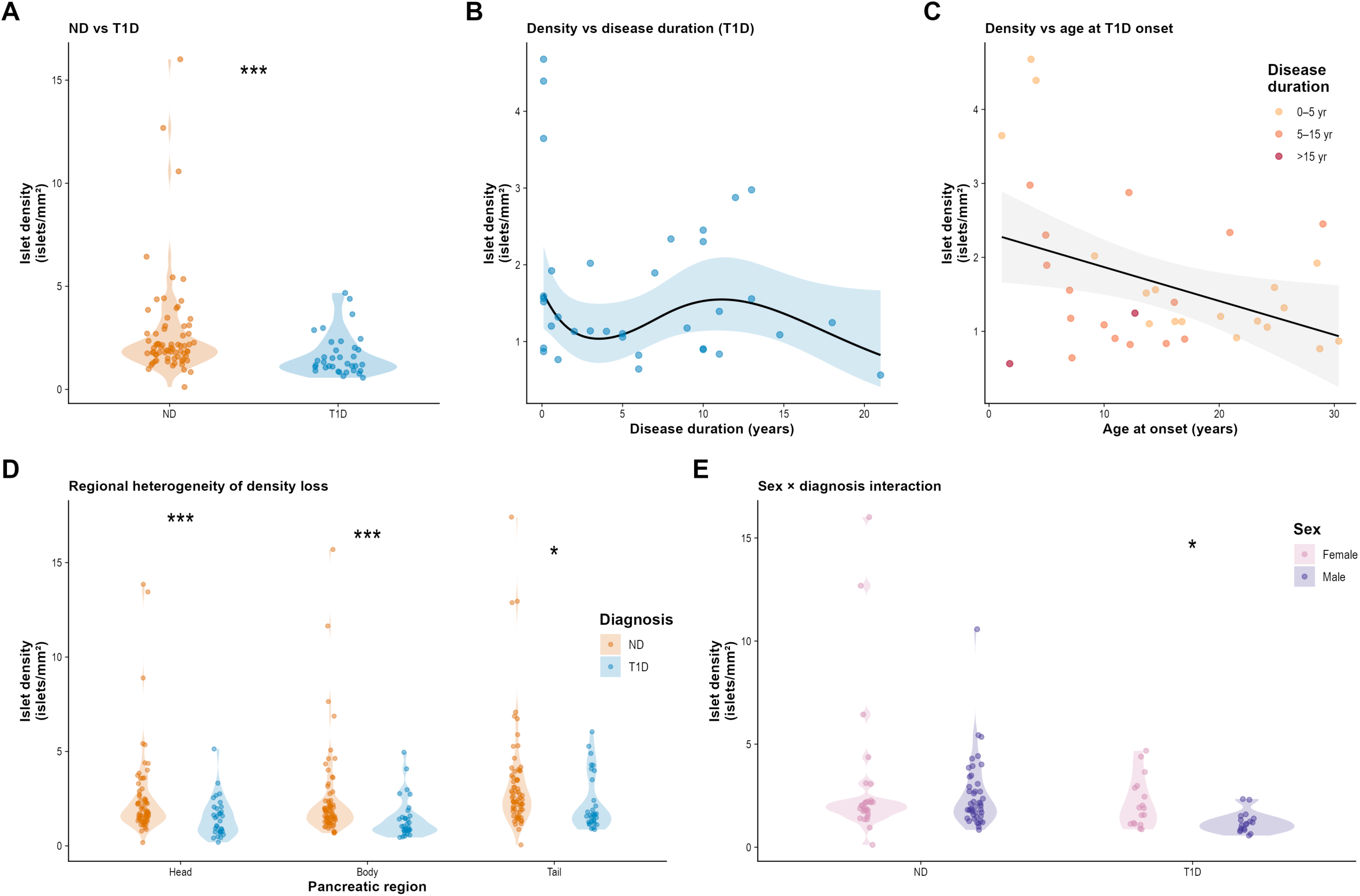
Islet density is reduced in T1D with regional, sex-dependent, and age-related heterogeneity. **(A)** Islet density (islets/mm^2^) in non-diabetic (ND, n = 71) and type 1 diabetic (T1D, n = 35) donors. There is a significant decrease in islet density in T1D donors compared to ND donors (p = 0.0008). Colored points represent individual donor means. **(B)** Islet density as a function of disease duration in T1D donors. The fitted line is from a Gamma GLMM with a natural spline (df = 3) on disease duration. A likelihood-ratio test against a null model without disease duration was not significant (p = 0.16), and the spline did not improve significantly over a linear duration term (p = 0.09), indicating that islet density does not change appreciably with disease duration in T1D donors. **(C)** Islet density as a function of age at T1D onset, with points colored by disease-duration category (0-5, 5-15, >15 years). The black line shows a linear trend with 95% confidence band; the model-adjusted age-at-onset slope (averaged over region and sex) is not statistically significant (−0.8%/year, p = 0.431). **(D)** Islet density by pancreatic region (head, body, tail) stratified by diagnosis. The density ratio (ND/T1D) is greatest in the pancreatic head (ratio 1.55, ND - T1D = 35.5% difference; p < 0.0001), intermediate in the body (ratio 1.43, ND - T1D = 30.2%; p = 0.0006), and smallest in the tail (ratio 1.24, ND - T1D = 19.5%; p = 0.0377). **(E)** Islet density stratified by sex and diagnosis. T1D males exhibit significantly reduced islet density compared to T1D females (ratio 1.47; p = 0.0192). No sex difference is observed in ND donors (ratio 0.97, p = 0.81). Points represent individual donor means. *p<0.05, **p<0.01, ***p<0.001.

Region of the pancreas exerted a strong effect on density. Across all donors, density was highest in the tail (2.16 islets/mm^2^; 95% CI: 1.95, 2.40) compared to the head (1.47 islets/mm^2^; 95% CI: 1.32, 1.63) or body (1.41 islets/mm^2^; 95% CI: 1.27, 1.57) (Head - Tail p < 0.0001; Body - Tail p < 0.0001). The magnitude of the T1D-associated reduction differed across regions, with the largest deficit in the head (ND - T1D = 35.5%, p < 0.0001), an intermediate reduction in the body (30.2%, p = 0.0006), and the smallest in the tail (19.5%, p = 0.0377) (**Figure 2D**, **Supplementary Table S2**). A pronounced sex × diagnosis interaction was also apparent: T1D males showed a marked density reduction relative to ND males (−42.1%, p < 0.0001), whereas T1D females did not differ significantly from ND females (−12.1%, p = 0.393). Within T1D donors, females retained 47.3% higher islet density than males (p = 0.019); there were no sex differences between males and females in the ND cohort (p = 0.807) (**Figure 2E**).

Donor age was associated with an overall decline in islet density across the cohort (−2.4% per year, p < 0.001; **Supplementary Figure S5A**). The ND versus T1D age-slope contrast was not significant (ND - T1D = −0.0088, p = 0.456), indicating comparable age-related slopes in ND and T1D donors averaged across regions. However, the three-way diagnosis × region × age interaction was significant: within T1D donors, age-related density decline was steepest in the head and significantly attenuated in the body (Head - Body p = 0.023) and tail (Head - Tail p = 0.001), with no equivalent regional slope differences in ND donors (all p > 0.05) (**Supplementary Figure S5C**). The Female - Male age-slope contrast was also not significant (−0.0041, p = 0.664; **Supplementary Figure S5D**). Within regions, there were no significant sex differences (all p > 0.05) (**Supplementary Figure S5E**). Within T1D donors, region-stratified disease-duration slopes also revealed regional heterogeneity: the head exhibited a near-flat or weakly negative duration slope, whereas the body and tail showed weakly positive slopes, with significant pairwise differences (Head – Body p = 0.004; Head - Tail p = 0.0001). Together, these results indicate that: 1) islet density is reduced in T1D with concentrated losses in the pancreatic head; 2) males are disproportionately affected; 3) and that surviving islets do not undergo further density loss as a function of disease duration.

### β-cell proportion is markedly reduced in T1D, with significant effects of pancreatic region and islet size

We next assessed the proportion of β-cells within islets across donor diagnosis, sex, region, age, and islet size (**Figure 3**, **Supplementary Figures S6-S7**, **Supplementary Table S2**). Heatmaps of donor-level β-cell proportions stratified by region, sex, and age group, together with split-violin distributions, revealed striking, near-complete β-cell loss in T1D donors across all subgroups (**Figure 3A-B**). Quantitatively, mean β-cell proportion fell from 60.9% (95% HPD: 53.5%, 67.8%) in ND donors to 2.6% (95% HPD: 1.3%, 4.4%) in T1D donors (posterior probability of ND > T1D >0.999), representing the single largest cellular shift observed in our study. Although β-cell loss was near-complete on average, the per-donor distribution was right-skewed: 3 of 34 T1D donors retained β-cell proportions within the non-diabetic range (43-81%), and these were predominantly the most recently diagnosed, all of whom had a disease duration of <0.6 years (see Figure 7E), consistent with the steep duration-dependent decline in β-cell content.

**Figure 3.**
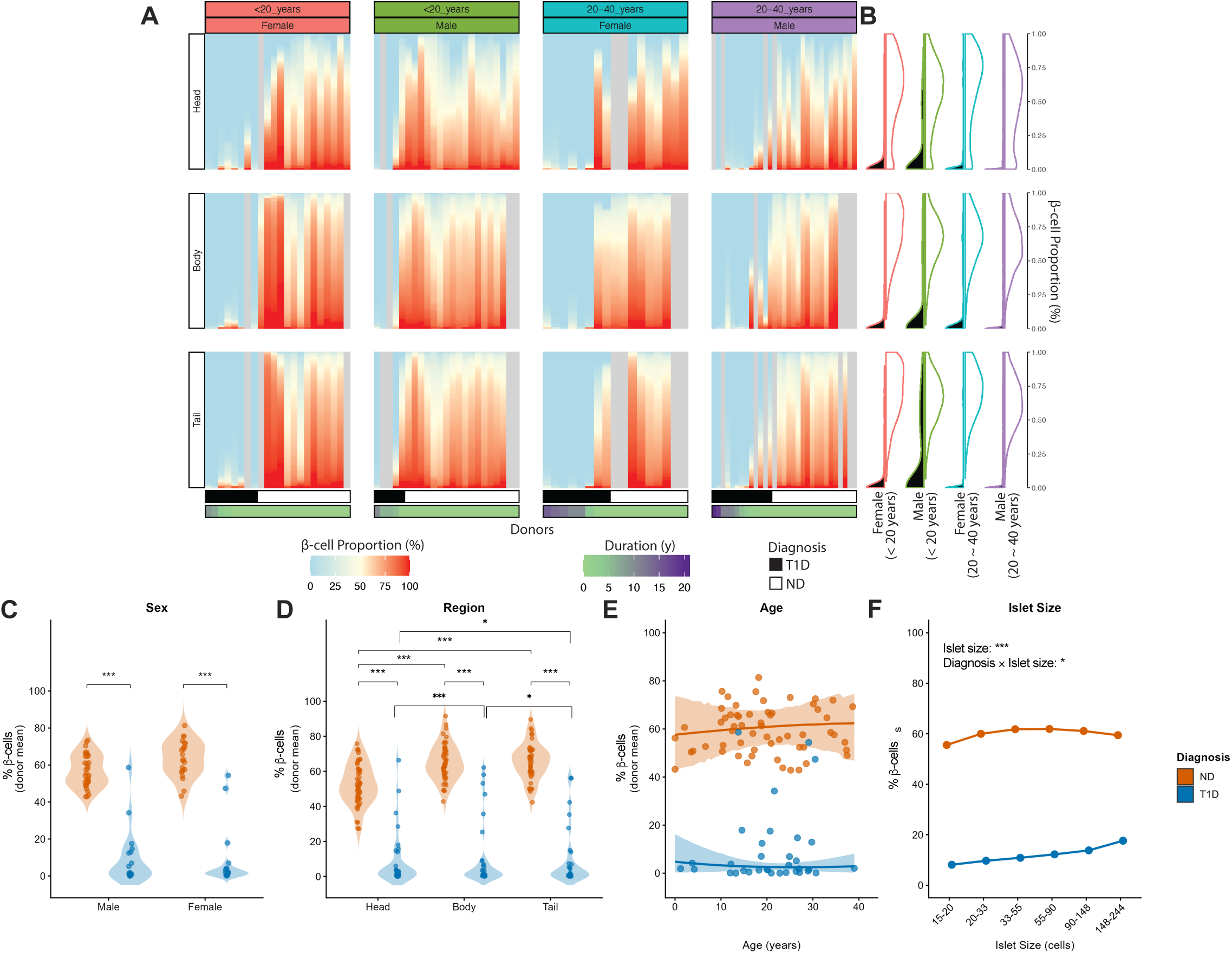
β-cell proportion is reduced in T1D, with significant effects of pancreatic region and islet size. **(A)** Heatmaps of β-cell proportion (% insulin-positive islet cells) across individual donors (columns) and pancreatic regions (head, body, tail; rows), stratified by age group and sex. Donors within each panel are ordered by diagnosis (T1D, black bar; ND, white bar) and disease duration. **(B)** Split violin plots displaying the distribution of β-cell proportions for T1D (left, black) and ND donors (right, white). Outline colors represent stratified groups defined by age group and sex; the plots are vertically aligned with the respective pancreatic regions labeled in Panel A. ND donors (colored curves) show broad distributions centered at high values, while T1D donors (dark curves near zero) show marked β-cell loss. **(C)** β-cell proportion (donor mean) by sex, stratified by diagnosis. Both T1D males and females (blue) exhibit significantly reduced β-cell proportion relative to their ND counterparts (orange; negative posterior probability = 1.0). Colored points represent individual donor means. No significant main effect of sex is observed (posterior probability of Female > Male = 0.73). **(D)** β-cell proportion (donor mean) by pancreatic region, stratified by diagnosis. The pancreatic head shows significantly lower β-cell proportion than either the body (ND: Head - Body = -10.3%, negative posterior probability = 1.0; T1D: Head - Body = - 1.5%, negative posterior probability = 1.0) or tail (ND: Head - Tail = −10.2%, negative posterior probability = 1.0; T1D: Head - Tail = −0.7%, negative posterior probability = 0.99). Though the body and tail do not differ in ND (Body - Tail = 0.1%, positive posterior probability = 0.53), there is a significant difference in T1D (Body - Tail = 0.7%, positive posterior probability = 0.97). **(E)** β-cell proportion (donor mean) as a function of donor age, stratified by diagnosis. ND donors (orange) maintain relatively stable β-cell proportion across age (posterior probability of positive age slope = 0.63). T1D donors (blue) cluster near zero regardless of age (posterior probability of positive age slope = 0.37). **(F)** β-cell proportion as a function of islet size (binned by cell count), stratified by diagnosis. Larger islets retain a higher proportion of β-cells in both ND and T1D donors (positive posterior probability = 1.0 for both). The islet size effect is significantly stronger in T1D than ND (positive posterior probability of T1D slope > ND slope = 0.99), indicating a diagnosis × islet size interaction. Points represent bin-level means of individual donors; lines connect successive islet size categories. ***Corresponds to posterior probability > 0.999; *corresponds to posterior probability > 0.95.

This reduction was preserved across both sexes (ND - T1D: females = 62.1%, pd > 0.999; males = 54.4%, pd > 0.999; **Figure 3C**) and across all three pancreatic regions (ND - T1D: head = 52.1%, body = 60.8%, tail = 61.6%; all pd > 0.999; **Figure 3D**). Within ND donors, the head exhibited a significantly lower β-cell proportion than the body (Head - Body = -10.3%, pd > 0.999) or tail (Head - Tail = −10.2%, pd > 0.999); these regional differences were diminished but preserved in T1D (Head - Body = −1.5%, pd > 0.999; Head - Tail = −0.7%, pd = 0.99), and a small but significant body-tail difference emerged in T1D islets only (Body - Tail = 0.7%, pd = 0.97) (**Figure 3D**). β-cell proportion did not change significantly with donor age in either diagnostic group (ND: pd of positive slope = 0.63; T1D: pd of positive slope = 0.37) (**Figure 3E**).

Islet size emerged as a particularly informative covariate. In both ND and T1D donors, larger islets retained a higher proportion of β-cells (ND slope = 0.0037, T1D slope = 0.0081, both pd > 0.99). The islet-size slope was significantly steeper in T1D than in ND (T1D - ND = 0.0044, pd = 0.99), indicating that surviving β-cells in T1D were disproportionately concentrated within larger islets (**Figure 3F**). Four- and three-way subgroup analyses corroborated these findings: across every combination of region, sex, and age, β-cell proportion was significantly reduced in T1D (**Supplementary Figure S6**). The largest ND - T1D difference was observed in the body of female donors (estimate = 0.652, pd > 0.999) while the smallest ND - T1D difference was in the male head (estimate = 0.455, pd > 0.999) (**Supplementary Figure S7A**). There were no significant diagnosis × age interactions in either males or females (no pd > 0.95; **Supplementary Figure S7B**). A significant diagnosis × islet-size interaction was observed in females (estimate = 0.013, pd > 0.999) but not in males (pd = 0.89) (**Supplementary Figure S7C**). Together, these results identify diagnosis as the dominant driver of β-cell loss, with additional contributing effects provided by pancreatic region and islet size.

### α-cell proportion is markedly increased in T1D islets

In parallel with β-cell loss, we observed reciprocal increases in α-cell proportion (**Figure 4**, **Supplementary Figures S8-S9**, **Supplementary Table S2**). Donor-level heatmaps and accompanying split-violin distributions showed elevated α-cell content across all subgroups in T1D donors (**Figure 4A-B**). Quantitatively, mean α-cell proportion increased from 14.3% (95% HPD: 11.8%, 17.1%) in ND to 60.4% (95% HPD: 54.9%, 66.1%) in T1D donors (ND - T1D = - 46.1%, pd > 0.999), with no significant main effect of sex (ND versus T1D: females −48.1%, males −44.2%, both pd > 0.999; sex contrast pd = 0.54) (**Figure 4C**).

**Figure 4.**
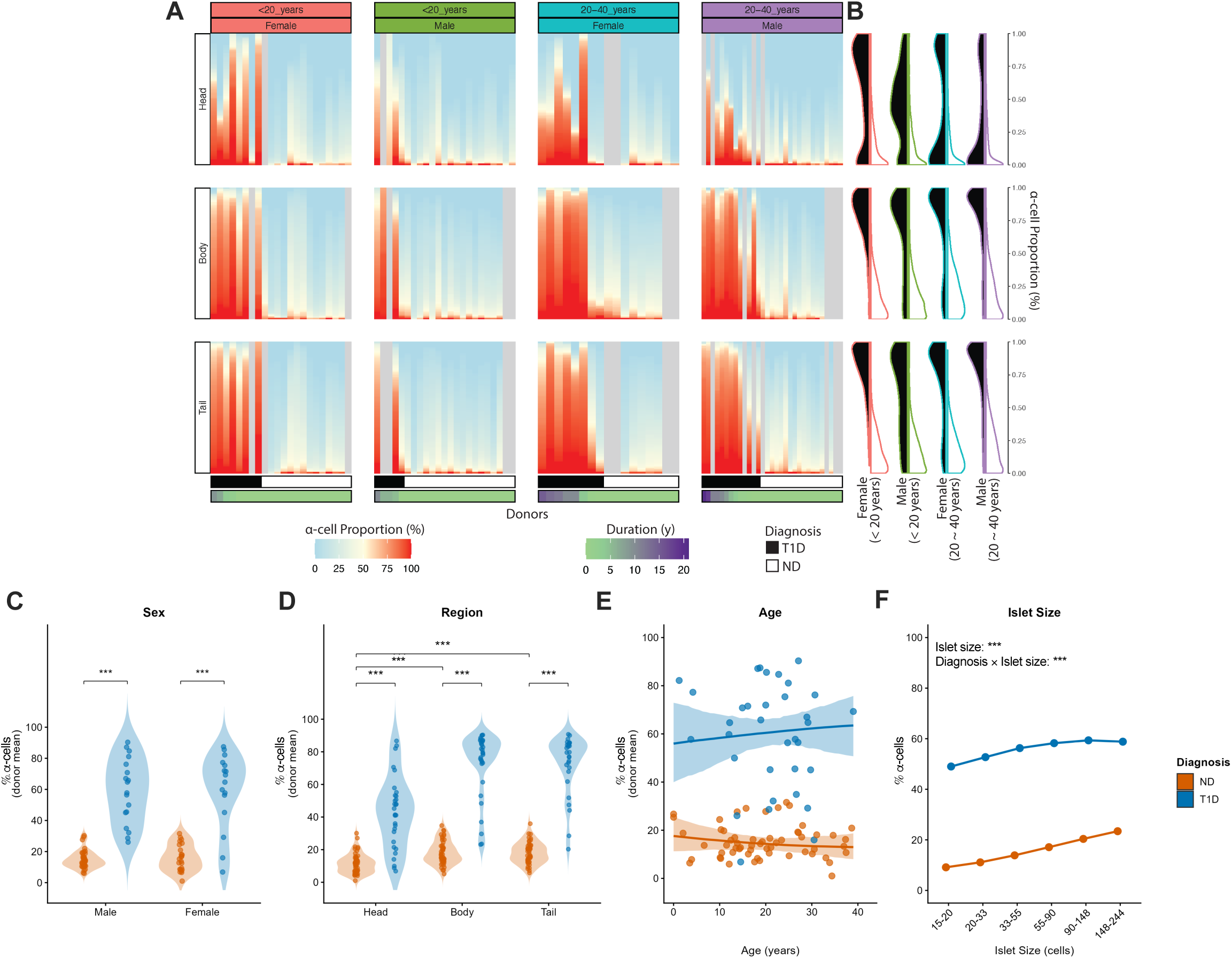
α-cell proportion is markedly increased in T1D islets. **(A)** Heatmaps of α-cell proportion (% glucagon-positive islet cells) across individual donors (columns) and pancreatic regions (head, body, tail; rows), stratified by age group and sex. Donors within each panel are ordered by diagnosis (T1D, black bar; ND, white bar) and disease duration. **(B)** Split violin plots displaying the distribution of α-cell proportions for T1D (left, black) and ND donors (right, white). Outline colors represent stratified groups defined by age group and sex; the plots are vertically aligned with the respective pancreatic regions labeled in Panel A. T1D donors (dark curves) show elevated α-cell content, while ND donors (colored curves) show lower, tightly distributed values. **(C)** α-cell proportion (donor mean) by sex, stratified by diagnosis. Both T1D males and females (blue) exhibit increased α-cell proportion compared to ND (orange; positive posterior probability = 1.0). Colored points represent individual donor means. No significant main effect of sex is observed (positive posterior probability of Female > Male = 0.54). **(D)** α-cell proportion (donor mean) by pancreatic region, stratified by diagnosis. There is a significant diagnosis × region interaction indicating a larger difference in α-cell proportion within islets of the body and tail than in the head of T1D versus ND donors (T1D - ND difference: Head = 33.5%, Body = 51.2%, Tail = 53.6%; all positive posterior probabilities = 1.0). A regional difference is also present in both ND (Head – Body = −7.3%, negative posterior probability = 1.0; Head – Tail = −7.3%, negative posterior probability = 1.0; Body– Tail = 0.04%, positive posterior probability = 0.51) and T1D (Head – Body = −24.9%, negative posterior probability = 1.0; Head - Tail = −27.3%, negative posterior probability = 1.0; Body - Tail = −2.4%, negative posterior probability = 0.89). **(E)** α-cell proportion (donor mean) as a function of donor age, stratified by diagnosis. Neither ND donors (posterior probability of negative age slope = 0.79) nor T1D donors (posterior probability of positive age slope = 0.70) show a significant age trend, and the diagnosis × age interaction is not significant (posterior probability of T1D slope > ND slope = 0.78). **(F)** α-cell proportion as a function of islet size (binned by cell count), stratified by diagnosis. Larger islets contain a higher proportion of α-cells in both ND and T1D donors (positive posterior probability = 1.0 for both). The islet size effect is significantly stronger in ND than T1D donors (T1D - ND slope difference = -0.02; negative posterior probability of T1D slope < ND slope = 1.0), indicating a diagnosis × islet size interaction. Points represent bin-level means of individual donors; lines connect successive islet size categories. ***Corresponds to posterior probability > 0.999.

A significant diagnosis × region interaction was evident, with the T1D-associated increase in α-cell proportion being larger in the body and tail than in the head (T1D - ND: head = 33.5%, body = 51.2%, tail = 53.6%; all pd > 0.999). Within ND donors, the head harbored a lower proportion of α-cells than the body or tail (Head - Body = −7.3%, Head - Tail = −7.3%; both pd > 0.999). These regional differences persisted in T1D donors and were further magnified (Head - Body = −24.9%, Head - Tail = −27.3%; both pd > 0.999) (**Figure 4D**). α-cell proportion showed no significant overall age effect in either diagnostic group (ND: pd of negative slope = 0.79; T1D: pd of positive slope = 0.70; diagnosis × age pd = 0.78) (**Figure 4E**).

As with β-cells, islet size was strongly associated with α-cell proportion. Larger islets contained a higher proportion of α-cells in both ND (slope = 0.054, pd > 0.999) and T1D donors (slope = 0.031, pd > 0.999) (**Figure 4F**). Four-way and three-way subgroup analyses confirmed that α-cell increases in T1D were robust across every combination of region, sex, and age (**Supplementary Figures S8-S9**, **Supplementary Table S2**). Diagnosis was again the primary driver alongside regional effects, particularly within the body and tail.

### The β/(α+β) ratio collapses in T1D and continues to decline with disease duration

Given the reciprocal changes in islet α-cell and β-cell composition in response to T1D, we further examined how these cell types changed relative to one another by calculating the islet β/(α+β) ratio (**Supplementary Figure S10**, **Supplementary Table S2**). Averaged across covariates, the ratio collapsed ∼90-fold from 0.889 (95% HPD: 0.835, 0.932) in ND donors to 0.010 (95% HPD: 0.003, 0.022) in T1D donors (ND - T1D = 0.88, pd > 0.999). This near-complete shift was preserved across both sexes (ND - T1D: females = 0.88, males = 0.88; both pd > 0.999; **Supplementary Figure S10B**) and across all three pancreatic regions [Head: ND, 0.927 (95% HPD: 0.889, 0.959) to T1D, 0.009 (95% HPD: 0.002, 0.020); Body: ND, 0.874 (95% HPD: 0.816, 0.927) to T1D, 0.014 (95% HPD: 0.004, 0.029); Tail: ND, 0.867 (95% HPD: 0.806, 0.921) to T1D, 0.008 (95% HPD: 0.002, 0.017); all pd > 0.999; **Supplementary Figure S10A**]. In ND donors, the head maintained a significantly higher ratio than either the body or tail (both pd > 0.999); this regional pattern was largely abolished in T1D, although a small body-tail difference persisted (Body - Tail = 0.006, pd > 0.999) (**Supplementary Figure S10A**).

Islet size and disease duration were also associated with changes to the β/(α+β) ratio. In ND donors, the ratio declined modestly with increasing islet size (slope = −0.039, pd > 0.999), reflecting a relative enrichment of α-cells in larger islets. By contrast, in islets from T1D donors, the slope was near zero (slope = 0.002, pd > 0.999; T1D - ND interaction pd > 0.999) (**Supplementary Figure S10C**). Donor age had no significant effect on the ratio in either group (**Supplementary Figure S10D**). Within T1D donors, however, the β/(α+β) ratio declined significantly with disease duration (overall slope = −0.062 per year, pd > 0.999), with consistent declines in the head (−0.053), body (−0.075), and tail (−0.058), all pd > 0.999 (**Supplementary Figure S10E**). Age at T1D onset showed no significant association with the ratio (slope = 0.006, pd = 0.78) (**Supplementary Figure S10F**). These data therefore suggest that disease duration is a more meaningful indicator of changes to the β/(α+β) ratio compared to age of T1D onset.

### δ-cells are largely resilient to T1D-induced changes

In contrast to β- and α-cells, δ-cells were largely preserved across diagnostic groups (**Figure 5**, **Supplementary Figures S11-S12**, **Supplementary Table S2**). Donor-level heatmaps and split-violin distributions showed broadly overlapping δ-cell distributions in ND and T1D donors across all subgroups (**Figure 5A-B**), and no significant main effect of diagnosis on δ-cell proportion [ND = 14.0% (95% HPD: 12.5%, 15.6%), T1D = 14.4% (95% HPD: 12.5%, 16.7%); ND - T1D = -0.4%, pd = 0.61]. We did, however, identify an effect of donor sex: in ND donors, males exhibited a higher δ-cell proportion than females [ND males = 15.9% (95% HPD: 13.9%, 18.2%), ND females = 12.1% (95% HPD: 9.9%, 14.4%); pd of male > female = 0.99]; a similar but smaller, non-significant trend was observed in T1D donors [T1D males = 15.0% (95% HPD: 12.0%, 18.5%), T1D females = 13.7% (95% HPD: 10.9%, 16.6%); pd = 0.70] (**Figure 5C**). No significant diagnosis × region interactions were observed (**Figure 5D**).

**Figure 5.**
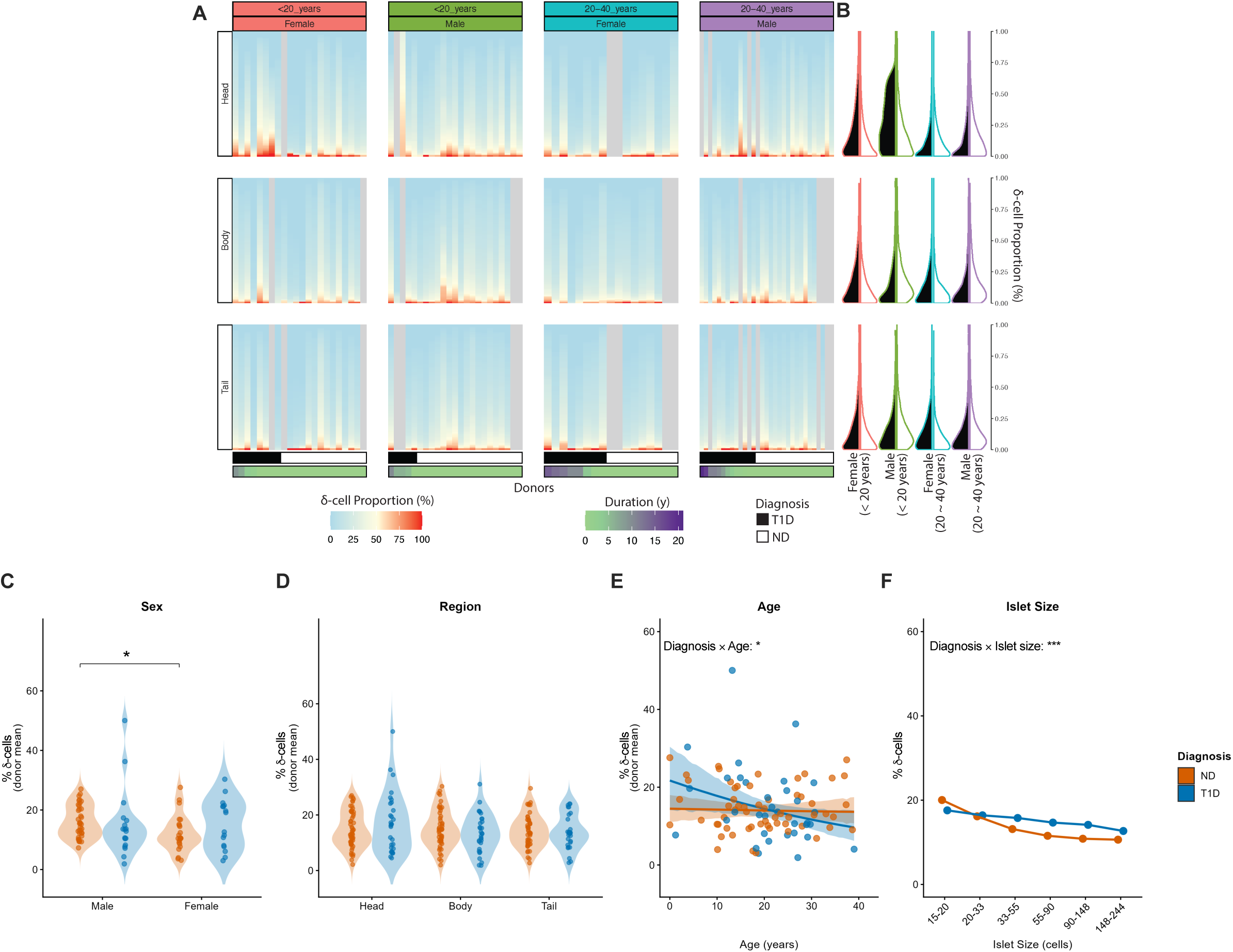
δ-cell proportion is unaffected by T1D diagnosis. **(A)** Heatmaps of δ-cell proportion (% somatostatin-positive islet cells) across individual donors (columns) and pancreatic regions (head, body, tail; rows), stratified by age group and sex. Donors within each panel are ordered by diagnosis (T1D, black bar; ND, white bar) and disease duration. **(B)** Split violin plots displaying the distribution of δ-cell proportions for T1D (left, black) and ND donors (right, white). Outline colors represent stratified groups defined by age group and sex; the plots are vertically aligned with the respective pancreatic regions labeled in Panel A. The distributions largely overlap between T1D and ND donors, consistent with the absence of a significant effect of diagnosis. **(C)** δ-cell proportion (donor mean) by sex, stratified by diagnosis. Male ND donors show higher δ-cell proportion than female ND donors (ND Female = 12.1%, ND Male = 15.9%; posterior probability of Male > Female = 0.99), while T1D donors show a weaker, non-significant difference (T1D Female = 13.7%, T1D Male = 15.0%; posterior probability of Male > Female = 0.70). **(D)** δ-cell proportion (donor mean) by pancreatic region, stratified by diagnosis. There is no significant diagnosis × region interaction in δ-cell proportion in both T1D and ND donors. **(E)** δ-cell proportion (donor mean) as a function of donor age, stratified by diagnosis. ND donors show no significant age trend (posterior probability of negative age slope = 0.60). T1D donors show a modest negative association with age (posterior probability of negative age slope = 0.98), with a significant diagnosis × age interaction (posterior probability of T1D slope < ND slope = 0.96; T1D-ND = - 0.28%/year). **(F)** δ-cell proportion as a function of islet size (binned by cell count), stratified by diagnosis. Larger islets contain a lower proportion of δ-cells in both ND and T1D donors (negative posterior probability = 1.0 for both). The islet size effect is significantly attenuated in T1D relative to ND (T1D - ND slope difference = 0.02; positive posterior probability = 1.0), indicating a diagnosis × islet size interaction. Points represent bin-level means of individual donors; lines connect successive islet size categories. *Corresponds to posterior probability > 0.95; ***corresponds to posterior probability > 0.999.

Though we identified a statistically significant diagnosis × age interaction (T1D - ND slope difference = −0.003 per year, pd of T1D slope < ND slope = 0.96), annual decreases in δ-cell proportion in T1D islets were negligible (−0.003 per year; pd of negative slope = 0.98); there was no significant effect in ND donors (pd = 0.60) (**Figure 5E**). Islet size negatively predicted δ-cell proportion in both groups, with larger islets containing fewer δ-cells (ND slope = −0.038, T1D slope = −0.016; both pd > 0.999) (**Figure 5F**).

According to three- and four-way subgroup analyses (**Supplementary Figures S11-S12**), within the head of T1D males, δ-cell proportion declined significantly with age (slope = −0.0091, pd = 0.998). Moreover, a flattened diagnosis × islet-size slope was observed in the head of both sexes (females: T1D - ND = 0.022, pd > 0.999; males: T1D - ND = 0.025, pd > 0.999), extending to the body (T1D - ND = 0.045, pd > 0.999) and tail (T1D - ND = 0.037, pd > 0.999) in males but not in females. Together, these results indicate that δ-cells are broadly preserved in T1D but exhibit sex-and region-specific responses to disease and aging.

### PP-cells are concentrated in the pancreatic head and largely unaffected by T1D

PP-cells were the sparsest islet cell population analyzed and exhibited the strongest regional patterning (**Figure 6**, **Supplementary Figures S13-S14**, **Supplementary Table S2**). Donor-level heatmaps revealed that PP-cell content is overwhelmingly concentrated within the pancreatic head, with markedly lower levels in the body and tail across all donors (**Figure 6A-B**). Importantly, no significant effect of diagnosis was observed when averaged across covariates [ND = 5.9% (95% HPD: 4.4%, 7.3%), T1D = 6.4% (95% HPD: 4.4%, 8.6%); ND - T1D = −0.5%, pd = 0.63].

**Figure 6.**
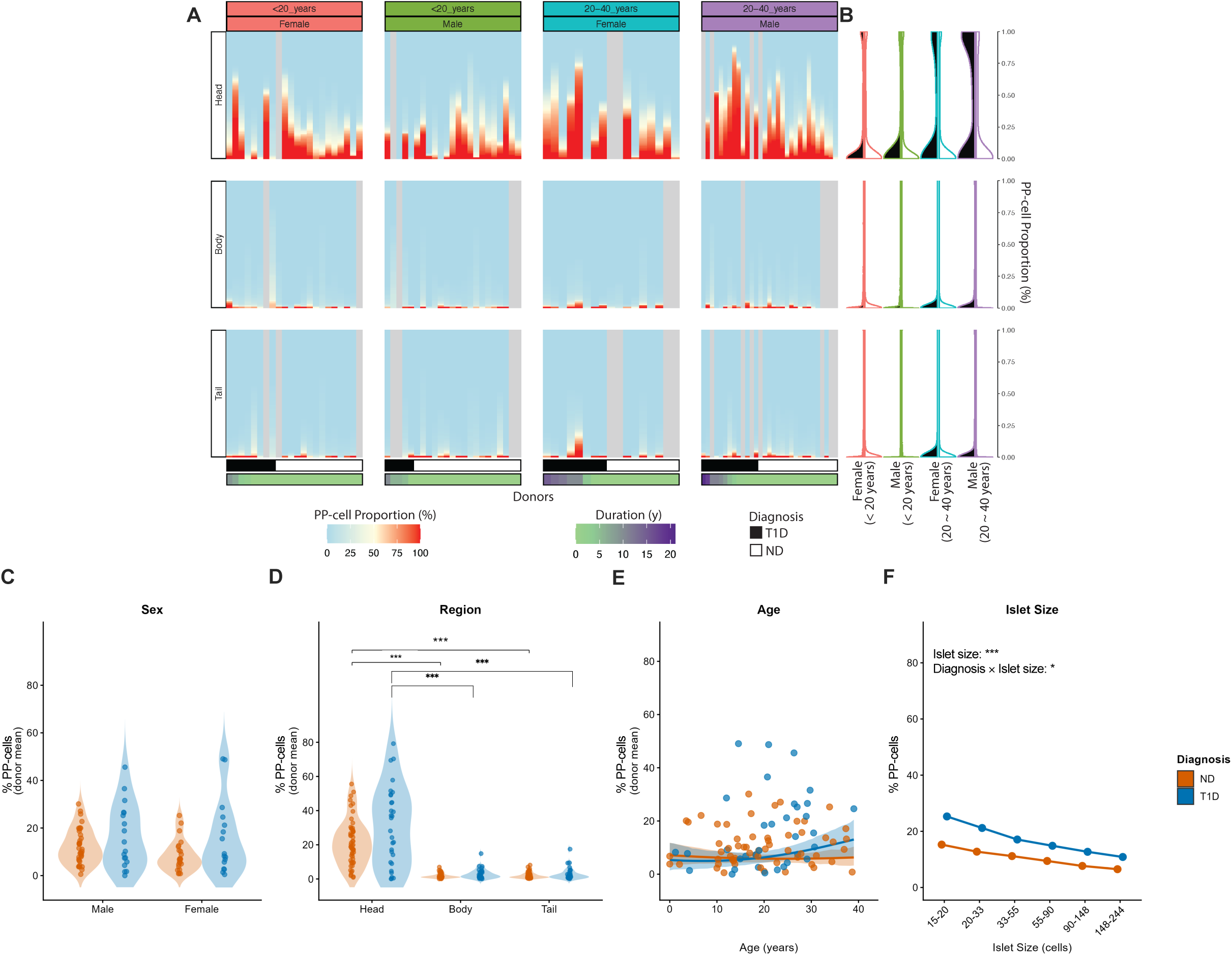
PP-cell proportion is primarily enriched in the pancreatic head and unaffected by T1D. **(A)** Heatmaps of PP-cell proportion (% PP-positive islet cells) across individual donors (columns) and pancreatic regions (head, body, tail; rows), stratified by age group and sex. Donors within each panel are ordered by diagnosis (T1D, black bar; ND, white bar) and disease duration. Islet PP-cell content is primarily concentrated within the pancreatic head. **(B)** Split violin plots displaying the distribution of PP-cell proportions for T1D (left, black) and ND donors (right, white). Outline colors represent stratified groups defined by age group and sex; the plots are vertically aligned with the respective pancreatic regions labeled in Panel A. **(C)** PP-cell proportion (donor mean) by sex, stratified by diagnosis. No significant effect of diagnosis is observed within both males (ND - T1D = 0.6%, positive posterior probability ND > T1D = 0.6) and females (ND - T1D = -1.6%, negative posterior probability ND < T1D = 0.8). **(D)** PP-cell proportion (donor mean) by pancreatic region, stratified by diagnosis. Regionally, most PP-cells are concentrated in the pancreatic head with markedly less in the body and tail. This relationship is preserved in both ND donors (Head - Body = 9.7%, positive posterior probability = 1.0; Head - Tail = 9.5%, positive posterior probability = 1.0) and T1D donors (Head - Body = 12.5%, positive posterior probability = 1.0; Head - Tail = 12.4%, positive posterior probability = 1.0). **(E)** PP-cell proportion (donor mean) as a function of donor age, stratified by diagnosis. ND donors show no significant age trend (posterior probability of negative age slope = 0.59). **(F)** PP-cell proportion as a function of islet size (binned by cell count), stratified by diagnosis. Larger islets contain a lower proportion of PP-cells in both ND and T1D donors (negative posterior probability = 1.0 for both). The islet size effect is significantly attenuated in T1D relative to ND (T1D - ND slope difference = -0.008; negative posterior probability = 0.99), indicating a diagnosis × islet size interaction. Points represent bin-level means of individual donors; lines connect successive islet size categories. All estimates are posterior medians with 95% highest posterior density intervals. *Corresponds to posterior probability > 0.95; ***corresponds to posterior probability > 0.999.

**Figure 7.**
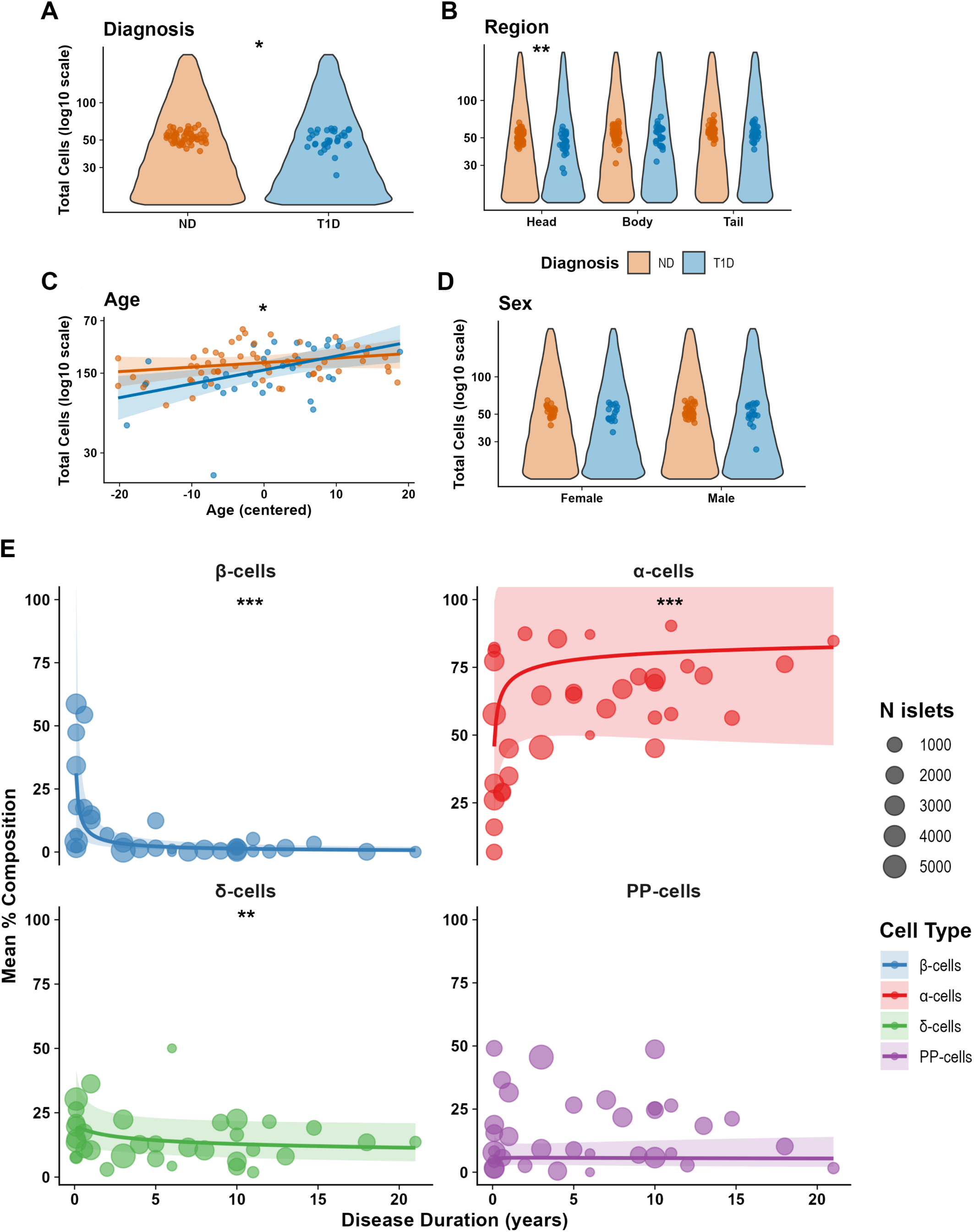
Total islet cell counts remain relatively stable in surviving islets of T1D donors with disease duration reshaping cell-type composition. **(A)** Violin plots of total islet cell counts (log_10_ scale) by diagnosis. T1D donors (blue) exhibit modestly lower total cell counts in surviving islets compared to ND donors (orange) (predicted means: ND = 53.4, T1D = 50.9; −4.8%, p = 0.039). **(B)** Violin plots of total islet cell counts by pancreatic region of individual donors, stratified by diagnosis. Within the pancreatic head, the effect of diagnosis is significant with a cell count reduction in T1D versus ND donors (−7.5%, p = 0.004); there is no significant effect of diagnosis in the body (p = 0.309) or tail (p = 0.147) regions. **(C)** Total islet cell counts as a function of donor age, stratified by diagnosis. Both groups show a positive age-cell count association with total islet cell counts increasing as a function of age. T1D donors exhibit a significantly faster rate of increase compared to ND donors (0.9% for T1D donors vs 0.3% for ND donors per year; p = 0.039). Lines represent model-predicted trends with 95% confidence bands. **(D)** Violin plots of total islet cell counts by sex, stratified by diagnosis. No significant main effect of sex is observed (Female = 52.4, Male = 51.9; p = 0.726), and the diagnosis × sex interaction is not significant (p = 0.852). For Panels A-D: *Corresponds p < 0.05; **p < 0.01. **(E)** Mean percent composition of β-cells, α-cells, δ-cells, and PP-cells as a function of disease duration (years) in individual T1D donors only. Shaded ribbons indicate 95% confidence intervals. β-cell proportion declines sharply in early disease (−2.9%/year of disease duration, negative posterior probability > 0.999) and approaches near-zero at longer durations, while α-cell proportion increases reciprocally (2.9%/year of disease duration, positive posterior probability > 0.999). δ-cell (−1.2%/year of disease duration, negative posterior probability = 0.999) and PP-cell (0.5%/year of disease duration, positive posterior probability = 0.809) proportions remain relatively stable across disease duration. **Corresponds to posterior probability = 0.999; ***corresponds to posterior probability > 0.999.

We similarly found no diagnosis × sex interaction nor sex as a main effect (**Figure 6C**). Consistent with our qualitative data, regional concentration was apparent in both ND donors [Head: 12.3% (95% HPD: 9.2%, 15.6%), Body: 2.5% (95% HPD: 1.7%, 3.4%), Tail: 2.7% (95% HPD: 1.9%, 3.7%); Head - Body and Head - Tail, both pd > 0.999] and T1D donors [Head: 14.6% (95% HPD: 9.7%, 19.7%), Body: 2.1% (95% HPD: 1.3%, 3.2%), Tail: 2.3% (95% HPD: 1.3%, 3.3%); Head - Body and Head - Tail both pd > 0.999] (**Figure 6D**). In contrast, donor age was not significantly associated with PP-cell proportion overall (**Figure 6E**). Notably, we also examined PP-cell proportion as a function of islet size. There was a strongly negative association with PP-cell proportion where larger islets contained a smaller proportion of PP-cells in ND and T1D groups (ND: -0.015, T1D: -0.023; both pd > 0.999), with a modestly steeper decline in T1D than in ND (T1D - ND slope = -0.008, pd = 0.987) (**Figure 6F**). This is consistent with the heavy concentration of PP-cells in smaller islets within the pancreatic head.

Three- and four-way subgroup analyses revealed region-dependent T1D effects (**Supplementary Figures S13-S14**). Within the head of T1D males, PP-cell proportion increased significantly with age (slope = 0.016, pd = 0.998) and this increase was quicker in the head of T1D donors as the cohort aged (T1D - ND age-slope difference = 0.015, pd = 0.995). Subgroup analyses also showed that in the head, PP-cell proportion declined more steeply with increasing islet size in T1D females compared to ND females (T1D - ND = -0.042, pd = 0.998), whereas in the body, the islet-size slope became significantly less negative in T1D in both sexes (females: T1D - ND = 0.008, pd > 0.999; males: T1D - ND = 0.005, pd > 0.999). No significant T1D - ND differences were observed in the tail. Collectively, these results indicate that PP-cells are largely resilient to T1D-induced changes in islet composition, with their distribution principally governed by pancreatic region.

### Total islet cell counts remain relatively stable in surviving T1D islets

Given the magnitude of the compositional shifts described above, we examined total islet cell counts in pancreatic islets of ND and T1D donors (**Figure 7, Supplementary Table S2**). Averaged across region and sex, total cell counts were only modestly lower in T1D islets compared to ND [ND = 53.4 cells/islet (95% CI: 52.0, 55.0); T1D = 50.9 cells/islet (95% CI: 49.0, 52.8); -4.8%, p = 0.039] (**Figure 7A**). This effect was driven primarily by the pancreatic head, the only region showing a significant T1D - ND difference (−7.5%, p = 0.004); no significant differences were observed in the body (p = 0.31) or tail (p = 0.15) (**Figure 7B**). Moreover, donor age was positively associated with total islet cell counts in both diagnostic groups. As donor age increased, the average number of cells within islets also significantly rose, albeit the rate of increase in T1D was steeper (T1D slope = 0.88%/year) than in ND (ND slope = 0.29%/year) (T1D - ND slope difference, p = 0.039). These data indicated that surviving T1D islets tended to be larger in older donors than would be predicted from the ND age trend alone (**Figure 7C**). In contrast, no significant main effect of sex was observed (p = 0.73), nor any diagnosis × sex interaction (p = 0.85) (**Figure 7D**).

To examine how cell type composition is reshaped over the course of T1D, we examined per-islet cell type proportions across disease duration in T1D donors (**Figure 7E**, **Supplementary Figure S15**). As expected, β-cell proportion declined steeply with disease duration [-20.3%/year (95% CI: -26.1%, -14.0%)], approaching near-zero at longer durations, while α-cell proportion increased reciprocally [6.9%/year (95% CI: 3.4%, 10.5%)]. In contrast, δ-cell proportion changed only slightly with disease duration [slope = -4.1%/year (95% CI: -7.8%, -0.1%)], and PP-cell proportion showed no significant change [slope = 4.3% per year (95% CI: -2.3%, 11.4%)] – both substantially smaller than the shifts in β-cells and α-cells. Altogether, these results indicate that despite extensive remodeling of islet composition from β-cells to α-cells, the total number of cells within surviving islets remained relatively stable throughout the progression of T1D.

### T1D alters total endocrine cell area in a cell type- and endocrine object type-specific manner

Beyond cell number, we asked whether T1D alters the area occupied by each endocrine cell population within EOs which include single endocrine objects (*i.e*., single cells), SEOs, and islets (**Figure 8**, **Supplementary Figures S16-S18, Supplementary Table S2**). β-cell area was significantly reduced in T1D donors at every level of EO organization. Single cell area fell from 129.8 µm^2^ (95% CI: 113.5, 148.4) in ND donors to 76.4 µm^2^ (95% CI: 61.5, 94.8) (−41.2% decrease, p < 0.001) in T1D donors. SEO β-cell area fell from 316.8 µm^2^ (95% CI: 241.3, 415.9) in ND donors to 23.6 µm^2^ (95% CI: 16.4, 34.0) (−92.6% decrease, p < 0.001) in T1D donors. Finally, islet-level β-cell area fell from 2,725.2 µm^2^ (95% CI: 1,954.9, 3,799.0) in ND donors to 81.1 µm^2^ (95% CI: 52.2, 126.0) (−97.0% decrease, p < 0.001) in T1D donors (**Figure 8A**). The progressive magnitude of β-cell area loss from single cells to SEOs to islets suggests remodeling of individual β-cells at the cellular level within surviving EOs in response to T1D-related insults. T1D-associated changes in α-cell area were more dependent on object type (**Figure 8B**). At the single-cell level, α-cell area was modestly reduced in T1D [ND: 104.4 µm^2^ (95% CI: 95.0, 114.7) to T1D: 71.2 µm^2^ (95% CI: 63.3, 80.2), -31.7% decrease, p < 0.001]. At the SEO level, α-cell area did not differ significantly between groups [ND: 123.0 µm^2^ (95% CI: 108.7, 139.2) to T1D: 129.8 µm^2^ (95% CI: 110.8, 152.2), 5.6% increase, p = 0.60]. At the islet level, however, α-cell area more than doubled in T1D donors [ND: 678.8 µm^2^ (95% CI: 579.2, 795.4) to T1D: 1,419.0 µm^2^ (95% CI: 1,157.0, 1,740.3), 109.1% increase, p < 0.001], reflecting marked expansion of α-cells within surviving islets. δ-cell area showed minimal T1D-related change at the single-cell [ND: 44.8 µm^2^ (95% CI: 37.5, 53.6) to T1D: 41.1 µm^2^ (95% CI: 32.8, 51.5), -8.3% decrease, p = 0.55] and islet [ND: 228.4 µm^2^ (95% CI: 197.7, 264.0) to T1D: 227.2 µm^2^ (95% CI: 188.7, 273.6), -0.5% decrease, p = 0.96] levels but was significantly increased within SEOs [ND: 47.2 µm^2^ (95% CI: 40.0, 55.7) to T1D: 69.4 µm^2^ (95% CI: 55.8, 86.3), 47.0% increase, p = 0.005] (**Figure 8C**). PP-cell area was significantly elevated in T1D donors in every object type, with single-cell, SEO, and islet PP-cell areas all approximately doubling in T1D versus ND. Among single PP-cells, total cell area substantially increased in T1D [ND: 33.7 µm^2^ (95% CI: 27.5, 41.4) to T1D: 78.1 µm^2^ (95% CI: 61.8, 98.8), 131.6% increase; p < 0.001]. PP-cell total area in T1D was also significantly raised both within SEOs [ND: 55.2 µm^2^ (95% CI: 45.7, 66.8) to T1D: 134.6 µm^2^ (95% CI: 107.7, 168.3), 143.6% increase; p < 0.001] and in islets [ND: 170.9 µm^2^ (95% CI: 137.7, 212.0) to T1D: 471.4 µm^2^ (95% CI: 361.3, 615.0), 175.8% increase; p < 0.001] (**Figure 8D**).

**Figure 8.**
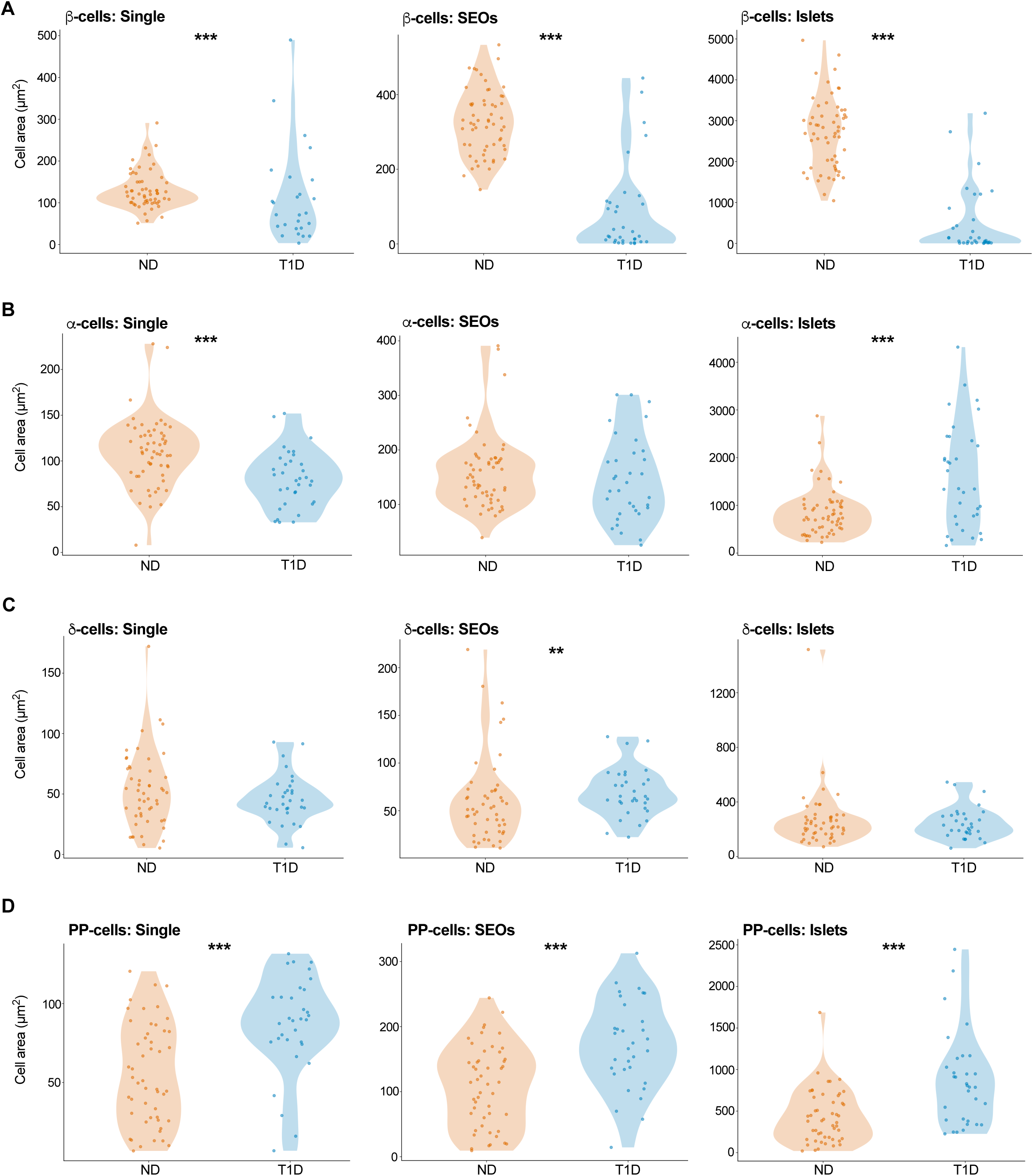
T1D alters total endocrine cell area. **(A)** β-cell area (µm^2^) in single cells (Single), small endocrine objects (SEOs; 2-14 cells per object), and islets (β15 cells per object), comparing ND (orange) and T1D (blue) donors. β-cell area is significantly reduced in T1D across all three object types (Single: 129.8 versus 76.4 µm^2^, -41.2%, p < 0.001; SEOs: 316.8 versus 23.6 µm^2^, -92.6%, p < 0.001; Islets: 2725.2 versus 81.1 µm^2^, -97.0%, p < 0.001). **(B)** α-cell area (µm^2^) by object type. α-cell area is reduced at the single-cell level (104.4 versus 71.2 µm^2^, -31.7%, p < 0.001), unchanged at the SEO level (123.0 versus 129.8 µm^2^, 5.6%, p = 0.596), and markedly increased at the islet level (678.8 versus 1419.0 µm^2^, 109.1%, p < 0.001) in T1D compared to ND donors. **(C)** δ-cell area (µm^2^) by object type. δ-cell area is unchanged at the single-cell (44.8 versus 41.1 µm^2^, -8.3%, p = 0.547) and islet (228.4 versus 227.2 µm^2^, -0.5%, p = 0.964) levels but significantly increased in SEOs (47.2 versus 69.4 µm^2^, 47.0%, p = 0.005) in T1D. **(D)** PP-cell area (µm^2^) by object type. PP-cell area is increased in T1D across all three object types (Single: 33.7 versus 78.1 µm^2^, 131.6%, p < 0.001; SEOs: 55.2 versus 134.6 µm^2^, 143.6%, p < 0.001; Islets: 170.9 versus 471.4 µm^2^, 175.8%, p < 0.001). Each point represents the mean cell area of individual donors; violin plots show the underlying distribution. Values reported are marginal means (ND versus T1D) averaged over region and sex. **p < 0.01; ***p < 0.001.

To distinguish whether these area changes reflected changes in per-cell area versus altered cell numbers, we computed per-cell area for each cell type within SEOs and islets (**Supplementary Figure S16**). β-cells in T1D were markedly smaller within SEOs [ND: 94.2 µm^2^/cell (95% CI: 75.4, 117.7) to T1D: 16.6 µm^2^/cell (95% CI: 12.3, 22.4), -82.4% decrease; p < 0.001] and islets [ND: 93.7 µm^2^/cell (95% CI: 74.6, 117.6) to T1D: 20.2 µm^2^/cell (95% CI: 14.9, 27.3), -78.4% decrease; p < 0.001], indicating individual β-cell atrophy independent of the loss in cell number. α-cells in T1D were also smaller per cell within SEOs [ND: 72.8 µm^2^/cell (95% CI: 65.7, 80.7) to T1D: 38.3 µm^2^/cell (95% CI: 33.6, 43.7), -47.4% decrease; p < 0.001) and islets [ND: 92.0 µm^2^/cell (95% CI: 83.0, 101.9) to T1D: 51.6 µm^2^/cell (95% CI: 45.2, 58.9), -43.9% decrease; p < 0.001), implying that the increase in total α-cell area at the islet level is driven by α-cell hyperplasia rather than hypertrophy. In contrast, δ-cells in T1D SEOs were enlarged on a per-cell basis [ND: 23.8 µm^2^/cell (95% CI: 19.9, 28.6) to T1D: 32.7 µm^2^/cell (95% CI: 25.8, 41.5), 37.1% increase; p = 0.035). There was no change in per-δ-cell area at the islet level [ND: 47.7 µm^2^/cell (95% CI: 40.5, 56.1) to T1D: 43.5 µm^2^/cell (95% CI: 35.3, 53.6), -8.7% decrease; p = 0.49). The per-cell areas of PP-cells in T1D were enlarged in SEOs [ND: 26.4 µm^2^/cell (95% CI: 22.2, 31.2) to T1D: 55.1 µm^2^/cell (95% CI: 45.1, 67.4), 109.2% increase; p < 0.001] and in islets [ND: 61.7 µm^2^/cell (95% CI: 51.5, 73.9) to T1D: 123.5 µm^2^/cell (95% CI: 98.6, 154.6), 100.1% increase; p < 0.001], indicating more general PP-cell hypertrophy across object types in T1D.

We next examined the impact of disease duration on the above patterns within the T1D cohort (**Supplementary Figures S17-S18**). Total β-cell area at the islet level declined significantly with disease duration (−20.0%/year, p = 0.011), whereas β-cell areas for single-cells (−6.6%/year, p = 0.28) and SEOs (−13.8%/year, p = 0.067) showed non-significant downward trends. Total α-cell islet area reciprocally increased with duration (7.7%/year, p = 0.002); α-cell areas for single-cells (1.5%/year, p = 0.34) and SEOs (3.6%/year, p = 0.10) showed non-significant upward trends. δ-cell area showed no significant duration-dependent changes across all object types: total δ-cell area (−1.5%/year, p = 0.40), single-cell area (0%/year, p = 1.0), SEO area (−0.1%/year, p = 0.93). While there was no change in total PP-cell area over time (1.6%/year, p = 0.42), PP-cell area significantly increased over time in single cells (5.6%/year, p = 0.008) and SEOs (5.2%/year, p = 0.009). Per-cell area changes with duration were generally minimal, with the notable exception of PP-cells within SEOs, where per-cell area increased modestly over T1D disease duration (4.1%/year, p = 0.017) (**Supplementary Figure S18**). Together, these analyses indicate that changes in cell type composition in EOs of T1D donors are accompanied by distinct, cell type-specific changes in per-cell area, suggesting β-cell atrophy, α-cell hyperplasia, and PP-cell hypertrophy each contributing to cytoarchitectural remodeling of T1D EOs.

### Pseudotime trajectory inference reveals region- and age-dependent islet remodeling along T1D progression

Since our donor cohort spans the full range of T1D disease durations from recent onset to long-standing disease, we leveraged this cross-sectional design to reconstruct trajectories of compositional change using pseudotime analysis (**Figure 9**, **Supplementary Figures S19-S21**). Uniform Manifold Approximation and Projection (UMAP) embedding of the islet-level cell-composition data (β-, α-, δ-, and PP-cell proportions) revealed distinct cell-type-defined regions of compositional space (**Figure 9A**). Stratifying these embeddings by diagnosis and pancreatic region uncovered T1D-associated islet clusters that were absent or rare in ND donors. Specifically, a residual β-cell cluster was present in the T1D head but largely absent from the T1D body and tail (purple arrow, **Figure 9B**), and a separate cluster was present across all three pancreatic regions of T1D donors but absent in ND. These data aligned spatially with the α-cell-enriched cluster in the global UMAP, consistent with the relocation of remaining endocrine cells into a new compositional state (red arrows, **Figure 9B**).

**Figure 9.**
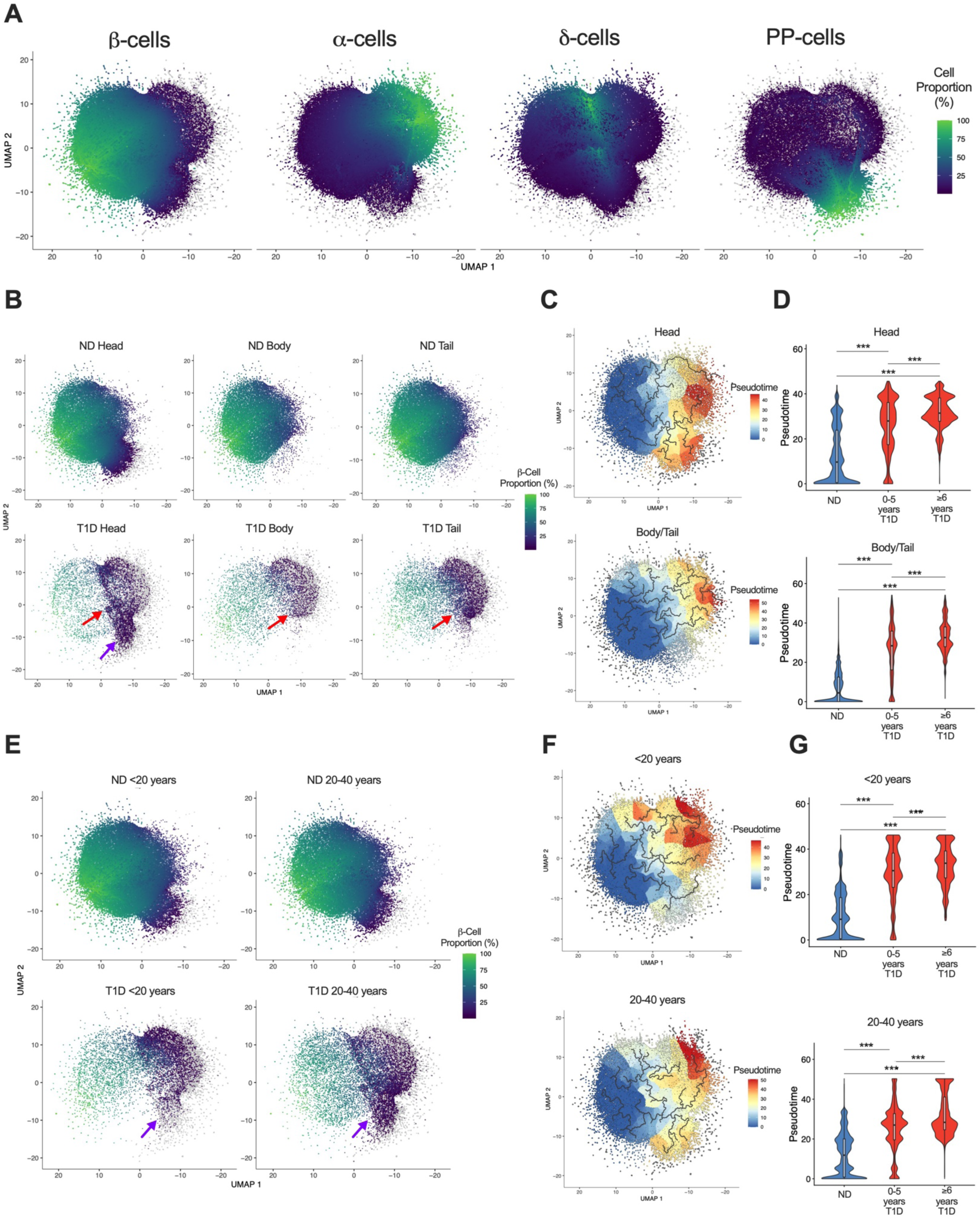
T1D disease progression inferred by islet pseudotime analysis. **(A)** UMAP visualization of cell types from ND and T1D donor islets, where each dot represents an islet, and the x and y axes present the top two principal components/UMAP dimensions. Islets are colored according to the proportions of β-cells, α-cells, 8-cells, and PP-cells, respectively. **(B)** UMAP visualization of islet β-cell composition (expressed as a proportion of the total cells within islets), split by diagnosis (ND, T1D) and pancreatic region (head, body, tail). The purple arrow indicates an islet cluster present in the pancreatic head of T1D donors but absent in ND donors. Red arrows highlight an islet cluster present in the head, body, and tail of T1D donors but absent from ND donor islets. **(C)** Trajectory inference of islets according to pancreatic region with islets from the pancreatic head (top panel) and body/tail (bottom panel) mapped onto the respective UMAPs and colored by pseudotime. **(D)** Comparison of pseudotime distributions for islets within the pancreatic head (top panel) and body/tail (bottom panel) among ND and T1D donors within 0-5 years and ≥6 years of illness onset via Wilcoxon rank-sum tests; the median value is represented by a horizontal line within the violin. Pseudotime values are significantly elevated in T1D donor islets relative to ND, and increased further with longer disease duration, in both pancreatic regions. **(E)** UMAP visualization of islet β-cell proportion, split by diagnosis (ND versus T1D) and age group (<20 years versus 20-40 years). Purple arrows denote an islet cluster present in T1D donor <20-year-old and 20-40-year-old age groups but absent in the respective ND age groups. **(F)** Trajectory inference of islets according to age group, with islets from donors aged <20 years (top panel) and 20-40 years (bottom panel) mapped onto the respective UMAPs and colored by pseudotime. **(G)** Comparison of pseudotime distributions among ND and T1D donors within 0-5 years and ≥6 years of illness onset in islets from donors aged <20 years (top panel) and 20-40 years (bottom panel) via Wilcoxon rank-sum tests; the median value is represented by a horizontal line within the violin. As in Panel D, pseudotime values are significantly elevated in T1D versus ND donors and increased further with longer disease duration in both age groups. ***p<0.001.

Pseudotime trajectory inference within these UMAPs allowed us to order islets along a putative T1D-progression axis (**Figure 9C**). Comparing pseudotime distributions between ND donors and T1D donors stratified by disease duration (recent onset, 0-5 years; longer duration, ≥6 years) revealed significant shifts toward higher pseudotime in T1D islets within both the pancreatic head and the combined body/tail regions (all p < 2 × 10^-16^; **Figure 9D**). Pseudotime values increased further with longer disease durations in both regional analyses (Head: p = 9.8 × 10^-6^; Body/Tail: p < 2 × 10^-16^), indicating that compositional remodeling continues to progress over time.

Stratifying the same analysis by donor age group (<20 years vs 20-40 years) revealed a similar pattern (**Figure 9E**). Within each age group, T1D donor islets occupied a distinct UMAP region from age-matched ND donors (purple arrows, **Figure 9E**). Pseudotime trajectory inference produced clear separations between ND and T1D in both age groups (**Figure 9F**), and pairwise Wilcoxon comparisons showed significant pseudotime elevations in T1D compared to ND, with further increases at longer disease durations within both age cohorts (all p < 2 × 10^-16^; **Figure 9G**).

Mapping α-cell, δ-cell, and PP-cell composition onto the same UMAP embeddings (**Supplementary Figure S20**) confirmed that the T1D-specific islet clusters identified for β-cells also reflected coordinated shifts in the other islet cell populations, particularly within the pancreatic head. Donor-level UMAP visualizations (**Supplementary Figures S19, S21**) showed that the broad trajectories observed in the aggregate data were apparent in individual donors as early as <1 year following T1D onset, while preserving substantial inter-donor heterogeneity. Together, these analyses point to region- and age-influenced compositional remodeling throughout T1D progression, with the pancreatic head and younger donors showing the most distinct trajectories.

## Discussion

The development and application of an integrated AI-guided imaging and statistical framework to 106 human pancreatic donors and over 2,000,000 individual islet candidates has enabled comprehensive multiplexed characterization of human islet composition in a sizeable cohort of T1D and ND donors. Using this high-throughput approach to islet analysis in the context of T1D permitted us to quantify the full complement of major endocrine cell types concurrently across large volumes of imaging data, revealing heterogeneous vulnerability to T1D-induced changes. The scale of the dataset further allowed us to systematically map how these shifts, and accompanying changes in cell number and size, depend on pancreatic region, donor age, disease duration, and islet size in sufficient detail to resolve biologically relevant interactions (see **Supplementary Figure S22** for a comprehensive summary of the study data). Overall, these results provide a comprehensive, statistically rigorous view of how T1D reshapes the cellular composition of human islets and establishes a framework for the high-throughput interrogation of metabolic tissue at scale.

The present study sits alongside a rapidly growing set of AI-assisted and 3D imaging studies of the human T1D pancreas and extends them in several respects. Most of the prior studies that resolved islet composition did so for β- and α-cells alone^28–30,49^, whereas our AI-guided pipeline simultaneously classified all four major endocrine populations across the cohort. Beyond this broader cell-type coverage, three analytical features distinguish the present work. First, rather than centering on developmental trajectory and age at onset, we used a unified Bayesian framework to dissect how endocrine composition varies as a function of clinical and anatomical covariates. Second, our cell-number-based taxonomy of single cells, SEOs, and islets, combined with per-cell-type area measurements, allowed compositional and cell-size remodeling to be tracked together across object classes. Third, we exploited the wide span of disease durations in our cohort by ordering islets along an inferred pseudotime axis of T1D progression using a trajectory-based analysis to demonstrate compositional remodeling throughout disease duration.

Together, this framework allowed us to deeply characterize T1D endocrine remodeling across cell types, anatomical location, and disease progression.

Our analyses confirm that long-standing T1D is accompanied by near-complete β-cell depletion^11,12^, where the magnitude is closely aligned with the ∼40-fold reduction in fractional β-cell area reported by Meier and colleagues in long-standing T1D^13^. Our high-throughput approach built upon these earlier studies to resolve the relationships between β-cell loss and disease covariates. Notably, the absence of significant associations between β-cell mass and either disease duration or age at onset reported in earlier manual histology cohorts^50^ does not survive in the larger sample and modeling framework adopted here. Specifically, we identified an inverse relationship between β-cell content and disease duration, particularly in the pancreatic body and tail, while age at disease onset emerged as a markedly more modest covariate. These data are consistent with previous observations that residual β-cells remain long into the course of T1D, particularly in later-onset cases^14^.

β-cell depletion in T1D is accompanied by reciprocal increases in both mean α-cell number per islet and proportion, where both increase ∼4-fold. However, this α-cell expansion does not reflect cellular hypertrophy since per-α-cell area decreases by ∼44% in islets and by 47% in SEOs in T1D. These data suggest α-cells undergo hyperplasia at the population level coupled with atrophy at the individual-cell level. This is characterized by overall more cells in islets and SEOs while the cells themselves are individually smaller. Such a pattern may therefore provide an anatomical/cellular basis for the inappropriate hyperglucagonemia and impaired glucose-sensing that is well established in T1D. Furthermore, the collapse of the β/(α+β) ratio in T1D quantifies the depth of this compositional inversion and is most pronounced in the pancreatic head, which retains the highest β/(α+β) ratio in non-diabetic states.

Notably, across our ND cohort, we discovered that α-cells comprised only ∼10-25% of islet cells (mean ∼14%), a far more β-cell-skewed composition than the 30-40% α-cell fraction widely accepted for human islets^51^, and one more reminiscent of rodent islets^52^. This discrepancy may be accounted for by our approach which relies on *in situ* sampling that spans the entire organ, including the PP-cell-rich lobe of the posterior head, where glucagon+ (α)-cells are the least frequent endocrine cell type^53^. Consistent with this, α-cell proportions were lowest in the head (∼9%) and higher in the body and tail (∼17%). Additionally, the canonical 30-40% figure derives largely from studies of isolated human islets, which do not faithfully preserve native islet organization. Indeed, the β-cell core/α-cell mantle arrangement found in intact (and T2D) tissue is lost in cultured isolated islets^54^, and the abundant small, β-cell-rich EOs typically do not survive the isolation process^30^. We therefore posit that isolated-islet preparations over-represent larger, comparatively α-cell-rich islets and overestimate the α-cell fraction of the intact pancreas. By resolving the full size and regional distribution of EOs *in situ*, our data likely provide a more representative estimate of native human islet composition.

δ-cells have been traditionally under-examined in T1D. Our data showed that δ-cells remained largely resilient, with preservation of δ-cell proportional content and per-cell area within islets along with only modest changes in δ-cell numbers per islet. Although δ-cells are thus quantitatively preserved, earlier work indicated that their spatial organization is altered. In a manual analysis of long-standing T1D, Tegehall and colleagues reported that δ-cells shift toward the islet periphery, that extra-islet single δ-cells become more than 3-fold more abundant, and that approximately twice as many α-cells come to lie directly adjacent to δ-cells^55^. Taken together, these findings suggest that δ-cells are affected in T1D primarily through cytoarchitectural reorganization rather than through changes in their number or islet proportion.

PP-cells have similarly been rarely examined in the context of T1D. In contrast to δ-cells, we found that PP-cells exhibited a striking hypertrophic response in T1D. Although overall PP-cell numbers per islet were largely preserved in T1D, individual PP-cells approximately doubled in size within SEOs and islets as well as extra-islet singlets. This yielded an ∼2-3-fold expansion in total PP-cell area without an equivalent increase in cell count. To our knowledge, PP-cell hypertrophy has not previously been reported in T1D. Earlier work either did not directly examine PP-cells^28–30,49^ or, where PP-cells were resolved, characterized their abundance and head-predominant distribution^31^ rather than their size. Because the phenotype combines preserved cell number with increased per-cell size, it would otherwise be missed by composition- or hormone-area-based metrics alone. The biological significance of PP-cell hypertrophy in T1D remains unclear and warrants further characterization. Although PP-cell-enriched islets were previously described in classic immunocytochemical studies of long-duration juvenile diabetes^12^, those observations reflected PP-cell abundance and distribution rather than the per-cell hypertrophy reported here. Together, these findings point to significant T1D-induced reshaping of the islet microenvironment across all four major endocrine cell types both at the population and individual cell levels.

Increasing evidence suggests that a substantial component of apparent β-cell loss in long-duration T1D reflects altered β-cell identity which encompasses both dedifferentiation and transdifferentiation rather than outright cellular destruction^22,56,57^. Dedifferentiating β-cells downregulate mature β-cell transcription factors, re-express progenitor genes, and can either lose hormone expression entirely or acquire glucagon expression. As a result, these cells are viewed as α-cells by hormone-based imaging despite their β-cell origin^56^. Lineage tracing in rodent diabetes models has further documented β-to-α-cell transdifferentiation as a discrete process^58,59^, and the reverse process, α-to-β-cell conversion following extreme β-cell ablation, has been demonstrated experimentally and is now under active investigation as a regenerative strategy for treating diabetes^59–61^. While 2D imaging of fixed endocrine markers cannot dissect the relative contributions of dedifferentiation, transdifferentiation, and *bona fide* α-cell proliferation to the compositional changes we observed in surviving T1D islets, the conservation of total cell counts places a quantitative constraint on any combination of these processes. Our data suggest that these processes must collectively preserve total cell numbers within surviving islets while reshaping their content from predominantly β-cell to predominantly α-cell identity.

A major methodological advance enabled by our AI-guided imaging analysis framework is the ability to study disease-induced cell type-specific alterations using both cell numbers per EO along with changes to per-cell area. We found that these two measures were non-interchangeable in T1D. Whereas β-cell loss encompasses reductions in both number and size within islets, α-cell expansion demonstrates increased cell numbers but decreased per-cell size. Together, our ability to examine these changes across large numbers of donors and EOs has shown that the conventional reliance on hormone-positive area as a proxy for cell abundance potentially conflates fundamentally distinct phenomena such as cellular hyperplasia, hypertrophy, and atrophy that arise from different underlying mechanisms and likely carry different functional consequences.

We found that a substantial fraction of insulin-expressing EOs in ND donors lack α-cells which recapitulates findings by Lehrstrand and colleagues^28^. Our data showed that 22.6% of insulin-expressing islets and 60.7% of all insulin-expressing EOs lacked α-cells, in line with the prior study’s estimates of ∼50%. Moreover, at the islet level, our 2D imaging data showed that α-cell-absent islets contributed 14.3% of total non-diabetic β-cell mass, matching closely with the ∼16% of total islet volume reported by Lehrstrand using a 3D imaging approach. Notably, the per-donor distribution of α-cell-less islet fractions in ND donors was well-behaved and symmetric (median 20.1%, IQR 12.7-26.8%), indicating that the observed pattern reflects a consistent anatomical feature rather than the contribution of outlier donors. More broadly, our data may reconcile two seemingly opposing observations. Compositionally, surviving β-cells became concentrated within larger islets, leaving small extra-islet EOs depleted of β-cells. This is consistent with the marked depletion of small extra-islet β-cells reported by Murrall and colleagues^30^ and the loss of small insulin-positive/glucagon-negative islets described by Rippa and colleagues using 3D imaging of T1D pancreata^29^. At the same time, the β-cells that persisted as single extra-islet cells were the least atrophied by area, suggesting a relative sparing of the extra-islet compartment. This agrees with long-standing observations that residual β-cells in chronic T1D persist predominantly as scattered extra-islet cells and small clusters^14^, and with a recent 3D imaging study of a whole pancreas from a late-onset T1D donor, in which most residual β-cells were similarly extra-islet and relatively enriched in the pancreatic head^49^. We posit that these patterns, compositional depletion of small β-cell-rich objects alongside relative preservation of the β-cells that survive outside islets, may reflect the “push-pull” dynamics by which the residual endocrine pancreas is reshaped over time.

Our framework uncovered substantial heterogeneity in T1D islet remodeling along multiple dimensions. The pancreatic head harbored the highest β/(α+β) ratio and the most pronounced absolute β-cell content in ND donors, while the body and tail exhibited the steepest declines in insulin content with disease duration. Pseudotime trajectory analysis identified region- and age-dependent paths of compositional change across the T1D duration spectrum, with the pancreatic head and younger donors exhibiting the most distinct trajectories. While these findings are in line with earlier reports of regional heterogeneity in islet distribution^23–26^, the cohort and analytical scale of the present work allow them to be quantified as part of a coherent multidimensional model rather than as isolated observations. These trajectories suggest that the progression of T1D across time is not uniform but instead is heterogenous, demonstrating islet remodeling that follows region- and disease stage-specific paths.

**Limitations of study.** We acknowledge that our four-marker panel quantifies the principal endocrine populations but does not account for immune-cell infiltration, vascular changes, or stromal remodeling. Highly multiplexed imaging approaches such as imaging mass cytometry^27,62^ and multiplexed immunohistochemistry^31^ will be required to integrate the compositional shifts we describe with the immunological events that drive them. Second, we restricted the present analysis to donors with clinically established T1D together with ND controls. Therefore, donors with islet autoantibody positivity (AAb+), who are critical for understanding the preclinical phase and have featured prominently in recent work^29–31^, were excluded due to the limited number of AAb+ donors available; future work will specifically focus on these AAb+ donors as an extension of the present pipeline. Third, the cohort was restricted to donors ≤40 years of age to limit confounding from age-related islet changes. We therefore acknowledge that trajectories in older T1D donors may follow distinct paths. Fourth, our imaging is 2D and cannot directly resolve 3D islet architecture, which recent 3D imaging studies^28,29,49^ have shown to be biologically important. Nevertheless, our cohort and islet-level statistical power substantially exceed what current 3D approaches can achieve. Finally, our analyses are cross-sectional; while pseudotime inference orders islets along a putative progression axis, true longitudinal trajectories cannot be directly observed in postmortem tissue.

**Conclusions.** By quantifying the complete endocrine compartment of more than two million islet candidates across 106 donors with statistical resolution that integrates diagnosis, region, sex, age, disease duration, and islet size, we provide a comprehensive view of how T1D reshapes the human pancreatic islet. Despite a coordinated rearrangement of islet cell composition, including profound β-cell loss accompanied by substantial increases in α-cells, overall cell numbers remain largely unchanged in surviving islets throughout disease duration. Alongside these changes, there are distinct morphological alterations at cellular and regional levels among the cell types. Overall, the AI-guided imaging and statistical framework introduced here provides a generalizable foundation for systematic interrogation of tissue and cellular imaging to further refine our understanding of the cellular events across the course of T1D.

## Acknowledgements

We are grateful for discussions and technical assistance provided by Drs. Irina Kusmartseva, Mark A. Atkinson, Helmut Hiller, Kacie Geelhoed, Rafaela Vieira, Maria Beery, Maciej Zerkowski, Spencer Revill, and Sara Amirault. This research was performed with the support of the Network for Pancreatic Organ donors with Diabetes (nPOD; RRID:SCR_014641), a collaborative type 1 diabetes research project supported by Breakthrough T1D and The Leona M. & Harry B. Helmsley Charitable Trust (Grant#3-SRA-2023-1417-S-B). The content and views expressed are the responsibility of the authors and do not necessarily reflect the official view of nPOD. Organ Procurement Organizations (OPO) partnering with nPOD to provide research resources are listed at https://npod.org/for-partners/npod-partners/. We express our deepest gratitude to the families of the organ donors for the gift of tissues. Additional support for this study was provided by the National Institutes of Health (R01DK124219 to ZF; R36DA057972 to JK; T32MH019986 to SJM; T32GM133353 to CW and JK, R35GM159862 to SL, R01DK135268 to GAR, R01DK139630- 01A1 to GAR), the Department of Defense (PR210207 to ZF), a Canada Research Chair (Tier-1, CRC-2024-00032 to GAR), a CIHR-JDRF Team grant (CIHR-IRSC TDP-186358 to GAR, JDRF 4-SRA-2023-1182-S-N to GAR), CRCHUM start-up funds (to GAR), and Innovation Canada John R. Evans Leader Awards (CFI 42649, 46539 to GAR). This research was also supported in part by the University of Pittsburgh Center for Research Computing through the resources provided. Specifically, this work used the HTC cluster, which is supported by NIH award number S10OD028483.

## Author Contributions

CW, TBT, HT, PNJ, and ZF conceived the project. CW performed training and validation of AI algorithms on human donor pancreatic images. CW, JY, AMG, HT, PNJ, and JK established the image analysis workflow. CW, TBT, HT, AMG, SJM, JY, LLP, JJL, HCC, AFC, HMS, LMW, GCT, and SL performed the data processing analyses. HT, LLP, TBT, SJM, and ZF performed the statistical analyses. CW, TBT, HT, LLP, and ZF wrote the manuscript with contributions from EMR and GAR. As the guarantor of this work, ZF had complete access to all the data and takes full responsibility for the integrity of the data and accuracy of analysis.

