## Supplementary Materials for "AI-guided analysis of human pancreatic islet sociology reveals distinct cell compositional changes in type 1 diabetes"

**Supplementary Figures S1-S22**

**Supplementary Table S1. Donor demographic information**
**Supplementary Table S2. Statistics for main and supplementary figures**

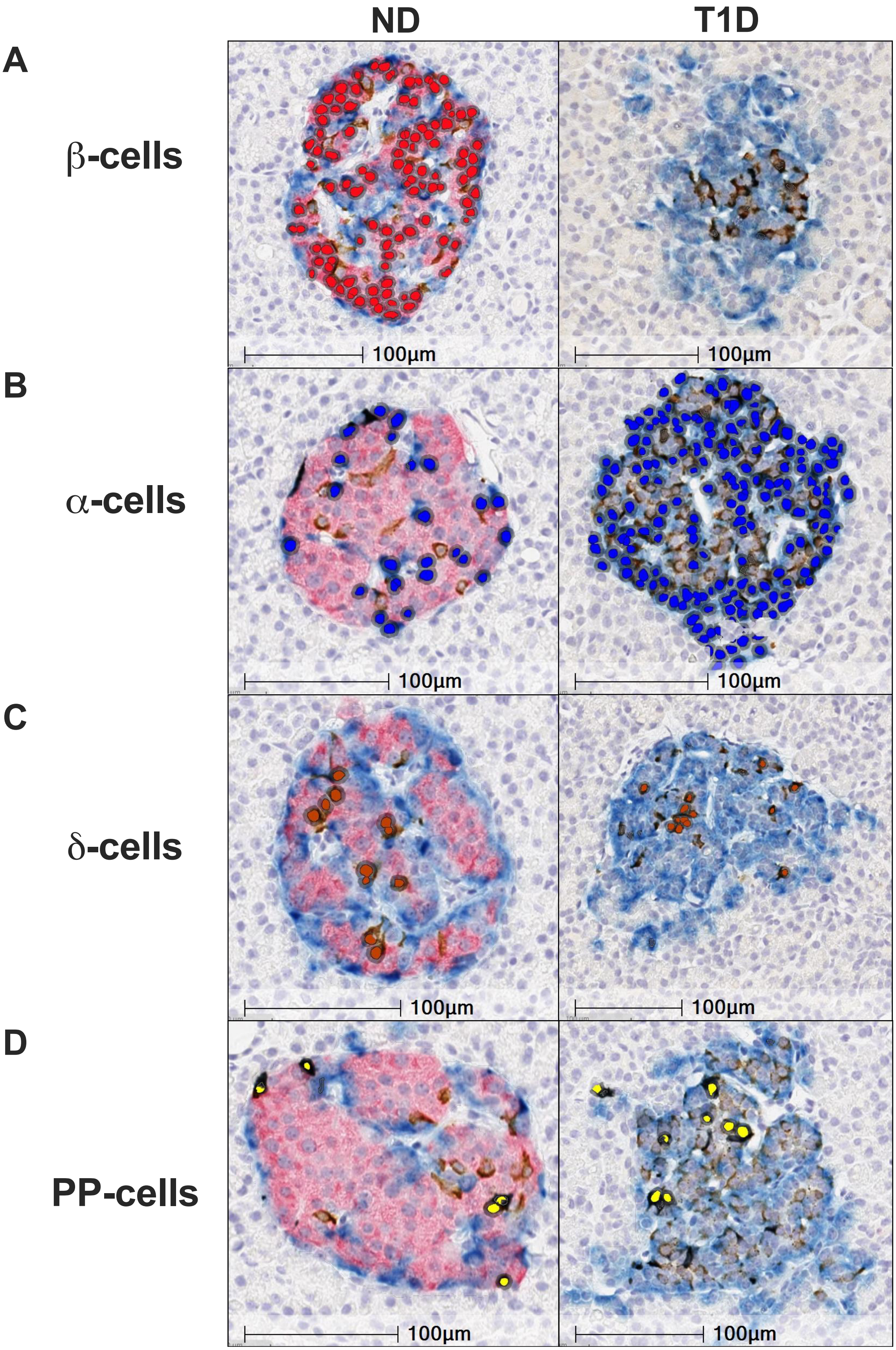
ND
T1D
A
β-cells
B
α-cells
C
δ-cells
D
PP-cells
100µm
100µm
100µm
100µm
100µm
100µm
100µm
100µm

**Supplementary Figure S1. AI-guided annotation of distinct islet cell types in non-diabetic and type 1 diabetes donor pancreatic tissue. (A-D)** Representative images of islets within pancreatic tissue from non-diabetic (ND) and type 1 diabetic (T1D) donors featuring annotated nuclei from different islet cell types via trained AI nuclei classifiers: **(A)** β-cells (in red); **(B)** α-cells (in blue); **(C)** δ-cells (in brown); and **(D)** PP-cells (in yellow). Scale bars=100 μm.

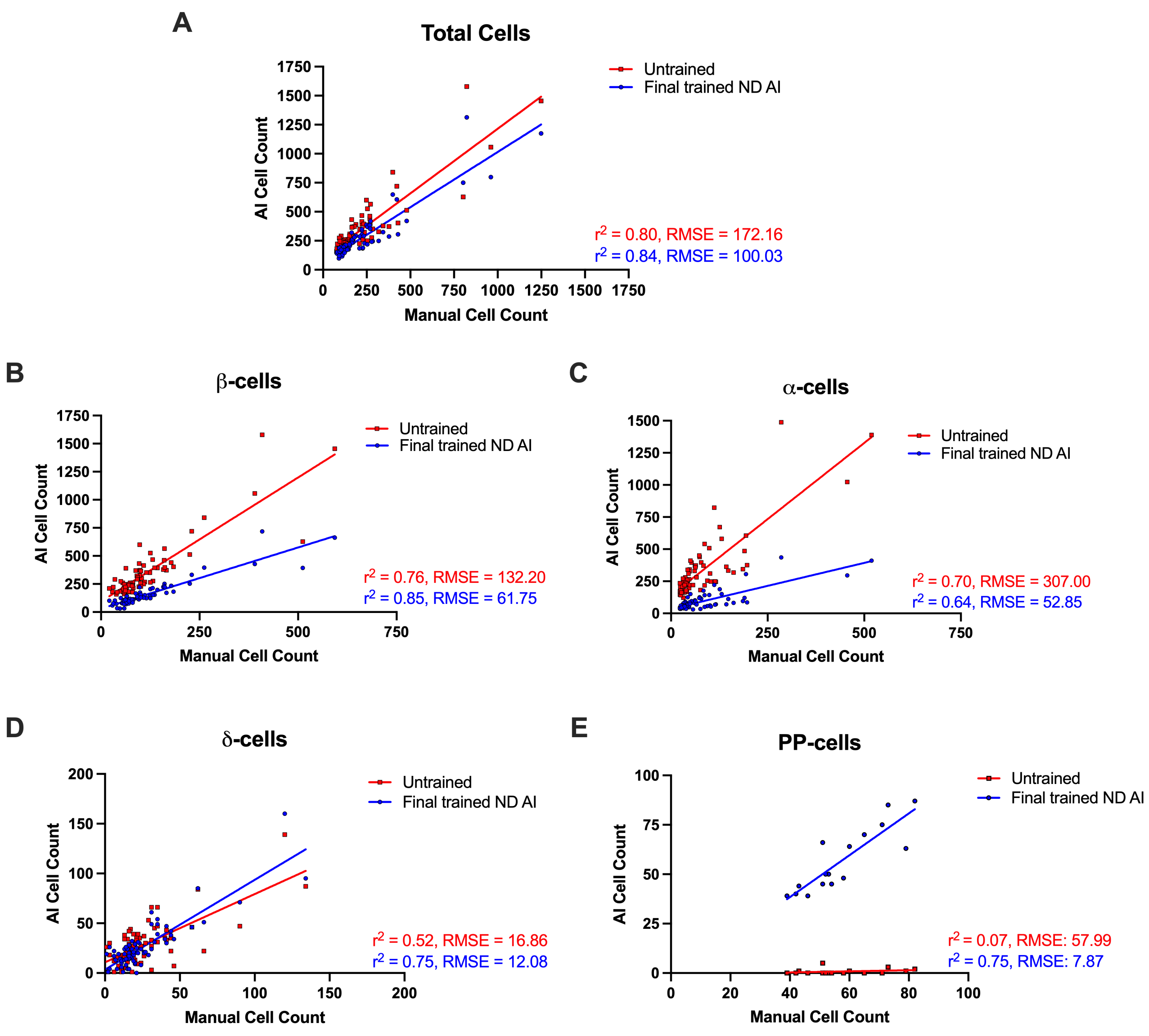


**Supplementary Figure S2. Determination of trained versus untrained AI nuclei classifier accuracy in islet annotation within ND donor pancreatic tissue. (A-E)** The accuracies of untrained (red squares) versus trained (blue circles) AI-guided nuclei classifiers in annotating different islet cell types (β-cells, α-cells, δ-cells, PP-cells) are determined by comparisons to manually annotated islets in pancreatic tissue from ND donors. Relationships between untrained and trained AI-guided versus manual approaches are graphed as correlation plots via linear regression, with accompanying $r^2$ and root mean squared error (RMSE) values for numbers of: **(A)** total cells per islet, comprising all islet cell types combined (termed 'Total Cells'); **(B)** β-cells per islet; **(C)** α-cells per islet; **(D)** δ-cells per islet; and **(E)** PP-cells per islet. Each point represents an individual islet.

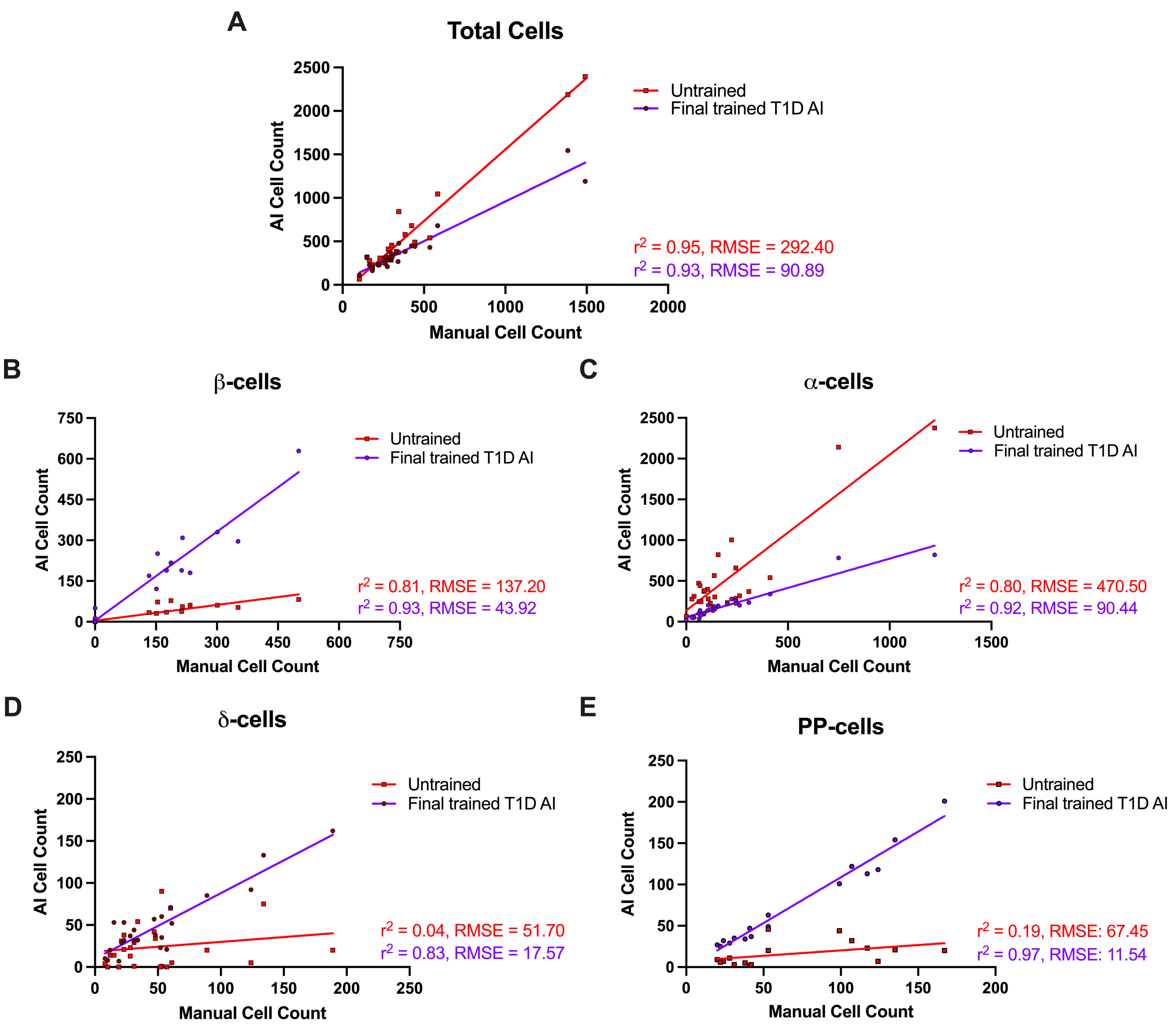


**Supplementary Figure S3. Determination of trained versus untrained AI nuclei classifier accuracy in islet annotation within T1D donor islets. (A-E)** The accuracies of untrained (red squares) versus trained (purple circles) AI-guided nuclei classifiers in annotating different islet cell types (β-cells, α-cells, δ-cells, PP-cells) are determined by comparisons to manually annotated islets in pancreatic tissue from T1D donors. Relationships between untrained and trained AI-guided versus manual approaches are graphed as correlation plots via linear regression, with accompanying $r^2$ and RMSE values for numbers of: **(A)** total cells per islet, comprising all islet cell types combined (termed 'Total Cells'); **(B)** β-cells per islet; **(C)** α-cells per islet; **(D)** δ-cells per islet; and **(E)** PP-cells per islet. Each point represents an individual islet.

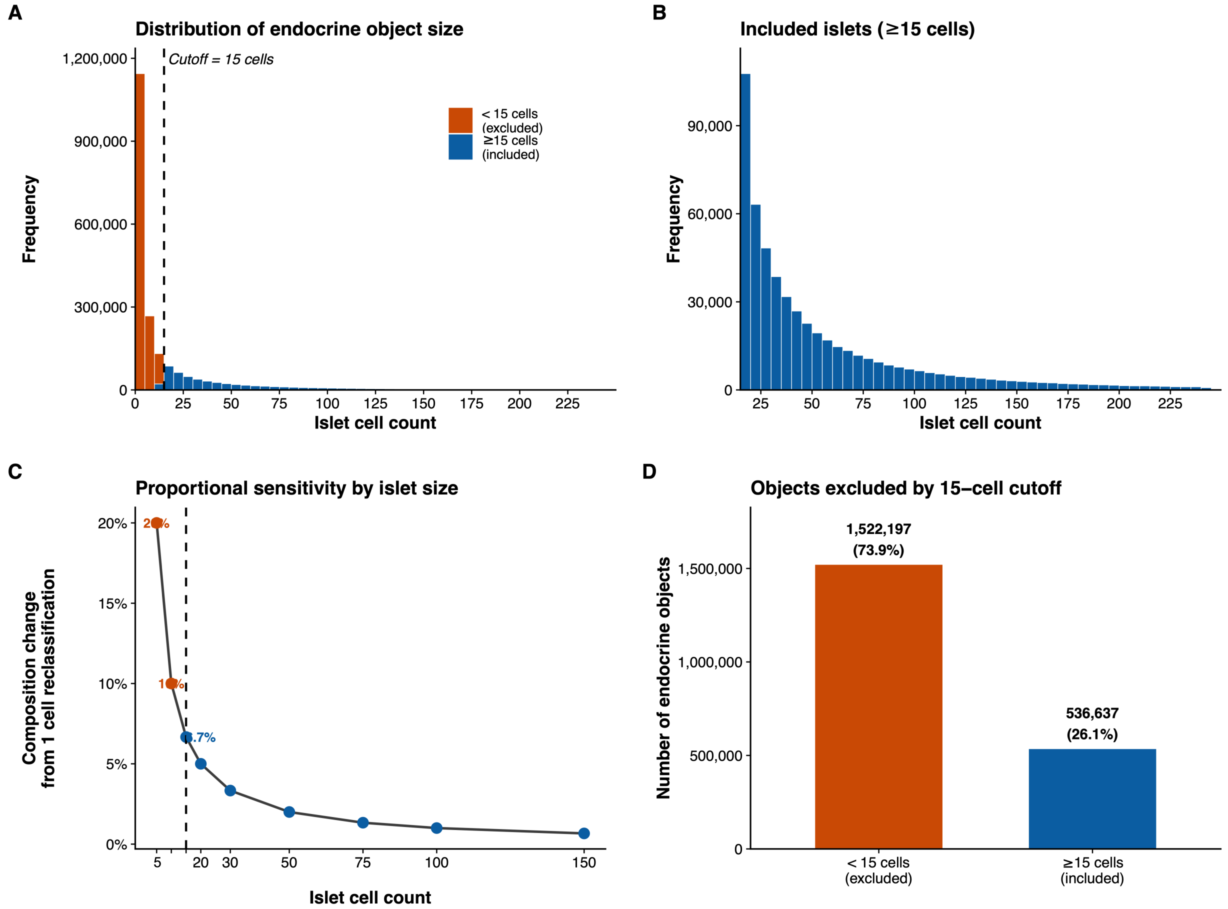


**Supplementary Figure S4. Establishment of islet inclusion criteria according to islet cell thresholding. (A)** Distribution of endocrine object (EO) size (cell count per object) across all donors, with objects containing <15 islet cells (orange) and ≥15 islet cells (blue) indicated. Dashed line denotes the 15-cell threshold. **(B)** Size distribution of included islets (≥15 cells) after applying the 15-cell threshold. **(C)** Proportional sensitivity analysis demonstrating the impact of a single cell reclassification on islet composition as a function of EO size. **(D)** Summary of excluded versus included EOs according to the 15-cell threshold.

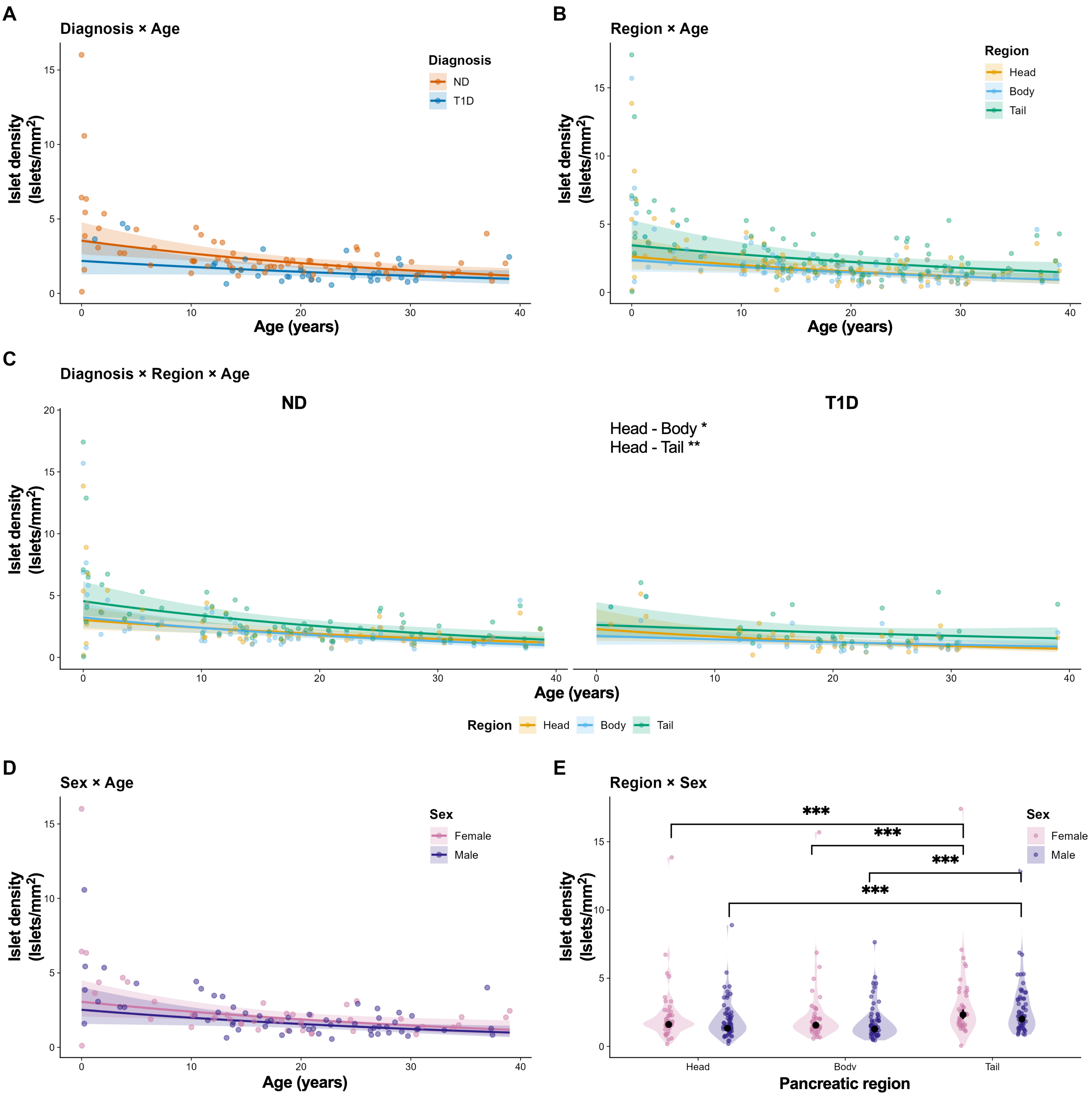


**Supplementary Figure S5. Analysis of interactions between donor age, diagnosis, sex, and pancreatic region for islet density. (A)** Predicted islet density as a function of age, stratified by diagnosis. The diagnosis × age interaction is not significant ($p = 0.527$), indicating that the rate of age-related density decline does not differ between ND and T1D donors. Points represent individual donor means. **(B)** Predicted islet density as a function of age, stratified by pancreatic region and averaged across diagnosis and sex. There are no significant region × age interactions for both the pancreatic body ($p = 0.533$) and tail ($p = 0.142$) relative to the head; there were no significant region × age interactions between body versus tail ($p = 0.729$). **(C)** Three-way diagnosis × region × age interaction. Panels show model-predicted density trajectories for ND and T1D donors across age within each pancreatic region. The ND vs T1D slope difference is not significant within any individual region (all $p > 0.05$). Among T1D donors, the age-density slope in the pancreatic head is significantly steeper compared to the body ($p = 0.024$) and tail ($p = 0.001$), indicating that age-related islet density loss in T1D is most pronounced in the pancreatic head. No significant regional slope differences are observed in ND donors (all $p > 0.05$). **(D)** Predicted islet density as a function of age, stratified by sex. The sex × age interaction is not significant ($p = 0.953$). **(E)** Islet density by pancreatic region stratified by sex. There are no significant region × sex interactions for the body (males: $p = 0.307$; females: $p = 0.661$) relative to the head. There are significant interactions relative to head (males: $p < 0.0001$; females: $p < 0.0001$), and body versus tail (males: $p < 0.0001$; females: $p < 0.0001$). Colored points represent individual donor means; black points with error bars indicate model-estimated marginal means with 95% confidence intervals. Prediction lines in Panels A-D are population-level estimates from the GLMM at reference levels of non-displayed covariates.

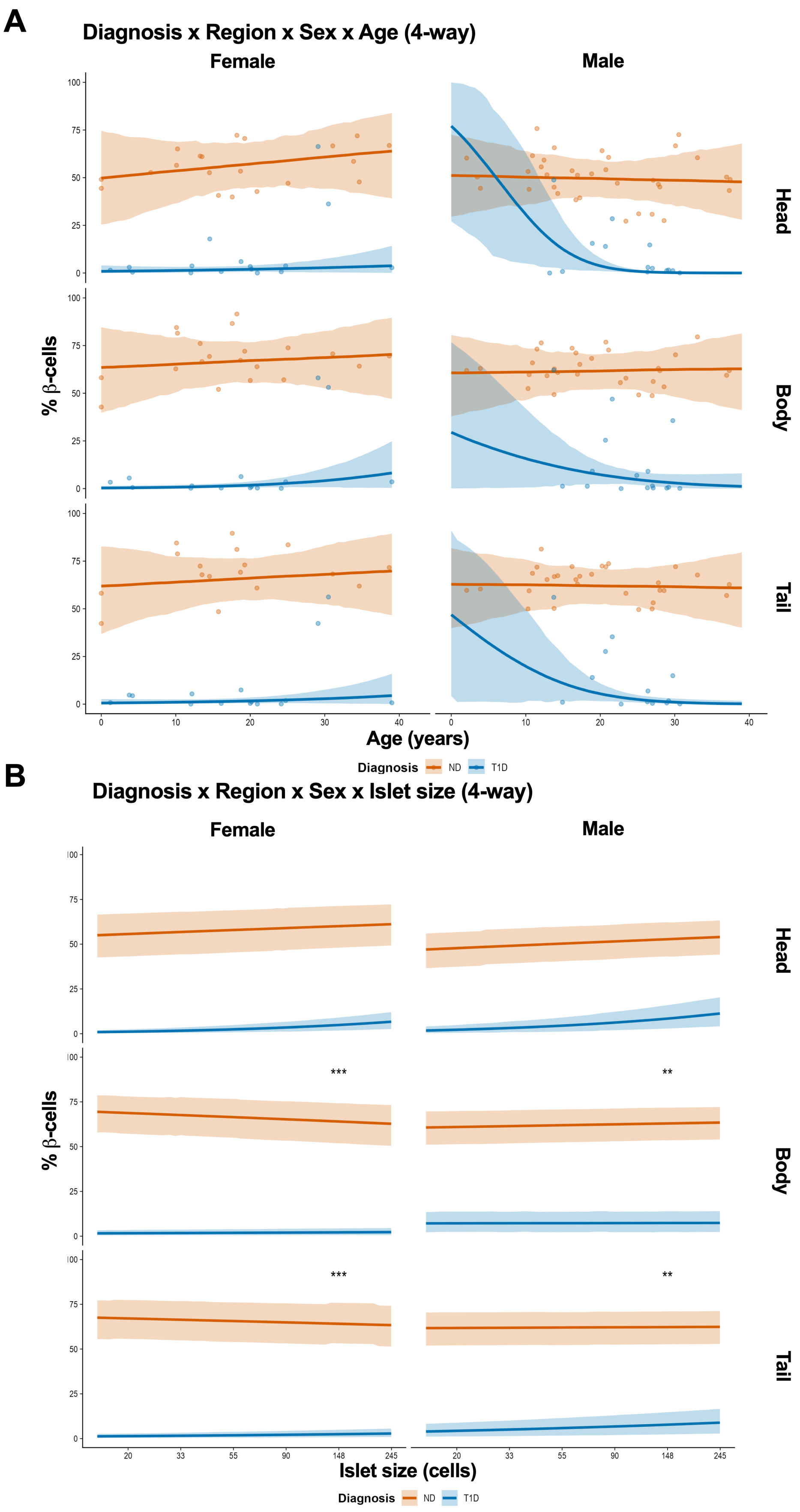
A
Diagnosis x Region x Sex x Age (4-way)
Female
Male
Head
Body
Tail
% β-cells
0
25
50
75
100
0
10
20
30
40
Age (years)
Diagnosis
ND
T1D
B
Diagnosis x Region x Sex x Islet size (4-way)
Female
Male
Head
Body
Tail
% β-cells
***
**
20
33
55
90
148
245
Islet size (cells)
Diagnosis
ND
T1D

**Supplementary Figure S6. Four-way interaction analyses of β-cell proportion in ND versus T1D donors across pancreatic regions, age, sex, and islet size. (A)** Percentage of β-cells as a function of donor age (years), stratified by diagnosis (ND, orange; T1D, blue), pancreatic region (head, body, tail), and sex. Each point represents an individual donor observation. Fitted lines and 95% confidence bands are derived from a Bayesian ordered beta regression model with diagnosis × region × sex × age interactions and donor as a random effect (ND: n = 17-35; T1D: n = 13-16). In ND donors, β-cell proportion remains relatively stable across age in both sexes and all pancreatic regions. In T1D donors, β-cell proportion is near-zero across most ages and regions, consistent with extensive β-cell loss. **(B)** Percentage of β-cells as a function of islet size (number of cells per islet), stratified by diagnosis (ND, orange; T1D, blue), pancreatic region (head, body, tail), and sex. Fitted lines and 95% confidence bands are derived from a Bayesian ordered beta regression model with diagnosis × region × sex × islet size interactions and donor as a random effect (ND: n = 17-35; T1D: n = 13-16). There are significant diagnosis × islet size interactions for both ND and T1D female and male donors. All estimates are posterior medians with 95% highest posterior density intervals. *Probability of direction (pd) > 0.975, **pd > 0.995, ***pd > 0.999.

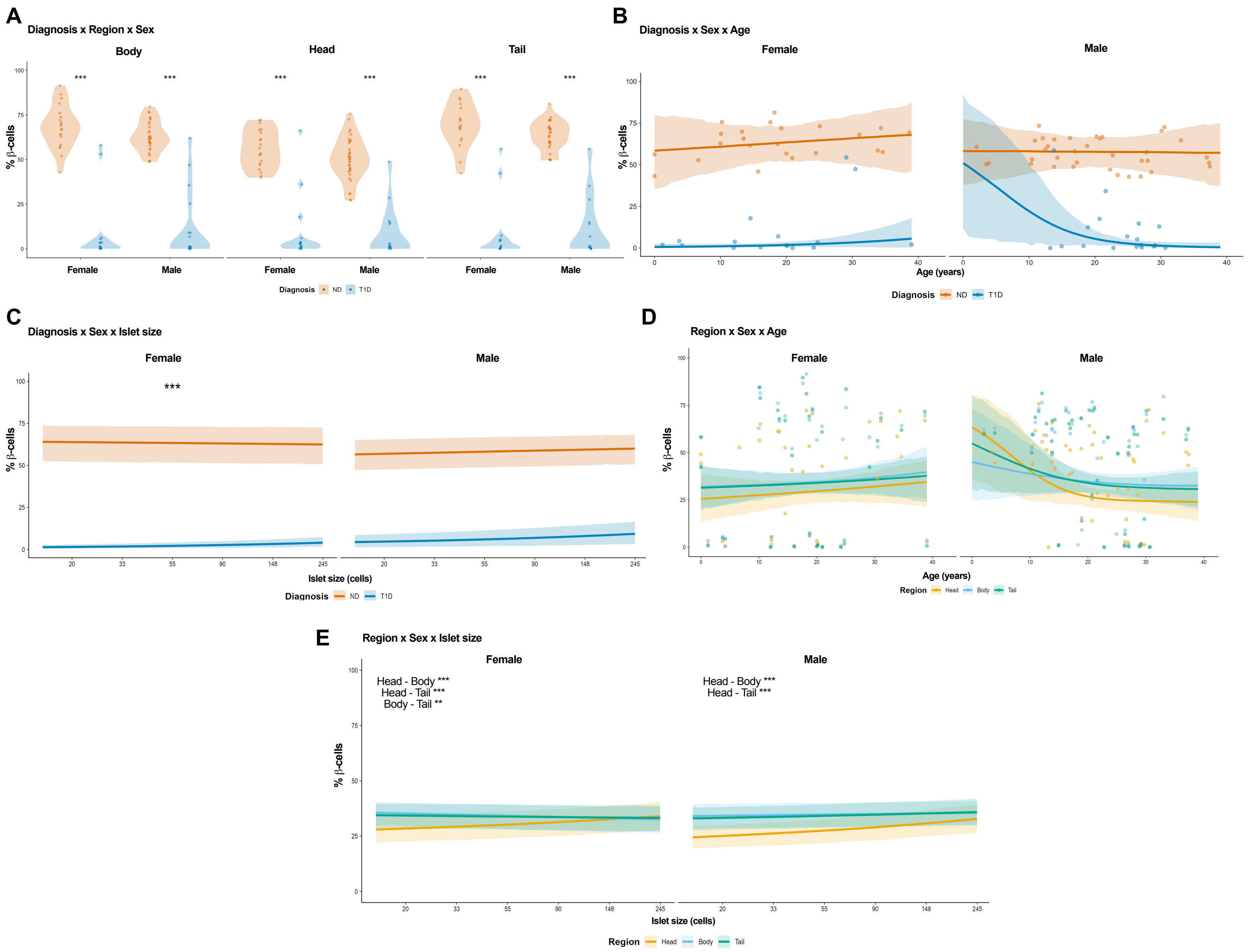


**Supplementary Figure S7. Three-way interaction analyses reveal heterogeneity in β-cell proportion.** All panels are derived from a 4-way Bayesian ordered beta regression model with diagnosis × region × sex × age and diagnosis × region × sex × islet size interactions and donor as a random effect (ND: n = 17-35; T1D: n = 13-16). **(A)** β-cell proportion (%) by pancreatic region (body, head, tail) and sex, stratified by diagnosis. Violin plots show the distribution of individual donor observations; points represent individual donors. β-cell proportion is significantly reduced in T1D compared to ND in all region × sex comparisons. The ND-T1D difference is largest in the pancreatic body of female donors (estimate = 0.652, 95% HPD [0.536, 0.751], pd > 0.999) and smallest in the pancreatic head of males (estimate = 0.455, 95% HPD [0.349, 0.552], pd > 0.999).

**(B)** β-cell proportion as a function of donor age (years), stratified by diagnosis and sex and collapsed across pancreatic region. Each point represents an individual donor observation. Fitted lines with 95% confidence bands are shown. β-cell proportion is stable across age in both sexes for ND and T1D donors. **(C)** β-cell proportion as a function of islet size (number of cells per islet), stratified by diagnosis and sex and collapsed across pancreatic region. Fitted lines with 95% confidence bands are shown. The diagnosis × islet size interaction is significant in females (estimate = 0.013, 95% HPD [0.007, 0.020], pd > 0.999). **(D)** β-cell proportion as a function of donor age, stratified by pancreatic region and sex, and collapsed across diagnosis. Each point represents an individual donor observation. Fitted lines with 95% confidence bands are shown. β-cell proportion is generally stable across age in all regions for both sexes. **(E)** β-cell proportion as a function of islet size, stratified by pancreatic region and sex, collapsed across diagnosis. Fitted lines with 95% confidence bands are shown. In females, significant region differences in islet size slopes are observed for all pairwise comparisons. (Head - Body: estimate = 0.029, 95% HPD [0.023, 0.036], pd > 0.999; Head - Tail: estimate = 0.023, 95% HPD [0.018, 0.029], pd > 0.999; Body - Tail: estimate = -0.006, 95% HPD [-0.010, -0.002], pd = 0.9995). In males, the head differs significantly from the body (Head - Body: estimate = 0.020, 95% HPD [0.011, 0.032], pd > 0.999) and tail (Head - Tail: estimate = 0.016, 95% HPD [0.011, 0.023], pd > 0.999), while the body and tail do not differ significantly. All estimates are posterior medians with 95% highest posterior density (HPD) intervals. *Probability of direction (pd) > 0.975, **pd > 0.995, ***pd > 0.999.

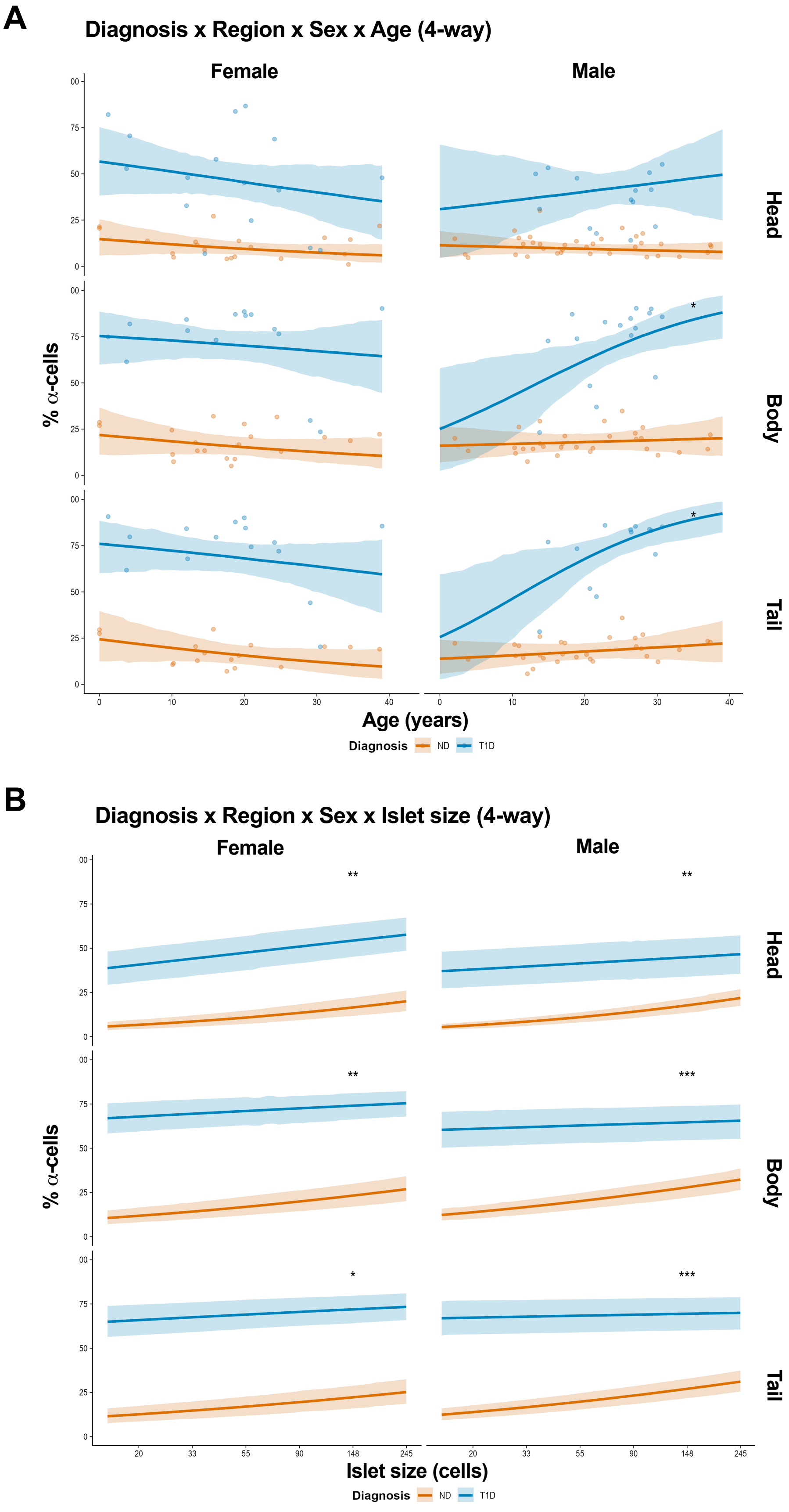
A
Diagnosis x Region x Sex x Age (4-way)
Female
Male
Head
Body
Tail
% α-cells
Age (years)
Diagnosis
ND
T1D
B
Diagnosis x Region x Sex x Islet size (4-way)
Female
Male
Head
Body
Tail
% α-cells
Islet size (cells)
Diagnosis
ND
T1D

**Supplementary Figure S8. Four-way interaction analyses of α-cell proportion in ND versus T1D donors across pancreatic regions, age, sex, and islet size. (A)** Percentage of α-cells as a function of donor age (years), stratified by diagnosis (ND, orange; T1D, blue), pancreatic region (head, body, tail), and sex. Each point represents an individual donor observation. Fitted lines and 95% confidence bands are derived from a Bayesian ordered beta regression model with diagnosis × region × sex × age interactions and donor as a random effect (ND: n = 17-35; T1D: n = 13-16). In ND donors, α-cell proportion is relatively stable across age in all pancreatic regions and both sexes. In T1D donors, α-cell proportion is markedly elevated in all pancreatic regions and both sexes across age. **(B)** Percentage of α-cells as a function of islet size (number of cells per islet), stratified by diagnosis (ND, orange; T1D, blue), pancreatic region (head, body, tail), and sex. Fitted lines and 95% confidence bands are derived from a Bayesian ordered beta regression model with diagnosis × region × sex × islet size interactions and donor as a random effect (ND: n = 17-35; T1D: n = 13-16). There are significant diagnosis × islet size interactions for both ND and T1D female and male donors. All estimates are posterior medians with 95% highest posterior density (HPD) intervals. *Probability of direction (pd) > 0.975, **pd > 0.995, ***pd > 0.999.

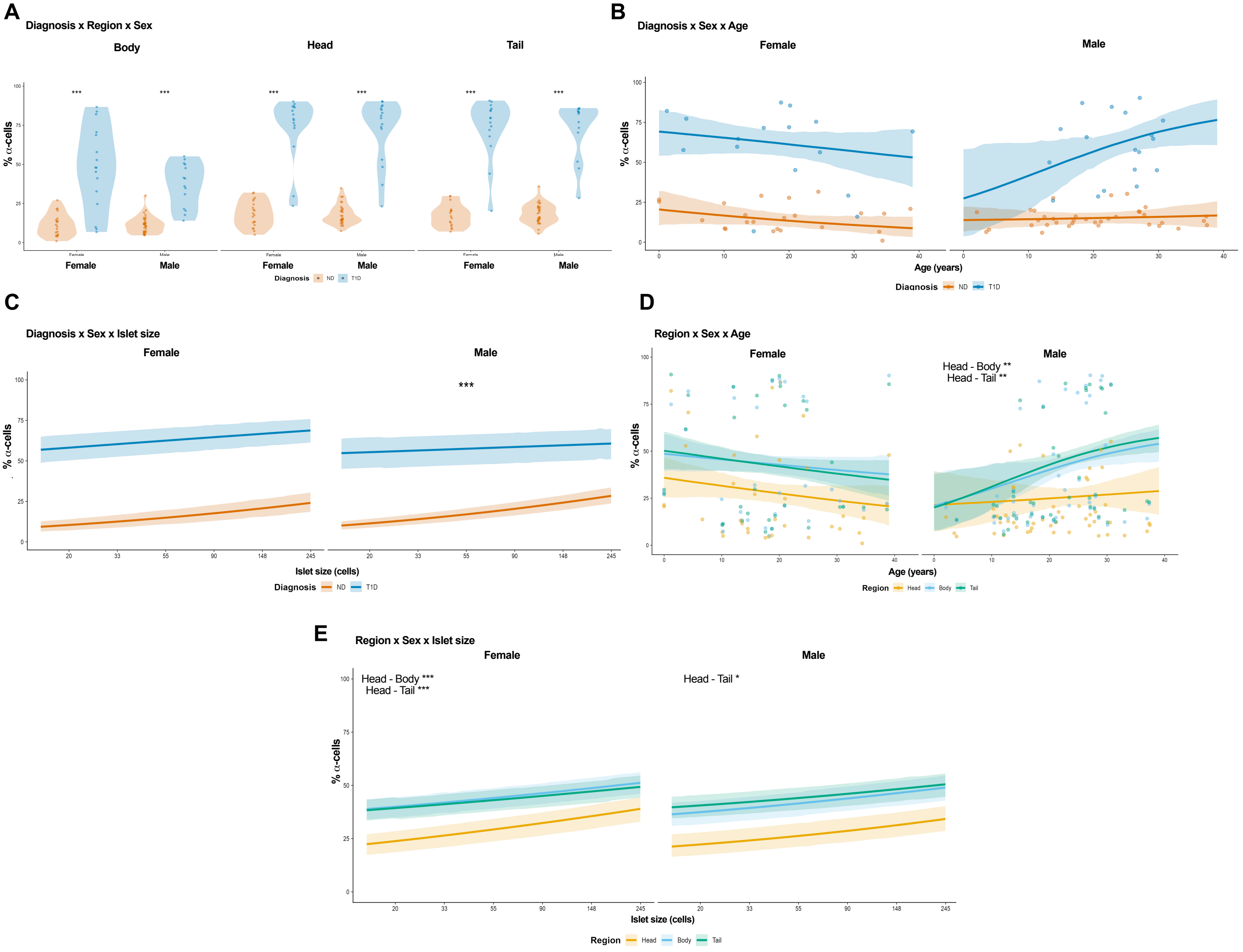


**Supplementary Figure S9. Three-way interaction analyses reveal heterogeneity in α-cell proportion.** All panels are derived from a 4-way Bayesian ordered beta regression model with diagnosis × region × sex × age and diagnosis × region × sex × islet size interactions and donor as a random effect (ND: n = 17-35; T1D: n = 13-16). **(A)** α-cell proportion (%) by pancreatic region (body, head, tail) and sex, stratified by diagnosis. Violin plots show the distribution of individual donor observations; points represent individual donors. α-cell proportion is significantly increased in T1D compared to ND in all region × sex comparisons. **(B)** α-cell proportion as a function of donor age (years), stratified by diagnosis and sex, collapsed across pancreatic region. Each point represents an individual donor observation. Fitted lines with 95% confidence bands are shown. Differences in α-cell proportion between T1D and ND donors in both sexes as a function of age is not significant. **(C)** α-cell proportion as a function of islet size (number of cells per islet), stratified by diagnosis and sex and collapsed across pancreatic region. Fitted lines with 95% confidence bands are shown. The diagnosis × islet size interaction is significant in males. **(D)** α-cell proportion as a function of donor age, stratified by pancreatic region and sex, and collapsed across diagnosis. Each point represents an individual donor observation. Fitted lines with 95% confidence bands are shown. In males, there are significant differences between the trajectories of changes to islet α-cell proportion in the head versus body (estimate = -0.0078, 95% HPD [-0.0137, -0.0026], pd = 0.998) as well as in the head versus tail (estimate = -0.0089, 95% HPD [-0.0146, -0.0036], pd = 0.999). **(E)** α-cell proportion as a function of islet size, stratified by pancreatic region and sex, collapsed across diagnosis. Fitted lines with 95% confidence bands are shown. In females, there are significant region-specific differences in the change in α-cell proportion as islet size increases for the head versus body (Head - Body = 0.014, 95% HPD [0.007, 0.021], pd > 0.999) and head versus tail (Head - Tail = 0.018, 95% HPD [0.012, 0.025], pd > 0.999). Males also show differences in α-cell proportion as islet size increases for the head versus tail (Head - Tail = 0.006, 95% HPD [0.0002, 0.012], pd = 0.979). All estimates are posterior medians with 95% highest posterior density (HPD) intervals. *Probability of direction (pd) > 0.975, **pd > 0.995, ***pd > 0.999.

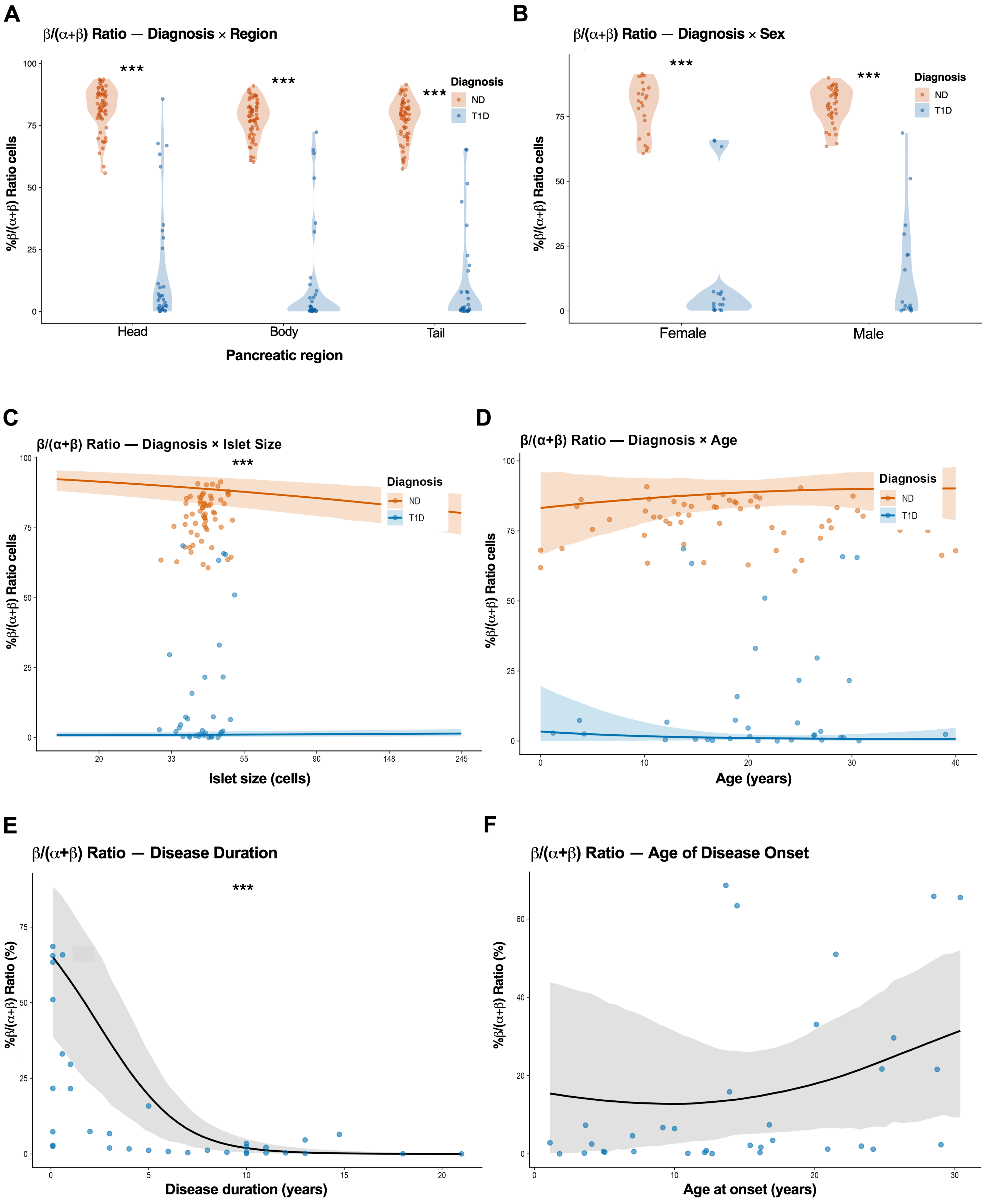
A
β/(α+β) Ratio — Diagnosis × Region
***
Diagnosis
ND
T1D
%β/(α+β) Ratio cells
Head
Body
Tail
Pancreatic region
B
β/(α+β) Ratio — Diagnosis × Sex
Female
Male
C
β/(α+β) Ratio — Diagnosis × Islet Size
Islet size (cells)
D
β/(α+β) Ratio — Diagnosis × Age
Age (years)
E
β/(α+β) Ratio — Disease Duration
%β/(α+β) Ratio (%)
Disease duration (years)
F
β/(α+β) Ratio — Age of Disease Onset
Age at onset (years)

**Supplementary Figure S10. The β/(α+β) ratio is reduced in T1D across all pancreatic regions and both sexes. (A)** β/(α+β) ratio by pancreatic region, stratified by diagnosis. ND donors maintain high ratios across all regions (Head = 0.93, Body = 0.87, Tail = 0.87), while T1D donors show near-complete β-cell loss (Head = 0.009, Body = 0.014, Tail = 0.008). Among ND donors, the pancreatic head has a significantly higher ratio than the body or tail (posterior probability = 1.0 for both comparisons), while the body and tail do not differ (posterior probability of body > tail = 0.83). **(B)** β/(α+β) ratio by sex, stratified by diagnosis. T1D donors show markedly reduced ratios relative to ND donors in both females (ND = 0.89, T1D = 0.008; posterior probability > 0.999) and males (ND = 0.89, T1D = 0.010; posterior probability > 0.999). No significant main effect of sex is observed (posterior probability of female > male = 0.53). **(C)** β/(α+β) ratio as a function of islet size, stratified by diagnosis. In ND donors, the ratio decreases with increasing islet size (slope = -0.039; posterior probability of negative slope = 1.0). In T1D donors, the slope is near zero but weakly positive (slope = 0.002; posterior probability of positive slope = 1.0). The diagnosis × islet size interaction is significant (T1D - ND slope difference = 0.041; posterior probability > 0.999). Lines represent model-predicted trends with 95% credible bands. **(D)** β/(α+β) ratio as a function of donor age, stratified by diagnosis. ND donors maintain relatively stable ratios across age (slope = 0.002/year; posterior probability of positive slope = 0.73). T1D donors cluster near zero regardless of age (slope = -0.0003/year; posterior probability of negative slope = 0.74). The diagnosis × age interaction is not significant (posterior probability of T1D slope < ND slope = 0.77). Lines represent model-predicted trends with 95% credible bands. **(E)** β/(α+β) ratio as a function of disease duration (years) in T1D donors only. The ratio declines significantly with increasing disease duration (slope = -0.062/year; posterior probability of negative slope > 0.999). This decline is consistent across all pancreatic regions (head: -0.053, body: -0.075, tail: -0.058; posterior probability of negative slope > 0.999 for all). The line represents the model-predicted trend with 95% credible band. **(F)** β/(α+β) ratio as a function of age at disease onset (years) in T1D donors only. The overall association between age at T1D onset and the ratio is not significant (slope = 0.006; posterior probability of positive slope = 0.78). The line represents the model-predicted trend with 95% credible band. All estimates are posterior medians with 95% highest posterior density intervals. Each point represents an individual donor. *Posterior probability > 0.95; ***Posterior probability > 0.999.

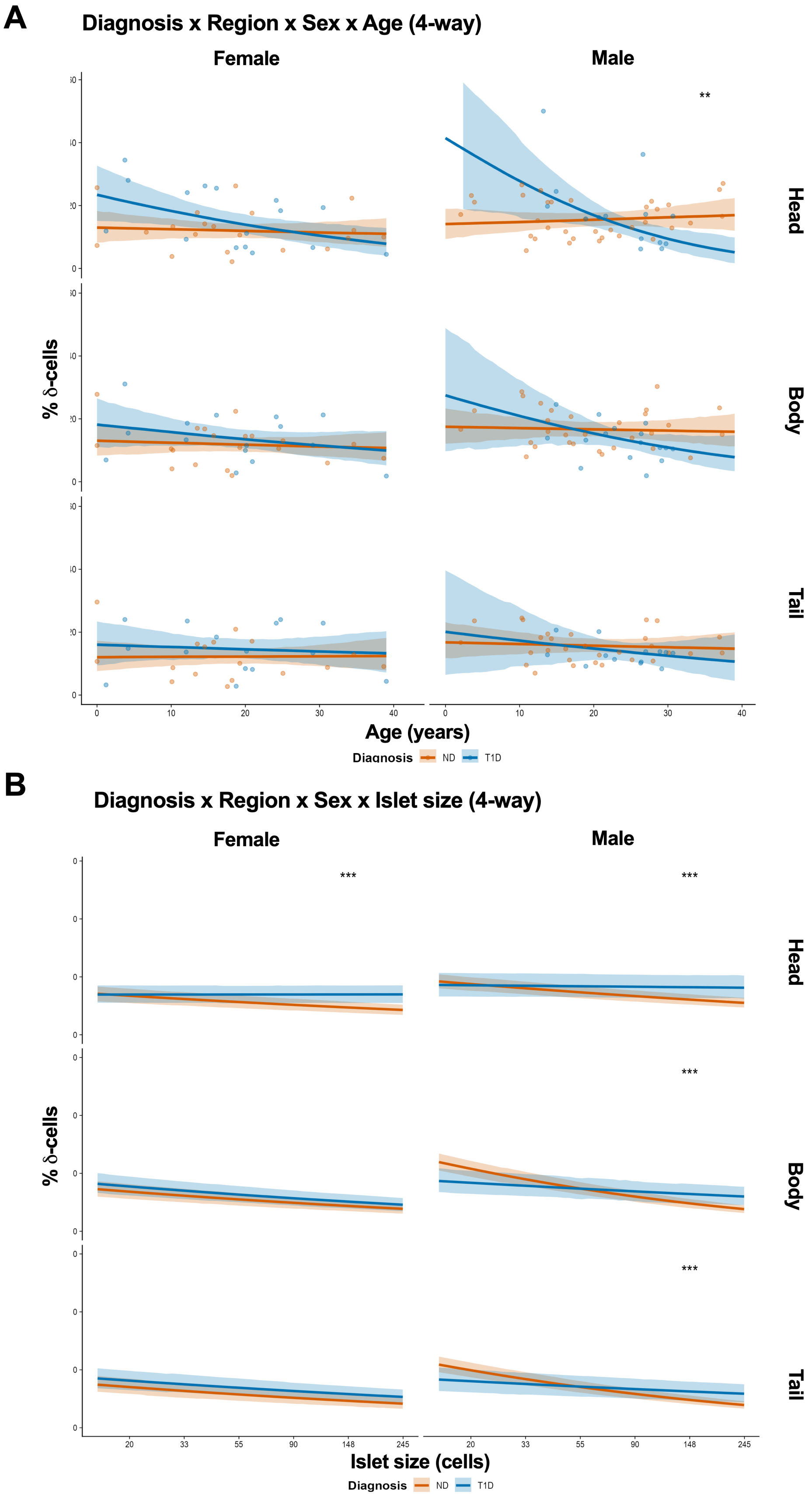
A
Diagnosis x Region x Sex x Age (4-way)
Female
Male
**
Head
Body
Tail
% δ-cells
0
10
20
30
40
Age (years)
Diagnosis
ND
T1D
B
Diagnosis x Region x Sex x Islet size (4-way)
Female
Male
***
***
***
***
Head
Body
Tail
% δ-cells
20
33
55
90
148
245
Islet size (cells)
Diagnosis
ND
T1D

**Supplementary Figure S11. Four-way interaction analyses of δ-cell proportion across diagnosis, pancreatic region, sex, age, and islet size. (A)** Percentage of δ-cells as a function of donor age (years), stratified by diagnosis (ND, orange; T1D, blue), pancreatic region (head, body, tail), and sex. Each point represents an individual donor observation. Fitted lines and 95% confidence bands are derived from a Bayesian ordered beta regression model with diagnosis × region × sex × age interactions and donor as a random effect (ND: n = 17-35; T1D: n = 13-16). In T1D males, δ-cell proportion in the head decreases significantly with age (slope = −0.0091, 95% HPD [−0.0167, −0.0023], pd = 0.998). **(B)** Percentage of δ-cells as a function of islet size (number of cells per islet), stratified by diagnosis (ND, orange; T1D, blue), pancreatic region (head, body, tail), and sex. Fitted lines and 95% confidence bands are derived from a Bayesian ordered beta regression model with diagnosis × region × sex × islet size interactions and donor as a random effect (ND: n = 17-35; T1D: n = 13-16). In both sexes, within the pancreatic head region, there is less change in δ-cell proportion with increasing islet size in T1D compared to ND (females: T1D – ND = 0.0215, 95% HPD [0.0164, 0.0260], pd > 0.999; males: T1D - ND = 0.0246, 95% HPD [0.0200, 0.0293], pd > 0.999). In males, this pattern extends to the body (T1D - ND = 0.0451, 95% HPD [0.0366, 0.0538], pd > 0.999) and tail (T1D - ND = 0.0371, 95% HPD [0.0296, 0.0440], pd > 0.999), whereas no significant T1D-ND difference is observed in females for the body or tail. All estimates are posterior medians with 95% highest posterior density intervals. **Probability of direction (pd) > 0.995, ***pd > 0.9995.

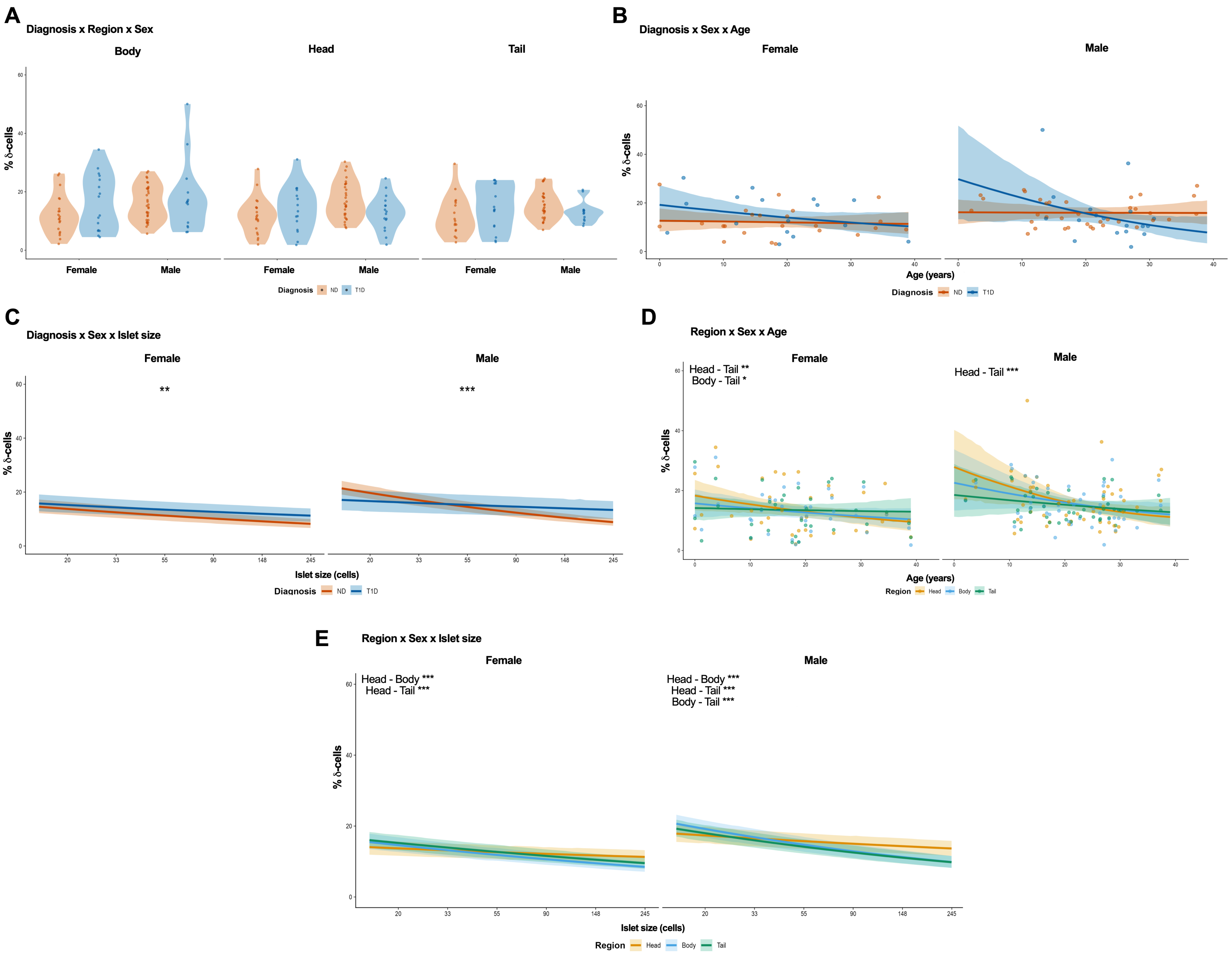


**Supplementary Figure S12. Three-way interaction analyses reveal heterogeneity in δ-cell proportion.** All panels are derived from a 4-way Bayesian ordered beta regression model with diagnosis × region × sex × age and diagnosis × region × sex × islet size interactions and donor as a random effect (ND: n = 17-35; T1D: n = 13-16). **(A)** δ-cell proportion (%) by pancreatic region (body, head, tail) and sex, stratified by diagnosis. Violin plots show the distribution of individual donor observations; points represent individual donors. No significant differences in δ-cell proportion between ND and T1D are observed in any region × sex combination. **(B)** δ-cell proportion as a function of donor age (years), stratified by diagnosis and sex and collapsed across pancreatic region. Each point represents an individual donor. Fitted lines with 95% confidence bands are shown. δ-cell proportion is stable across age in both sexes for ND and T1D donors with no significant differences between T1D and ND across age. **(C)** δ-cell proportion as a function of islet size (number of cells per islet), stratified by diagnosis and sex and collapsed across pancreatic region. Fitted lines with 95% confidence bands are shown. In both sexes, there is less change in δ-cell proportion with increasing islet size in T1D versus ND (females: T1D - ND = 0.0070, 95% HPD [0.0024, 0.0119], pd = 0.999; males: T1D - ND = 0.0356, 95% HPD [0.0300, 0.0410], pd > 0.999). **(D)** δ-cell proportion as a function of donor age, stratified by pancreatic region and sex, and collapsed across diagnosis. Each point represents an individual donor observation. Fitted lines with 95% confidence bands are shown. In females, the head shows faster decline in δ-cell proportion across age than the tail (Head - Tail = -0.0019, 95% HPD [-0.0029, -0.0009], pd > 0.999) and the body declines faster than the tail (Body - Tail = -0.0011, 95% HPD [-0.0020, -0.0001], pd = 0.985). In males, the head-tail difference is also significant (Head - Tail = -0.0027, 95% HPD [-0.0044, -0.0010], pd > 0.999). **(E)** δ-cell proportion as a function of islet size, stratified by pancreatic region and sex, collapsed across diagnosis. Fitted lines with 95% confidence bands are shown. In females, the head shows less change with increasing islet size compared to the body (Head - Body = 0.0163, 95% HPD [0.0124, 0.0201], pd > 0.999) and tail (Head - Tail = 0.0141, 95% HPD [0.0110, 0.0178], pd > 0.999). In males, all pairwise region differences are significant (Head - Body = 0.0266, 95% HPD [0.0227, 0.0307], pd > 0.999) and tail (Head - Tail = 0.0210, 95% HPD [0.0177, 0.0244], pd > 0.999), and the body shows less change than the tail (Body - Tail = -0.0056, 95% HPD [-0.0091, -0.0022], pd > 0.999). All estimates are posterior medians with 95% highest posterior density (HPD) intervals. *Probability of direction (pd) > 0.975, **pd > 0.995, ***pd > 0.999.

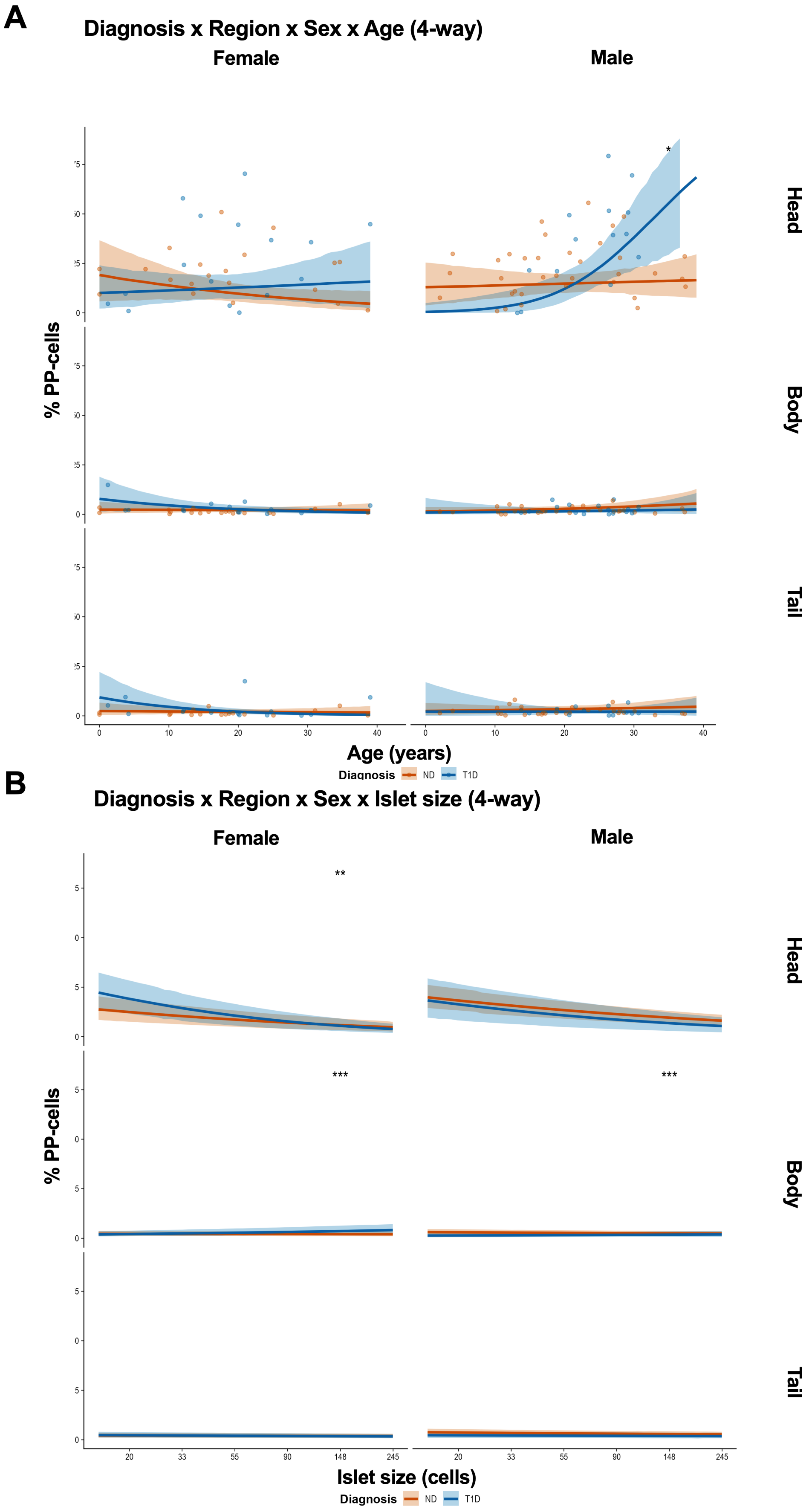
A
Diagnosis x Region x Sex x Age (4-way)
Female
Male
Head
Body
Tail
% PP-cells
*
0
10
20
30
40
Age (years)
Diagnosis
ND
T1D
B
Diagnosis x Region x Sex x Islet size (4-way)
Female
Male
**
***
***
Head
Body
Tail
% PP-cells
20
33
55
90
148
245
Islet size (cells)
Diagnosis
ND
T1D

**Supplementary Figure S13. Four-way interaction analyses of PP-cell proportion by diagnosis, pancreatic region, sex, and age and islet size.** All panels are derived from a 4-way Bayesian ordered beta regression model with diagnosis × region × sex × age and diagnosis × region × sex × islet size interactions and donor as a random effect (ND: n = 17-35; T1D: n = 13-16). **(A)** PP-cell proportion (%) as a function of donor age (years), stratified by diagnosis (ND, orange; T1D, blue), pancreatic region (head, body, tail; rows), and sex (female, male; columns). Each point represents an individual donor. Fitted lines with 95% confidence bands are shown. In the head region of T1D males, PP-cell proportion increases significantly with age (slope = 0.0162, 95% HPD [0.0063, 0.0262], pd = 0.9982), and the T1D-ND age slope difference is significant (T1D - ND = 0.015, 95% HPD [0.005, 0.027], pd = 0.995). **(B)** PP-cell proportion as a function of islet size (number of cells per islet), stratified by diagnosis, pancreatic region, and sex. Fitted lines with 95% confidence bands are shown. In the head, PP-cell proportion declines with increasing islet size in both diagnoses, with a significantly steeper decline in T1D females compared to ND females (T1D - ND = -0.042, 95% HPD [-0.073, -0.011], pd = 0.998). In the body, T1D shows a significantly more positive (or less negative) islet size slope compared to ND in both females (T1D - ND = 0.008, 95% HPD [0.003, 0.013], pd > 0.999) and males (T1D - ND = 0.005, 95% HPD [0.003, 0.008], pd > 0.999). No significant T1D - ND differences are observed in the tail. All estimates are posterior medians with 95% highest posterior density (HPD) intervals. *Probability of direction (pd) > 0.975, **pd > 0.995, ***pd > 0.999.

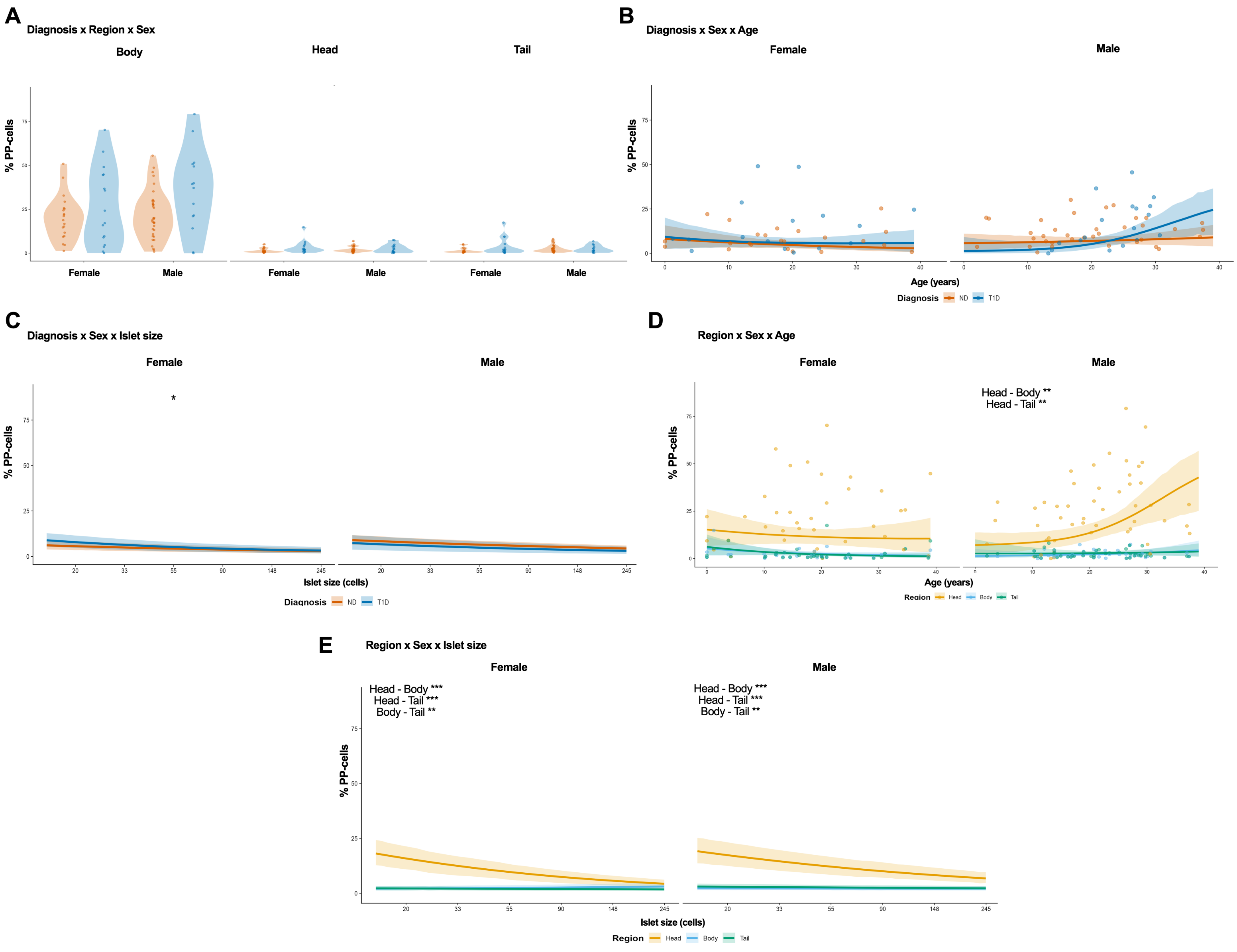


**Supplementary Figure S14. Three-way interaction analyses reveal heterogeneity in PP-cell proportion.** All panels are derived from a 4-way Bayesian ordered beta regression model with diagnosis × region × sex × age and diagnosis × region × sex × islet size interactions and donor as a random effect (ND: n = 17-35; T1D: n = 13-16). **(A)** PP-cell proportion (%) by pancreatic region (body, head, tail) and sex, stratified by diagnosis. Violin plots show the distribution of individual donor observations; points represent individual donors. PP-cell proportion is highest in the head across all diagnosis × sex combinations, with no significant diagnosis effect within any region × sex interaction. **(B)** PP-cell proportion as a function of donor age (years), stratified by diagnosis and sex, and collapsed across pancreatic region. Each point represents an individual donor. Fitted lines with 95% confidence bands are shown. PP-cell proportion is generally stable across age. In T1D males, the PP-cell proportion rises more rapidly over time as subjects age compared to ND males (slope = 0.0055, 95% HPD [0.0012, 0.0098], pd= 0.986). **(C)** PP-cell proportion as a function of islet size (number of cells per islet), stratified by diagnosis and sex, and collapsed across pancreatic region. Fitted lines with 95% confidence bands are shown. In females, the proportion of PP-cells in islets decreases to a greater degree in T1D versus ND as islet size increases (T1D - ND = -0.012, 95% HPD [-0.022, -0.002], pd = 0.991). No significant difference is observed in males (T1D - ND = 0.001, pd = 0.540). **(D)** PP-cell proportion as a function of donor age, stratified by pancreatic region and sex, and collapsed across diagnosis. Each point represents an individual donor observation. Fitted lines with 95% confidence bands are shown. In males, the PP-cell proportion rises more rapidly in the pancreatic head over time as subjects age (slope = 0.0086, 95% HPD [0.0032, 0.0146], pd = 0.997), while body and tail show no significant trends. The head-body (estimate = 0.008, 95% HPD [0.003, 0.013], pd = 0.997) and head-tail (estimate = 0.008, 95% HPD [0.004, 0.014], pd = 0.999) age slope differences are significant in males. In females, no significant age-related regional differences are observed. **(E)** PP-cell proportion as a function of islet size, stratified by pancreatic region and sex, collapsed across diagnosis. Fitted lines with 95% confidence bands are shown. In the head, PP-cell proportion declines steeply with increasing islet size, while body and tail regions show near-zero or slightly positive slopes. Females show significant region-specific differences in the change in PP-cell proportion as islet size increases: head versus body (Head - Body = −0.061, 95% HPD [-0.078, -0.044], pd > 0.999), head versus tail (Head - Tail = -0.056, 95% HPD [-0.071, -0.040], pd > 0.999), and body versus tail (Body - Tail = 0.005, 95% HPD [0.002, 0.008], pd > 0.999). Males also show differences for the head versus body (Head - Body = -0.049, 95% HPD [-0.063, -0.036], pd > 0.999), head versus tail (Head - Tail = -0.047, 95% HPD [-0.060, -0.035], pd > 0.999), and body versus tail (Body - Tail = 0.002, 95% HPD [0.001, 0.005], pd = 0.999). All estimates are posterior medians with 95% highest posterior density (HPD) intervals. *Probability of direction (pd) > 0.975, **pd > 0.995, ***pd > 0.999.

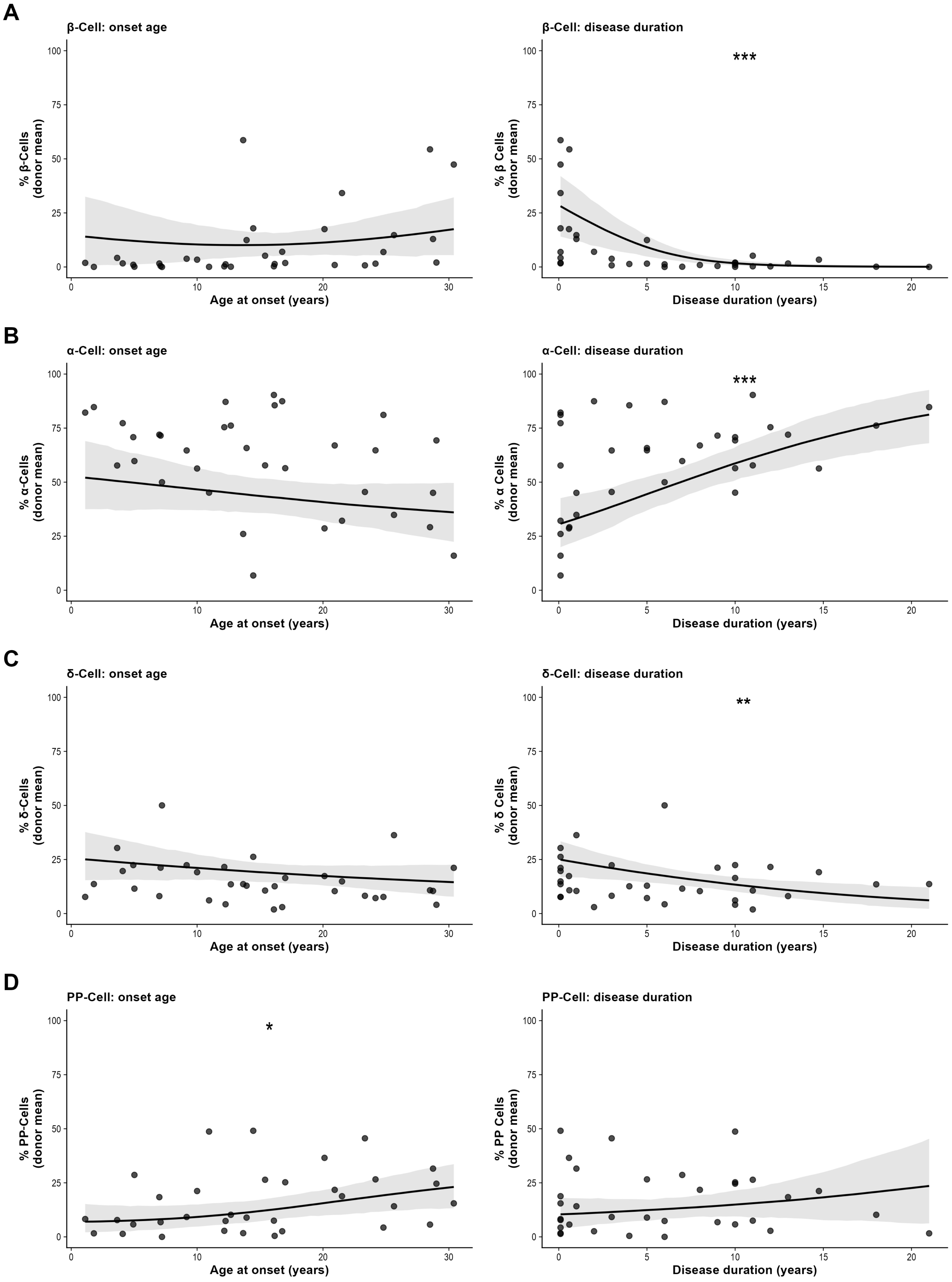

A
β-Cell: onset age
% β-Cells (donor mean)
Age at onset (years)
β-Cell: disease duration
***
% β Cells (donor mean)
Disease duration (years)
B
α-Cell: onset age
% α-Cells (donor mean)
Age at onset (years)
α-Cell: disease duration
***
% α Cells (donor mean)
Disease duration (years)
C
δ-Cell: onset age
% δ-Cells (donor mean)
Age at onset (years)
δ-Cell: disease duration
**
% δ Cells (donor mean)
Disease duration (years)
D
PP-Cell: onset age
*
% PP-Cells (donor mean)
Age at onset (years)
PP-Cell: disease duration
% PP Cells (donor mean)
Disease duration (years)

**Supplementary Figure S15. Disease duration drives progressive remodeling of islet cell composition in T1D. (A)** β-cell proportion (donor mean) as a function of age at disease onset (left) and disease duration (right) in T1D donors. β-cell proportion shows no significant association with age at onset (slope = 0.1%/year; posterior probability of positive slope = 0.589) but declines significantly with increasing disease duration (slope = -2.89%/year; posterior probability of negative slope > 0.999). **(B)** α-cell proportion (donor mean) as a function of age at onset (left) and disease duration (right) in T1D donors. α-cell proportion shows no significant association with age at onset (slope = -0.6%/year; posterior probability of negative slope = 0.89) but increases significantly with disease duration (slope = 2.89%/year; posterior probability of positive slope > 0.999). **(C)** δ-cell proportion (donor mean) as a function of age at onset (left) and disease duration (right) in T1D donors. δ-cell proportion shows no significant association with age at onset (slope = -0.36%/year; posterior probability of negative slope = 0.91) but declines modestly with disease duration (slope = -1.19%/year; posterior probability of negative slope = 0.999). **(D)** PP-cell proportion (donor mean) as a function of age at onset (left) and disease duration (right) in T1D donors. PP-cell proportion shows a significant positive association with age at onset (slope = 0.64%/year; posterior probability of positive slope = 0.98). Disease duration has no significant effect on PP-cell proportion (slope = 0.45%/year; posterior probability of positive slope = 0.81). Each point represents one T1D donor's mean cell-type proportion; lines represent model-predicted trends with 95% credible bands. All estimates are posterior medians with 95% highest posterior density intervals. *Posterior probability > 0.95; **Posterior probability > 0.99; ***Posterior probability > 0.999.

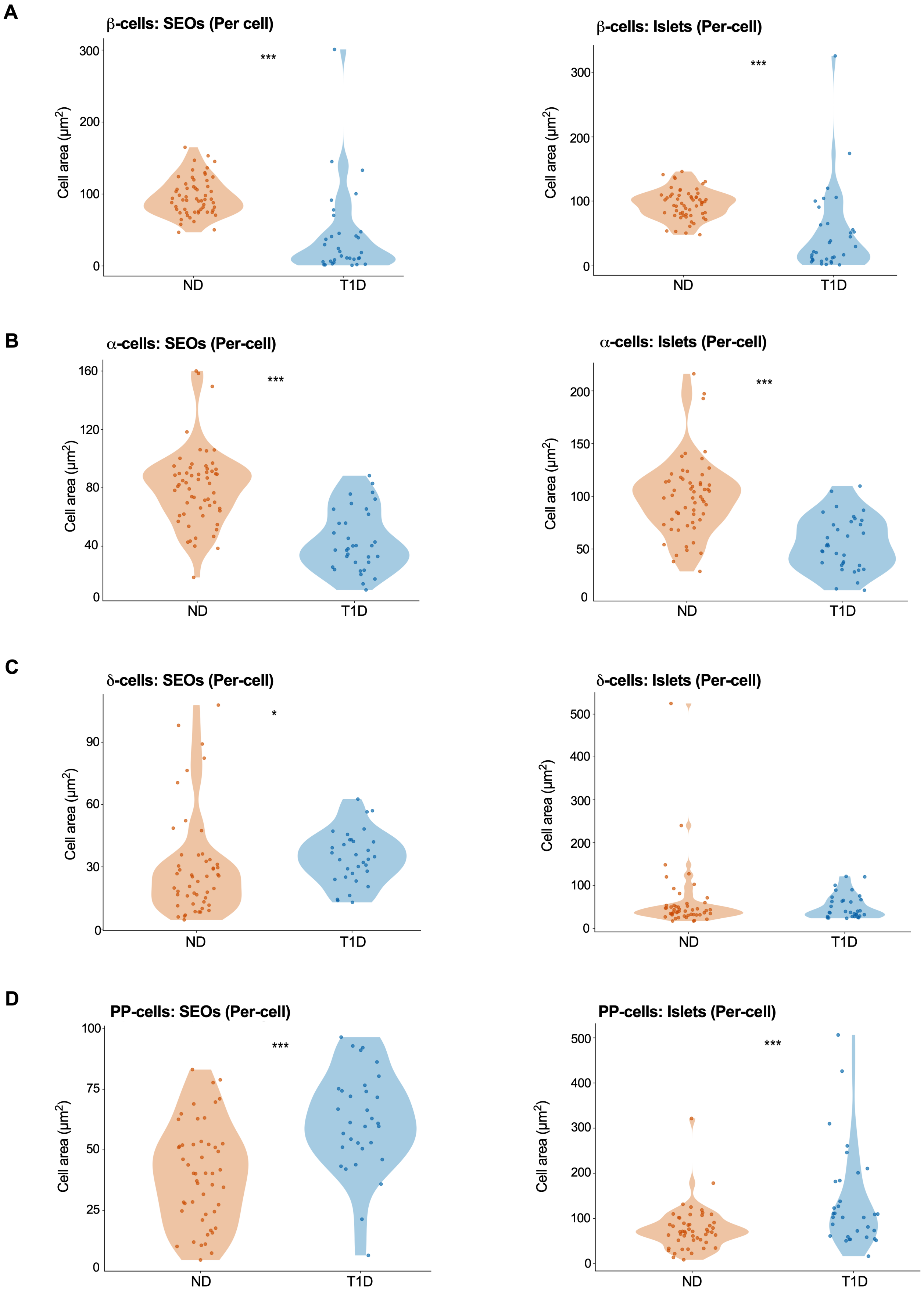

A
β-cells: SEOs (Per cell)
***
Cell area (µm2)
300
200
100
0
ND
T1D
β-cells: Islets (Per-cell)
***
Cell area (µm2)
300
200
100
0
ND
T1D
B
α-cells: SEOs (Per-cell)
***
Cell area (µm2)
160
120
80
40
0
ND
T1D
α-cells: Islets (Per-cell)
***
Cell area (µm2)
200
150
100
50
0
ND
T1D
C
δ-cells: SEOs (Per-cell)
*
Cell area (µm2)
90
60
30
0
ND
T1D
δ-cells: Islets (Per-cell)
Cell area (µm2)
500
400
300
200
100
0
ND
T1D
D
PP-cells: SEOs (Per-cell)
***
Cell area (µm2)
100
75
50
25
0
ND
T1D
PP-cells: Islets (Per-cell)
***
Cell area (µm2)
500
400
300
200
100
0
ND
T1D

**Supplementary Figure S16. T1D alters individual cell area within endocrine objects across cell types. (A)** β-cell per-cell area (μ$m^2$) within small endocrine objects (SEOs) and islets, comparing ND (orange) and T1D (blue) donors. β-cells within T1D SEOs and islets are markedly smaller than in ND (SEOs: 94.2 versus 16.6 μ$m^2$, -82.4%, $p < 0.001$; Islets: 93.7 versus 20.2 μ$m^2$, -78.4%, $p < 0.001$). **(B)** α-cell per-cell area (μ$m^2$) within SEOs and islets. α-cells within T1D SEOs and islets are significantly smaller than in ND (SEOs: 72.8 versus 38.3 μ$m^2$, -47.4%, $p < 0.001$; Islets: 92.0 versus 51.6 μ$m^2$, -43.9%, $p < 0.001$). **(C)** δ-cell per-cell area (μ$m^2$) within SEOs and islets. δ-cells within T1D SEOs are significantly larger than in ND (23.8 versus 32.7 μ$m^2$, 37.1%, $p = 0.035$); islet-level per-cell area does not differ (47.7 versus 43.5 μ$m^2$, -8.7%, $p = 0.493$). **(D)** PP-cell per-cell area (μ$m^2$) within SEOs and islets. PP-cells within T1D SEOs and islets are significantly larger compared to ND (SEOs: 26.4 versus 55.1 μ$m^2$, 109.2%, $p < 0.001$; Islets: 61.7 versus 123.5 μ$m^2$, 100.1%, $p < 0.001$). Each point represents an individual donor's mean per-cell area; violin plots show the underlying distribution. Values reported are marginal means (ND versus T1D) averaged over region and sex. *$p < 0.05$; **$p < 0.01$; ***$p < 0.001$.

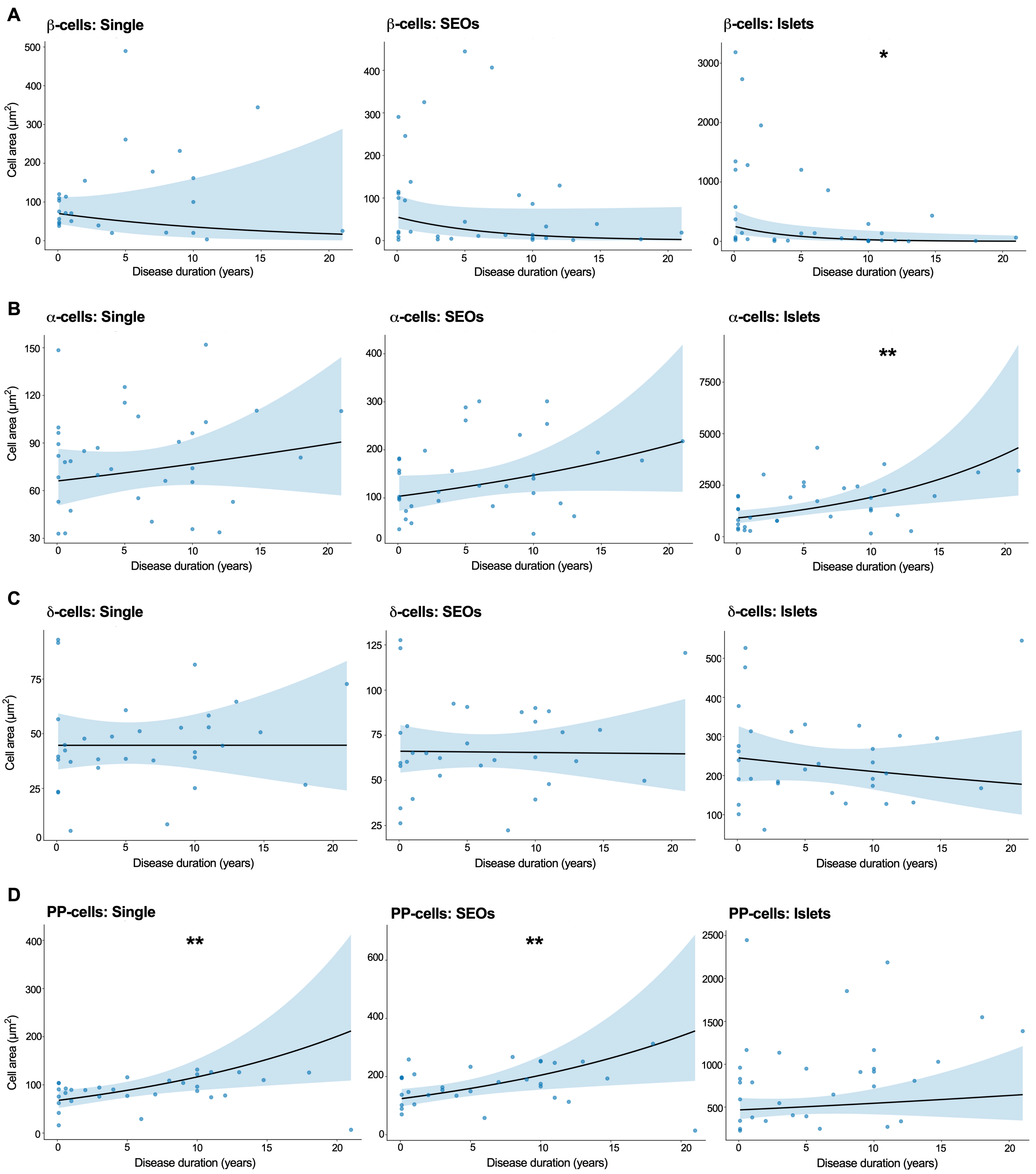

A
β-cells: Single
β-cells: SEOs
β-cells: Islets
*
B
α-cells: Single
α-cells: SEOs
α-cells: Islets
**
C
δ-cells: Single
δ-cells: SEOs
δ-cells: Islets
D
PP-cells: Single
**
PP-cells: SEOs
**
PP-cells: Islets
Cell area (μm²)
Disease duration (years)

**Supplementary Figure S17. Total endocrine cell area in T1D donors is progressively reshaped over time. (A)** β-cell area ($\mu m^2$) in single cells (Single; left), small endocrine objects (SEOs; middle), and islets (right) as a function of disease duration in T1D donors. β-cell islet area declines significantly with disease duration (slope = -0.223/year, -20.0%/year, $p = 0.011$); single-cell and SEO areas do not show significant alterations over disease duration (Single: slope = -0.068/year, -6.6%/year, $p = 0.284$; SEOs: slope = -0.148/year, -13.8%/year, $p = 0.067$). **(B)** α-cell area ($\mu m^2$) by object type as a function of disease duration. α-cell islet area increases significantly with disease duration (slope = 0.074/year, 7.7%/year, $p = 0.002$); single-cell and SEO areas do not significantly change over time (Single: slope = 0.015/year, 1.5%/year, $p = 0.339$; SEOs: slope = 0.036/year, 3.6%/year, $p = 0.100$). **(C)** δ-cell area ($\mu m^2$) by object type as a function of disease duration. δ-cell area shows no significant association with disease duration in any object type (Single: slope = 0.000/year, $p = 1.000$; SEOs: slope = -0.001/year, -0.1%/year, $p = 0.928$; Islets: slope = -0.015/year, -1.5%/year, $p = 0.398$). **(D)** PP-cell area ($\mu m^2$) by object type as a function of disease duration. PP-cell area increases significantly with disease duration in single cells and SEOs (Single: slope = 0.054/year, 5.6%/year, $p = 0.007$; SEOs: slope = 0.051/year, 5.2%/year, $p = 0.009$); islet-level area does not exhibit significant changes over disease duration (slope = 0.015/year, 1.6%/year, $p = 0.418$). Each point represents an individual T1D donor's mean cell area; lines represent model-predicted trends with 95% confidence bands. *$p < 0.05$; **$p < 0.01$; ***$p < 0.001$.

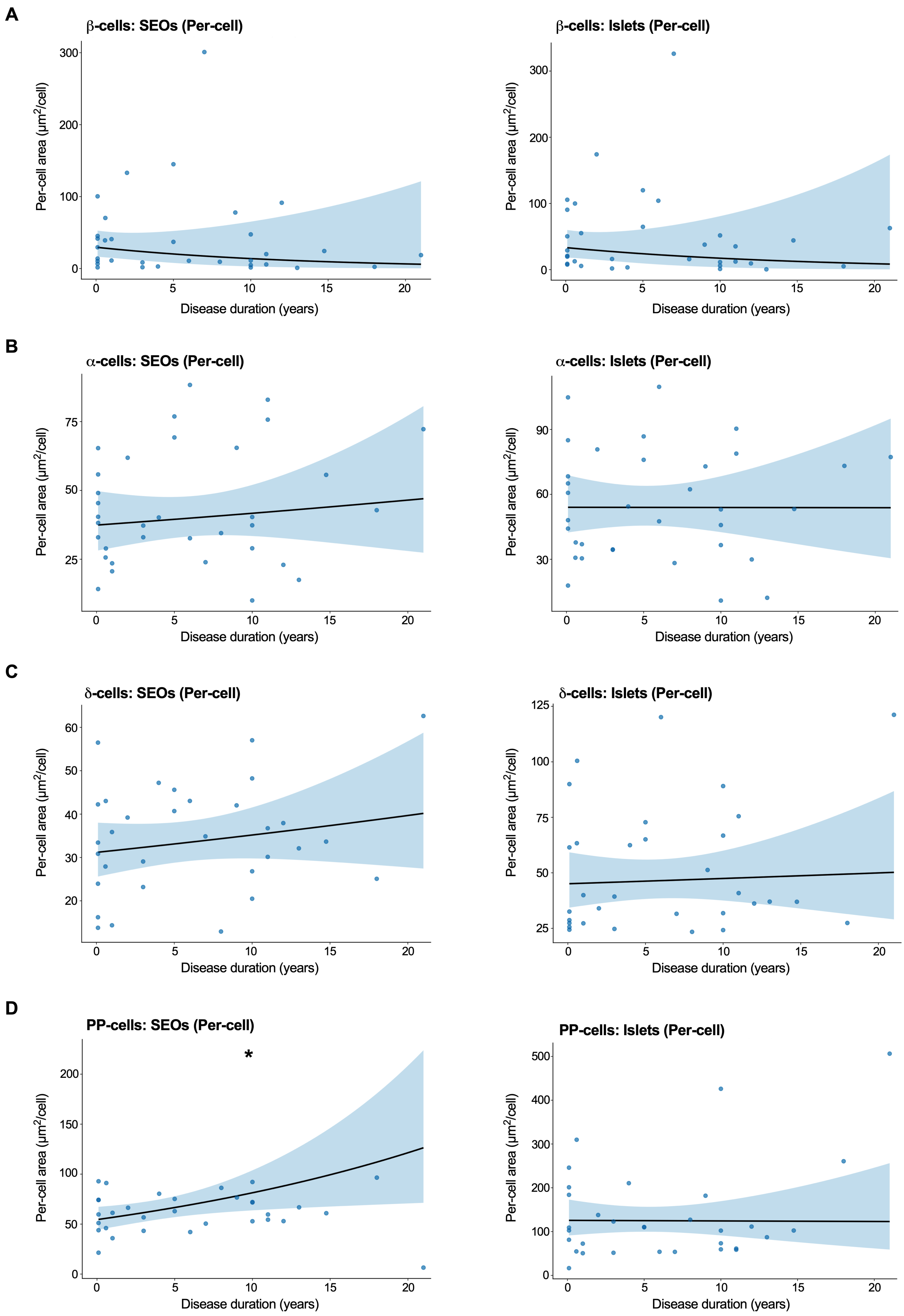
A
β-cells: SEOs (Per-cell)
β-cells: Islets (Per-cell)
B
α-cells: SEOs (Per-cell)
α-cells: Islets (Per-cell)
C
δ-cells: SEOs (Per-cell)
δ-cells: Islets (Per-cell)
D
PP-cells: SEOs (Per-cell)
*
PP-cells: Islets (Per-cell)
Per-cell area (μm²/cell)
Disease duration (years)
0
5
10
15
20

**Supplementary Figure S18. Disease duration has minimal effect on per-cell area within endocrine objects in T1D. (A)** β-cell per-cell area (μm$^2$) within small endocrine objects (SEOs; left) and islets (right) as a function of disease duration in T1D donors. β-cell per-cell area shows no significant association with disease duration in either object type (SEOs: slope = -0.076/year, -7.3%/year, p = 0.279; Islets: slope = -0.066/year, -6.3%/year, p = 0.361). **(B)** α-cell per-cell area (μm$^2$) within SEOs and islets as a function of disease duration. α-cell per-cell area shows no significant association with disease duration in either object type (SEOs: slope = 0.011/year, 1.1%/year, p = 0.538; Islets: slope = -0.0002/year, -0.02%/year, p = 0.991). **(C)** δ-cell per-cell area (μm$^2$) within SEOs and islets as a function of disease duration. δ-cell per-cell area shows no significant association with disease duration in either object type (SEOs: slope = 0.012/year, 1.2%/year, p = 0.317; Islets: slope = 0.005/year, 0.5%/year, p = 0.770). **(D)** PP-cell per-cell area (μm$^2$) within SEOs and islets as a function of disease duration. PP-cell per-cell area within SEOs shows a significant positive association with disease duration (slope = 0.040/year, 4.1%/year, p = 0.017); no significant association is observed within islets (slope = -0.001/year, -0.1%/year, p = 0.965). Each point represents an individual T1D donor's mean per-cell area; lines represent model-predicted trends with 95% confidence bands. Slopes are reported on the log scale; *p < 0.05; **p < 0.01; ***p < 0.001.

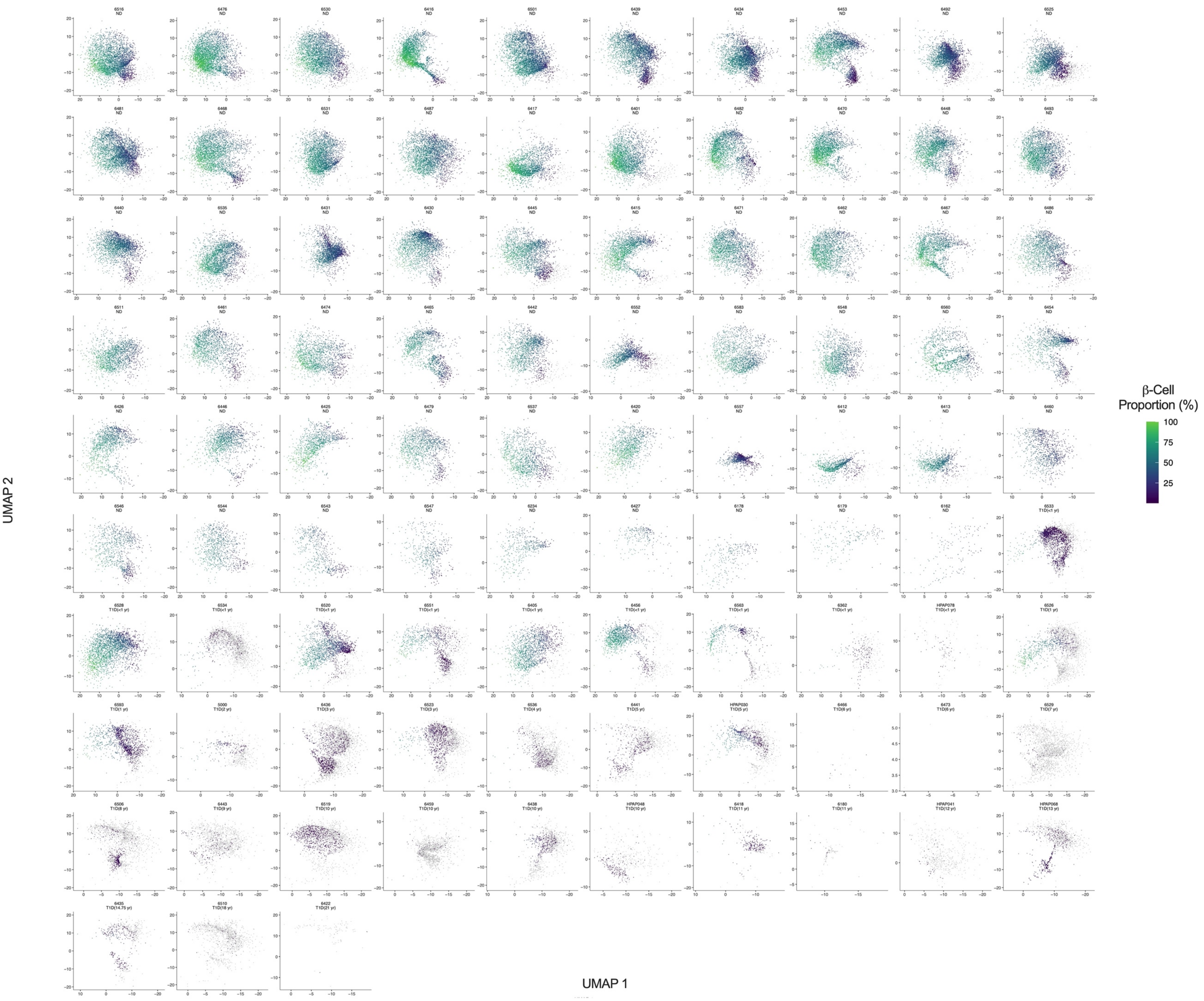


**Supplementary Figure S19. UMAP visualization of islet β-cell composition per donor.** UMAP visualization of islet β-cell composition (expressed as a percentage of the total cells within islets) from islets of individual donors, where each dot represents an islet, and the x and y axes present the top two reduced dimensions. Each UMAP visualization is labeled by donor ID, diagnosis (ND, T1D) and disease duration, as defined by the number of years following initial T1D diagnosis.

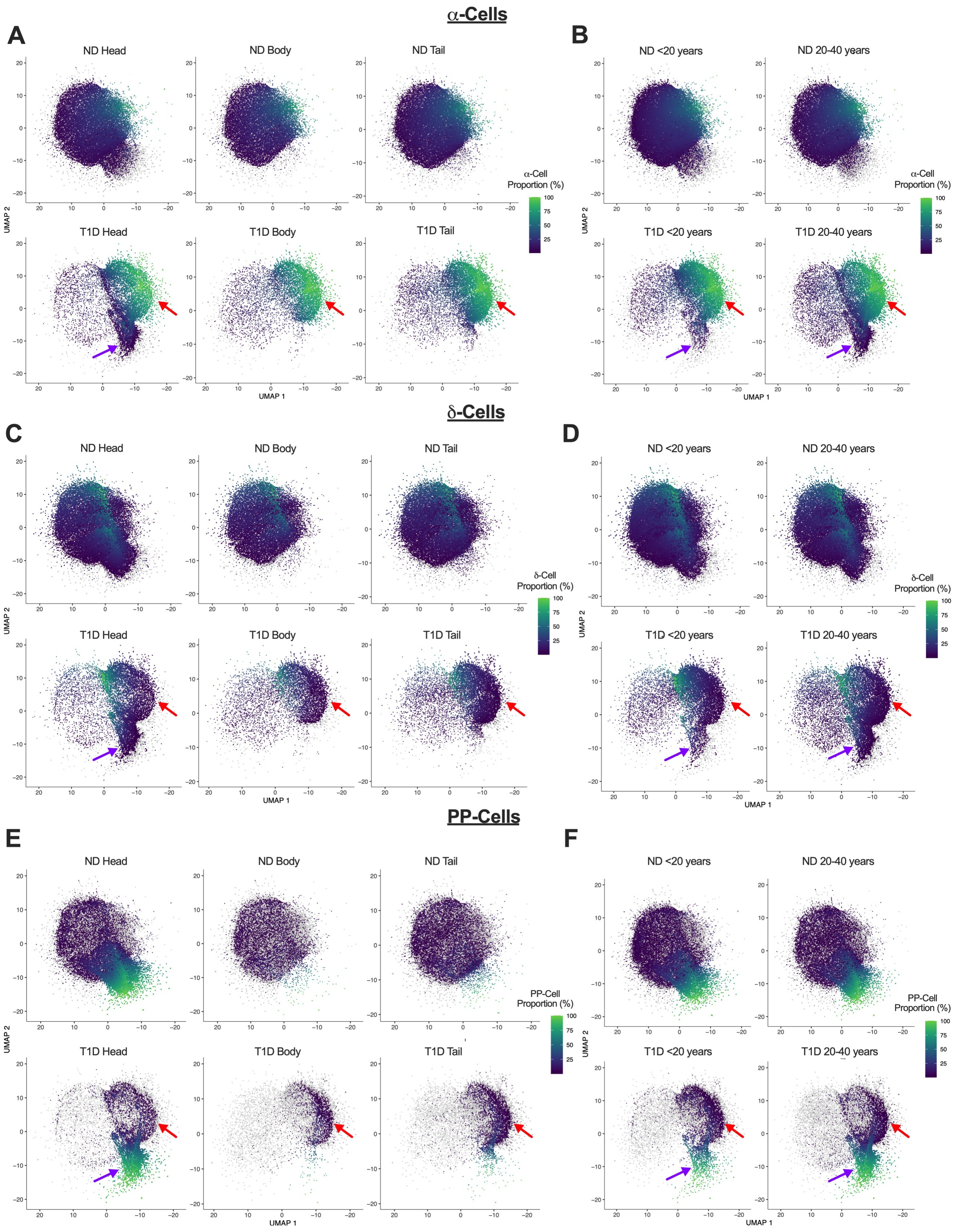
α-Cells
A
ND Head
ND Body
ND Tail
T1D Head
T1D Body
T1D Tail
α-Cell Proportion (%)
100
75
50
25
UMAP 2
UMAP 1
B
ND <20 years
ND 20-40 years
T1D <20 years
T1D 20-40 years
α-Cell Proportion (%)
δ-Cells
C
ND Head
ND Body
ND Tail
T1D Head
T1D Body
T1D Tail
δ-Cell Proportion (%)
D
ND <20 years
ND 20-40 years
T1D <20 years
T1D 20-40 years
δ-Cell Proportion (%)
PP-Cells
E
ND Head
ND Body
ND Tail
T1D Head
T1D Body
T1D Tail
PP-Cell Proportion (%)
F
ND <20 years
ND 20-40 years
T1D <20 years
T1D 20-40 years
PP-Cell Proportion (%)

**Supplementary Figure S20. UMAP visualizations of islet α-cell, δ-cell, and PP-cell populations split by diagnosis, pancreatic region, and age groups.** UMAP visualization of islet cell types from ND and T1D donor islets, where each dot represents an islet, and the x and y axes present the top two reduced dimensions. Islets are colored according to the respective proportions of α-cells, δ-cells, and PP-cells within islets. **(A-B)** UMAP visualization of islet α-cell composition (expressed as a proportion of the total cells within islets), split by diagnosis (ND versus T1D) and pancreatic region (head, body, tail) **(A)**; or diagnosis and age group (<20 years versus 20-40 years) **(B)**. **(C-D)** UMAP visualization of islet δ-cell composition (expressed as a proportion of the total cells within islets), split by diagnosis and pancreatic region **(C)**; or diagnosis and age group **(D)**. **(E-F)** UMAP visualization of islet PP-cell composition (expressed as a proportion of the total cells within islets) split by diagnosis and pancreatic region **(E)**; or diagnosis and age group **(F)**. In the respective UMAP visualizations, purple arrows indicate a T1D donor-specific islet cluster mainly present in both the pancreatic head as well as in both <20 years and 20-40-years age groups. Red arrows highlight a T1D donor-specific islet cluster present in all three pancreatic regions and both age groups.

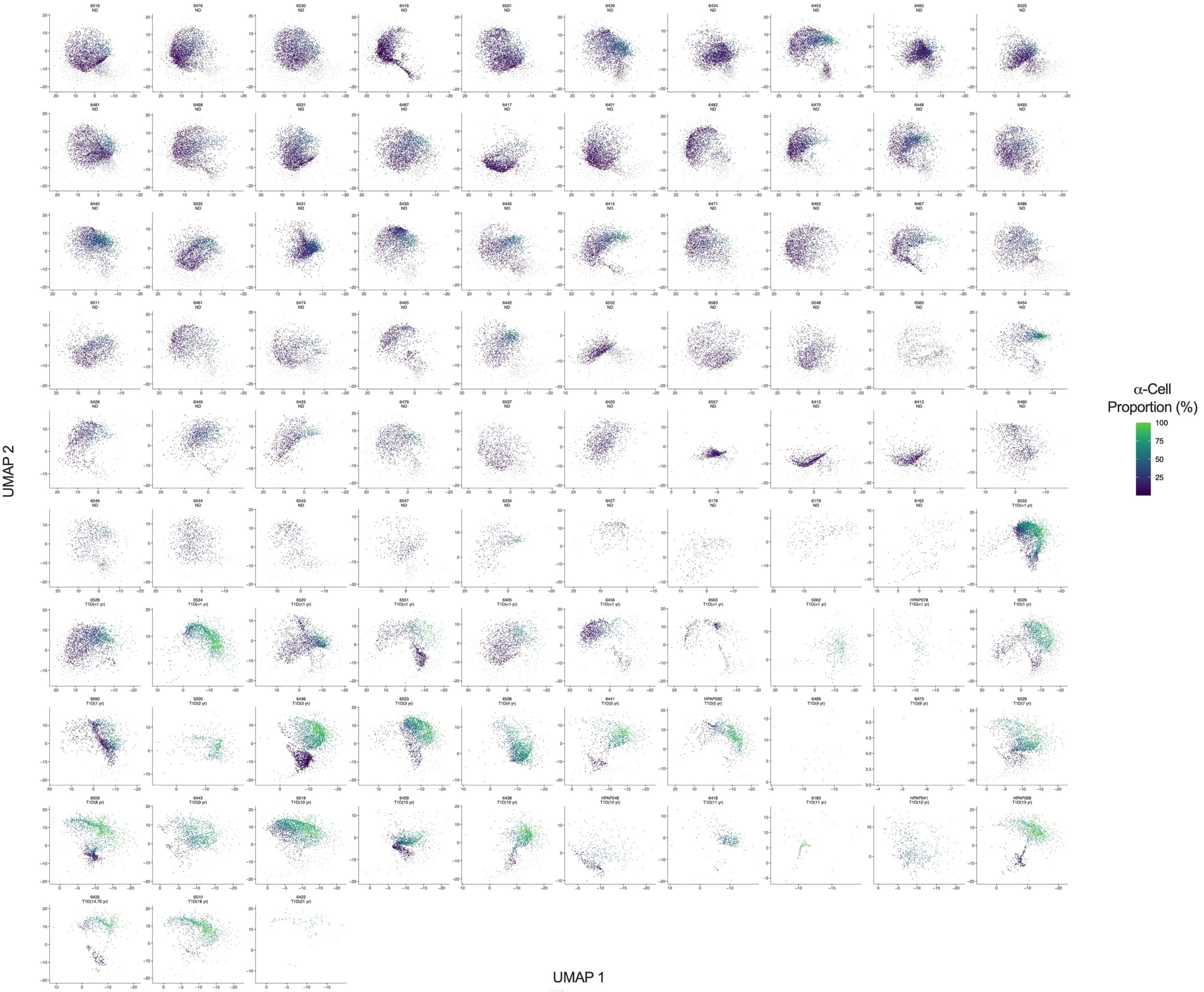


**Supplementary Figure S21. UMAP visualization of islet α-cell composition per donor.** UMAP visualization of islet α-cell composition (expressed as a proportion of the total cells within islets) from islets of individual donors, where each dot represents an islet, and the x and y axes present the top two reduced dimensions. Each UMAP visualization is labeled by donor ID, diagnosis (ND versus T1D) and disease duration, as defined by the number of years following initial T1D diagnosis.

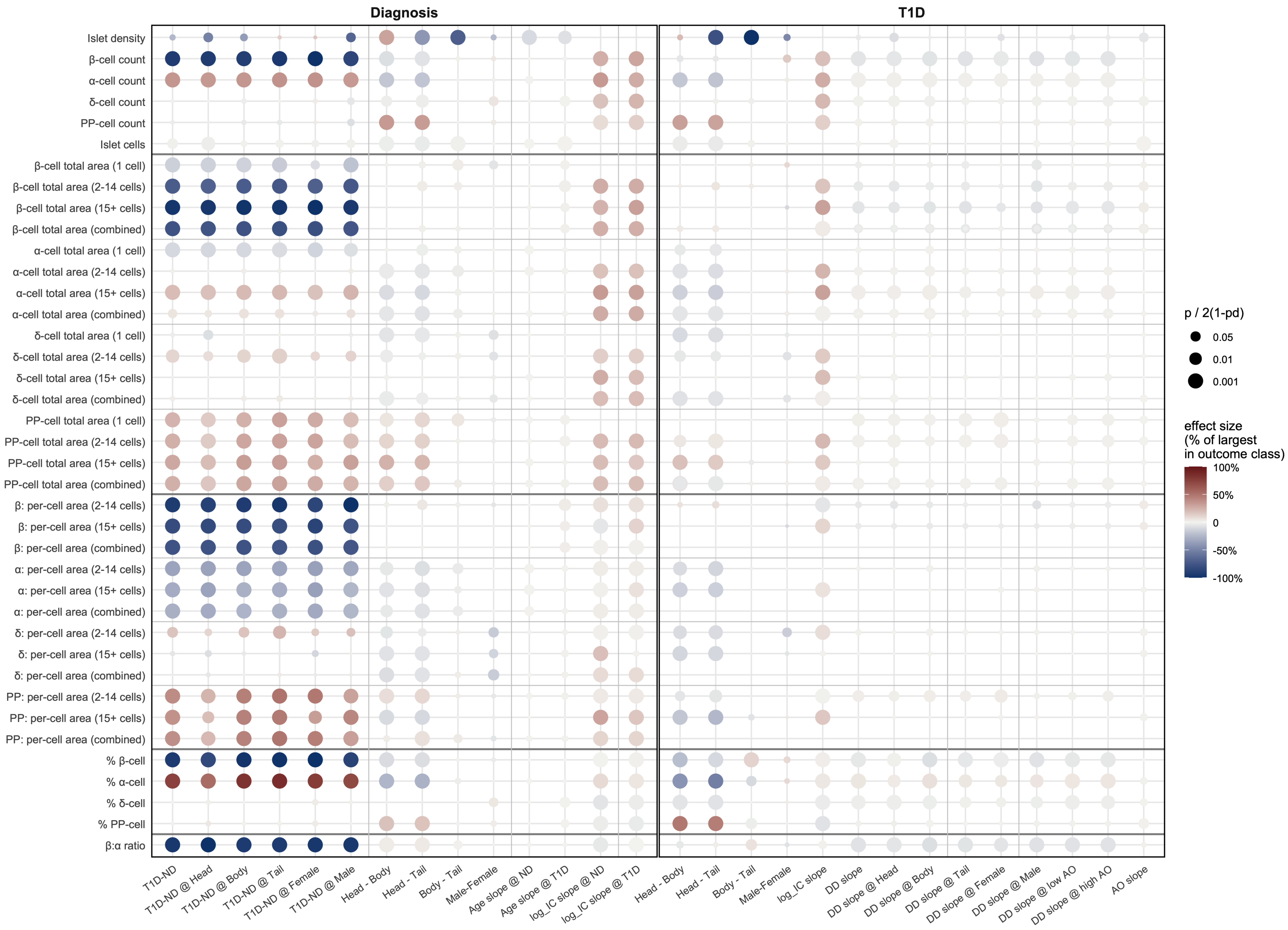


**Supplementary Figure S22. Comprehensive results summary.** Each dot represents p-value (by size) along with magnitude and direction of effect (by color) across the entire study. Reference sizes at p = 0.05, p = 0.01, and p = 0.001 are shown in the size legend.
